# New Microviridae isolated from Sulfitobacter reveals two cosmopolitan subfamilies of ssDNA phages infecting marine and terrestrial Alphaproteobacteria

**DOI:** 10.1101/2022.03.08.483405

**Authors:** Falk Zucker, Vera Bischoff, Eric Olo Ndela, Benedikt Heyerhoff, Anja Poehlein, Heike M. Freese, Simon Roux, Meinhard Simon, Francois Enault, Cristina Moraru

## Abstract

The *Microviridae* family represents one of the major clades of ssDNA phages. Their cultivated members are lytic and infect *Proteobacteria, Bacteroidetes*, and *Chlamydiae*. Prophages have been predicted in genomes from *Bacteroidales, Hyphomicrobiales*, and *Enterobacteraceae* and cluster within the “Alpavirinae”, “Amoyvirinae” and *Gokushovirinae*. We have isolated “Ascunsovirus oldenburgi” ICBM5, a novel phage distantly related to known *Microviridae*. It infects *Sulfitobacter dubius* SH24-1b and uses both a lytic and a carrier-state life strategy. Using ICBM5 proteins as a query, we uncovered in publicly available resources 65 new microviridae prophages and episomes in bacterial genomes and retrieved 47 environmental viral genomes (EVGs) from various viromes. Genome clustering based on protein content and phylogenetic analysis showed that ICBM5, together with *Rhizobium* phages, new prophages, episomes, and EVGs cluster within two new phylogenetic clades, here tentatively assigned the rank of subfamily and named “Tainavirinae” and “Occultatumvirinae”. They both infect *Rhodobacterales*. Occultatumviruses also infect *Hyphomicrobiales*, including nitrogen-fixing endosymbionts from cosmopolitan legumes. A biogeographical assessment showed that tainaviruses and occultatumviruses are spread worldwide, in terrestrial and marine environments. The new phage isolated here shed light onto new and diverse branches of the *Microviridae* tree, suggesting that much of the ssDNA phage diversity remains in the dark.

## Introduction

Viruses infecting bacteria and archaea are highly abundant in the ocean and outnumber their hosts by an order of magnitude (Bergh et al. 1989; Wommack and Colwell 2000). Through microbial cell lysis, modification of the host metabolism, and horizontal gene transfer, viruses emerge as major drivers of marine biogeochemical cycles (Suttle 2005; Roux et al. 2016; Touchon et al. 2016).

Single-stranded DNA (ssDNA) phages have been discovered quite early in the history of virology (Sertic and Boulgakov 1935; Loeb 1960) and have been extensively used as model systems and molecular biology tools (Sanger et al. 1977)(reviewed in (Székely and Breitbart 2016)). Nevertheless, most of the phages known to date, either cultivated or environmental, have double-stranded DNA (dsDNA) genomes.

A recent overhaul of the viral taxonomy (Koonin et al. 2020) has placed most ssDNA viruses, including filamentous and icosahedral phages, in the realm of *Monodnaviria*. But not all ssDNA phages are classified within the *Monodnaviria*. The *Obscuriviridae* family, with representatives infecting marine *Cellulophaga* species, is as yet unassigned to any kingdom (Bartlau et al. 2021). Furthermore, the *Cellulophaga* phage phi48:2 (Holmfeldt et al. 2013) has a protein content completely different from any of the above-mentioned phages and remains so far unclassified (Bartlau et al. 2021). Within *Monodnaviria*, filamentous phages are part of the *Loebvirae* kingdom. They have long, filamentous capsids, a circular ssDNA genome of 4.5 to 15 kb, and a chronic life cycle (Rakonjac et al. 2011; Hay and Lithgow 2019). Inovirus-like prophages have been predicted bioinformatically in a broad range of bacterial and archaeal phyla (Roux et al. 2019).

*Microviridae* is the other phage family in *Monodnaviria*, and it was recently placed in the Sangervirae kingdom. This family is very common and diverse. Its members are found in most ecosystems and are known to infect very different bacterial hosts. Its phages have small icosahedral capsids (^~^ 30 nm in diameter), a circular ssDNA of 4.4 to 6.3 kb, and all cultivated representatives are strictly lytic (Doore and Fane 2016). Viruses in the *Microviridae* family are further classified according to their genome composition and capsid structure into two ICTV-recognized subfamilies, the *Bullavirinae* and the *Gokushovirinae*. The *Bullavirinae* (former *Microvirinae*) infect enterobacteria. Gokushoviruses infect obligate parasitic bacteria, such as *Spiroplasma, Chlamydia*, and *Bdellovibrio* (Brentlinger et al. 2002; Chipman et al. 1998; Everson et al. 2002). A distinguishing feature of the gokushoviruses is the mushroom-like protrusion within their major capsid protein (MCP) (Chipman et al. 1998).

Due to their small and circular genomes, hundreds of *Microviridae-complete* genomes were assembled in metagenomic studies from diverse environments: in marine habitats (Székely and Breitbart 2016; Angly et al. 2006; Tucker et al. 2011; Labonté and Suttle 2013), freshwater habitats (Roux et al. 2012; López-Bueno et al. 2009), human gut or feces (Roux et al. 2012), stromatolites (Desnues et al. 2008), dragonflies (Rosario et al. 2012), sewage and sediments (Hopkins et al. 2014; Quaiser et al. 2015). These environmental microviruses greatly improve our understanding of the diversity of this phage family. First, the well-studied PhiX-like phages are rare in nature, and only a few genomes were detected to form a group, named pequeñoviruses, related to the *Bullavirinae* (Bryson et al. 2015; Doore and Fane 2016). Second, about half of the new genomes were affiliated to the other known subfamily, the *Gokushovirinae*. Finally, potential new subfamilies were also identified in several studies, for example, “Alpavirinae”, “Pichovirinae”, “Aravirinae” and “Stokavirinae” (Krupovic and Forterre 2011; Roux et al. 2012; Quaiser et al. 2015).

Recently cultivated ssDNA phages, infecting the marine bacteria *Citromicrobium bathyomarinum* <u>RCC</u>1878, a *Sphingomonadaceae*, and *Ruegeria pomeroyi* DSS-3, a *Rhodobacteraceae*, reveal further diversity of the *Microviridae* (Zheng et al. 2018; Zhan and Chen 2019a). The *Citromicrobium* phage was suggested to belong to a new subfamily in the *Microviridae* – the “Amoyvirinae” (Zheng et al. 2018), whereas the two ssDNA *Ruegeria* phages vB_RpoMi-Mini and vB_RpoMi-V15 are considered as unclassified *Microviridae* (Zhan and Chen 2019a).

For a long time, *Microviridae* were believed to be strictly lytic and incapable of lysogeny (Fane et al. 2006), until prophages were predicted bioinformatically in the genomes of *Bacteroidetes* (Krupovic and Forterre 2011). Further studies predicted *Microviridae*-like prophages in other *Bacteroidetes (Roux et al. 2012; Holmfeldt et al. 2013; Quaiser et al. 2015)* and in a *Caenibius tardaugens* strain, an alphaproteobacterium (Zheng et al. 2018). Even more, the ability of such a gokushovirus prophage to form viable virus particles was recently demonstrated by leveraging molecular cloning techniques (Kirchberger and Ochman 2020). Lacking integrases, these phages integrate into the host genome using its chromosome dimer resolution system (Krupovic and Forterre 2011; Kirchberger and Ochman 2020).

In our laboratory, we have established a large collection of marine phage isolates from the North Sea, infecting environmentally relevant heterotrophic bacteria belonging to the Roseobacter group. Through this work, we have screened for the presence of ssDNA phages, based on the hypothesis that the use of new hosts for phage isolation could reveal new *Microviridae* diversity. One of our phage isolates, infecting *Sulfitobacter* sp. SH24-1b and named ICBM5 was indeed a lytic, icosahedral ssDNA phage, distantly related from known *Microviridae*. Having discovered this novel phage, we wanted to know further how does it and its relatives compare with other *Microviridae* in terms of lifestyle, phylogenetic classification, integration in bacterial genomes, infected hosts, and spread in the environment.

## Materials and methods

### Isolation of the phage ICBM5

Phage ICBM5 was isolated from the coastal North Sea using a phage enrichment procedure, followed by plaque picking and purification. For this purpose, surface seawater was collected in June 2015 from the shoreline near Neuharlingersiel (53°42’09.8”N 7°41’58.9”E) during high tide, transported to the lab on ice and then 0.2 μm filtered (Rotilabo-syringe filters, Carl Roth, Germany). A phage enrichment was setup by mixing 9 parts of freshly filtered seawater with 1 part of 10x Marine Broth (MB) (see SI file 1 text) and adding an inoculum of exponentially growing *Sulfitobacter* sp. SH24-1b (Hahnke et al. 2013). After overnight incubation at 20°C and 100 rpm, cells and debris were removed from the enrichment by centrifugation (15 min, 4000 x g, 20 °C) and 0.2 μm filtration of the supernatant. To test for the presence of phages, the filtrate was spotted on a lawn of *Sulfitobacter* sp. SH24-1b. The clearing zone was then picked and single plaques were obtained and purified using the plaque assay (see SI file 1 text). Phage ICBM5 was stored either as phage lysate at +4°C or as glycerol stock of free phages or infected cells at −80°C (see SI file 1 text).

### Host range of the phage ICBM5

To determine the host range of ICBM5, a spot assay was performed with 94 bacterial strains (SI file 1 text and Table S2) covering the phylogenetic diversity of *Rhodobacteraceae*, at three different temperatures (15°C, 20°C, and 28°C). With those strains for which spots were observed, a plaque assay was performed for confirmation. Further details about the host range determination are found in SI file 1 text.

### Transmission electron microscopy of phage ICBM5

Phage ICBM5 was grown on double-layer agar plates at a phage inoculum concentration which resulted in confluent plaques. The phage fraction was collected, concentrated by polyethylene glycol (PEG, Promega, Austria) precipitation, and purified by CsCl2 centrifugation. Staining was done with uranyl acetate before transmission electron microscopy (TEM). Phages negatively stained were used for capsid size measurements. See SI file 1 text for further details.

### Testing the ssDNA nature of the ICBM5 phage genome

ICBM5 lysate was collected from plates with confluent plaques and the phage fraction was concentrated using PEG and purified by CsCl centrifugation. Further, the DNA was extracted using a phenol-chloroform procedure and exposed to four different enzymes: Exonuclease VII (Thermo Fisher Scientific, USA), TURBO DNase (Thermo Fisher Scientific, USA), S1 nuclease (Thermo Fisher Scientific, USA), and the restriction enzyme Hind III (New England BioLabs, England). Exonuclease VII and S1 strictly target single-stranded DNA, while TURBO DNase digests both, single and double-stranded DNA. Hind III targets only dsDNA. The digestion results were made visible via gel electrophoresis. For details about the individual steps in this procedure, see SI file 1 text.

### Genome sequencing of phage ICBM5

ICBM5 was grown to confluent plaques on double-layer agar plates. The phage fraction was concentrated with Amicon Ultra – 15 ml 100 kDa centrifugal filters (Merck Millipore, USA) and purified by gradient ultracentrifugation in Optiprep density gradient medium. (Sigma-Aldrich, USA). A treatment with DNase to remove free, chromosomal DNA, was followed by DNA purification using the ChargeSwitch gDNA Mini Bacteria Kit (ThermoFisher Scientific, USA). The ssDNA genome of the ICBM5 phage was converted to dsDNA by using the REPLI-g Mini kit (Qiagen, Netherlands) and then sequenced with Illumina (paired-end technology 2×300 bp) technology. For details about the individual steps in this procedure, see SI file 1 text.

### Assembly and annotation of the ICBM5 phage genome

The Illumina raw reads were cleaned with BBDuk in two steps. In the first step, the adaptors were removed, using the following parameters for BBDuk: “ktrim=r k=21 mink=8 tbo tpe ftm=5 rcomp=t ordered t=8”. In the second step, any contaminating reads (from the host, or from phiX174), as well as low-quality ends, were removed, using the following parameters for BBDuk: “k=31 rcomp=t hdist=1 qtrim=rl trimq=20, maq=20 minlen=30 ordered t=8”. Afterward, the cleaned reads were assembled with Tadpole (parameters “k=50 t=8”). Both BBDuk and Tadpole are part of the BBTools package (https://jgi.doe.gov/data-and-tools/bbtools/). After assembly, direct terminal repeats were detected at the end of the contig, indicating that the contig can be circularized and that the genome is complete. For further analyses, the genome was linearized and one of the repeats was removed. Open reading frames (ORFs) were predicted using the MetaGeneAnnotator (Noguchi et al. 2008) implemented in VirClust (Moraru 2021). A first ORF annotation was done by using Domain Enhanced Lookup Time Accelerated BLAST (DELTA-BLAST) to search for homologous proteins in the NR database (http://ncbi.nlm.nih.gov/). The ICBM5 phage genome is available in the NCBI GenBank database under the following accession number: OM782324. The sequences of the complete genome and the encoded proteins can also be found at the end of the SI file 1.

### Obtaining a Sulfitobacter sp. SH24-1b strain resistant to ICBM5

ICBM5 plaques were obtained by plating on a lawn of *Sulfitobacter* sp. SH24-1b. After 48 h of incubation, several turbid plaques were picked and resuspended to 50 μl MB medium. Dilutions of 10^-2^ – 10^-5^ were then plated on MB agar and incubated at 20 °C. Single colonies were picked after 24 h of incubation and transferred to new MB agar plates, for further experiments. The presence of ICBM5 in cultures derived from the respective single colonies was tested by PCR with primers specific for ICBM5. One culture was selected further for DNA extraction and sequencing using two long-read technologies: PacBio and Nanopore (see SI file 1 text).

### One-step growth experiment with phage ICBM5

*Sulfitobacter* sp. SH24-1b was grown in MB media for three consecutive generations, to ensure consistent growth. In the last generation, when the bacteria reached an OD of 0.3, an ICBM5 phage stock, which was prepared via gradient ultracentrifugation (See SI file 1 text), was added to a final multiplicity of infection (MOI) of 6.5. In parallel, a second flask was prepared where ASW base was added instead of phage stock, which served as a negative control over the course of infection. After 20 min of incubation at 20 °C, to allow phage adsorption, free phages were removed by centrifugation. The cells were resuspended in fresh media and incubated at 20 °C and 100 rpm for 200 min. Samples were collected every 15 min for phage counts by double-agar layer assays and phageFISH. For details, see SI file 1 text.

### ICBM5-targeted direct-geneFISH

To detect intracellular phages, we have used a modified version of the direct-geneFISH protocol (Barrero-Canosa et al. 2017), which we named phage-targeted genome-FISH. This protocol was applied both on the ICBM5 infected culture and on the control, not infected culture.

#### Design and synthesis of ICBM5 specific genome probes

To target the ICBM5 phage genome, 8 dsDNA polynucleotides (see SI file 1 text, Fig. S2 and Tab. S3) were designed using geneProber (gene-prober.icbm.de/). The corresponding dsDNA molecules were synthesized by IDT (Integrated DNA Technologies, USA) and labeled with Alexa Fluro 594 (Barrero-Canosa and Moraru 2021a), see SI file 1 text for details.

#### Immobilization on solid support

The cells from the one-step growth experiment were immobilized by spotting 20 μl of fixed cell suspension on SuperFrost Plus glass slides (Electron Microscopy Sciences, USA), on which silicone isolators (Grace Bio-Lab, USA) were placed to create wells. The cells were dried at 37°C, then washed for 1 min in 0.22 μm filtered, deionized and autoclaved water, and 10-30 sec in absolute ethanol (Th. Geyer, Germany).

#### Permeabilization and RNA removal

The samples were overlaid with a solution containing 0.5 mg/ml lysozyme (Sigma Aldrich, USA), RNase Cocktail (500 Uml^-1^ RNase A, 20.000 Uml^-1^ RNase T1) (Thermo Fisher, USA), 0.05 M EDTA pH 8.0, and 0.1 M Tris-HCl pH 8.0. Then, the samples were incubated for 30 min at 37°C, followed by washing twice for 5 min in 1 x PBS, 1 min in MQ water, 10-30 sec in absolute ethanol, and finally, by air drying.

#### Denaturation and hybridization

A hybridization buffer containing 45% formamide, 5 x SSC (750 mM NaCl, 0.075 mM sodium citrate), 20% dextran sulfate, 0.1% SDS, 20 mM EDTA, 0.25 mg/ml sheared salmon sperm DNA, 0.25 mg/ml yeast RNA and 1 % blocking reagent for nucleic acids (Roche, Switzerland) was used. To this buffer, the ICBM5 genome probes were added at a final concentration of 30 pg/μl for each polynucleotide probe. The samples were covered with the probe-hybridization buffer mixture, and denatured for 40 min at 85 °C, followed by 2 h of hybridization at 46 °C. After hybridization, the samples were quickly rinsed in washing buffer I (2 x SSC, 0.1 % SDS) at room temperature, and for 30 min washed in washing buffer II (0.1 x SSC, 0.1 % SDS) at 48°C. Finally, the samples were washed for 15 min in 1 X PBS, 1 min in water, and air-dried.

#### Counterstaining and embedding for microscopy

The samples were counterstained using 5 ng ml^-1^ DAPI dissolved in SlowFade Gold (Invitrogen, USA) and covered with a #1.5 high precision coverslip (Marienfeld, Germany).

#### Microscopy

Samples generated from phageFISH were visualized using an Axio Imager .72m fluorescent microscope (Carl Zeiss, Germany), with the help of its associated software AxioVision (version 4.8.2.0) (Carl Zeiss, Microlmaging GmbH, 2006-2010). For each field of view, a set of images was created using different exposure times for the two fluorescent channels, DAPI and Alexa594. For DAPI, the following filter set was used: 365 excitation, 445/50 emission, and 395 Beam Splitter. For Alexa594, the following filter set was used: 562/40 excitation, 624/40 emission, and 593 Beam Splitter. The exposure times were 40, 80, and 150 ms for DAPI, and 80, 200, 600, 1200, 3000, and 5000 ms for Alexa594. Each exposure time was saved individually as TIFF for further image analysis.

#### Image analysis

Image analysis was performed using CellProfiler v. 3.1.9 (McQuin et al. 2018) and had two purposes: I) to quantify the fraction of infected cells and II) to quantify the number of phage genomes per cell. In the beginning, a common analysis workflow consisting of cell detection and background removal was applied. Then, the fraction of infected cells was determined by manually investigating 500 cells per time point for the presence of a phage signal. The number of phage genomes per cell was determined only for a subset of 100 phage positive cells for each time point. For this purpose, for each cell, the total phage signal intensity was determined and then divided with the signal intensity of a single phage. Further details about the image analysis can be found in SI file 1 text.

### Detection and curation of ICBM5-like regions in bacterial genomes

Proteins from phage ICBM5 were used to query the NR database from NCBI, using DELTA-BLAST, with two iterations. Proteins detected as similar were downloaded in GenBank format, imported into Geneious v 9.1.5 (http://www.geneious.com (Kearse et al. 2012)), and were identified as part of a viral or bacterial genome based on their organism name and taxonomy (see next section for the analysis of viral sequences). Bacterial strains having hits with at least two different ICBM5 phage proteins were considered to potentially harbor ICBM5-like prophages and were selected for further analysis.

To determine the prophage borders, close relatives (same species) of each bacterial strains were searched in the GenBank sequence database, in order to have very similar bacterial genomes both with and without the prophage. Similar bacterial genomes were aligned using MAUVE (Darling et al. 2004) and and prophage regions were precisely identified from these alignments. We refer to these prophages as “sure border prophages” (SBPs). For the remaining bacterial strains, referred as “unsure border prophages” (UBPs), a larger genomic region surrounding the phage-like genes was selected and prophage regions were further refined. First, proteins of these UBPs were predicted using MetaGeneAnnotator and clustered with proteins of ICBM5, of other publicly available ssDNA phages and also of the sure border prophages. To this end, an all against all BLASTp (e-value < 1e-5 and bitscore > 50) was performed and proteins were clustered using the mcl program, with the parameters “-I 2 --abc”. Using the defined protein clusters, the following steps were defined to better identify the borders of UBPs: 1) the UBP genes were judged as phage genes if they were annotated as major capsid protein, replication initiation protein, lysis, or pilot protein, or if they grouped in protein clusters with proteins from the SBPs or the reference ssDNA phages; 2) if, on a UBP, genes encoding hypothetical proteins were located between the previously determined phage genes, they were kept and labeled as phage genes; 3) if, on a UPB, genes not classified in any of the above categories were located at the periphery of a region encoding phage genes, they were considered of bacterial origin and the UBPs were shortened correspondingly. Those genes that occurred in opposite direction on the contig, in comparison to the rest of the genes, were removed from the dataset as well as those genes at the border that were unique or could not be identified. If a ribosomal binding site could be determined for the first gene, this has been used as the start point for the prophage region. Otherwise, the start of the prophage region was determined at the start codon of the first gene.

### Detection of ICBM5-like regions in viral metagenomes

A set of 2 944 publicly available viromes (see SI file 2) were downloaded as raw reads that were first cleaned by removing potential adapters with cutadapt (Martin M. 2011) and trimmed using Trimmomatic (Bolger et al. 2014). Each dataset was then individually assembled into contigs using Newbler 2.6 (454; Life Sciences, Branford, CT, USA), IDBA_UD (Peng et al. 2012) or megahit (Li et al. 2015) with default parameters, depending on the sequencing technology. Details about these viromes (database source, sequencing technology, publication, etc) can be found in SI file 2. In order to detect *Microviridae* similar to ICBM5, circular contigs between 3 and 8 Kb were extracted and their proteins were predicted using prodigal (Hyatt et al. 2012). Using MMseqs (Steinegger and Soedin 2017) with a threshold of 50 on the bit-score, these proteins were compared to the MCP sequence of ICBM5 and 16 circular contigs were found to have a protein similar to ICBM5 MCP (Steinegger et al. 2019). In addition to these contigs found in the above viromes, we also added the environmental viral genomes (EVGs) retrieved from NCBI using DELTA-BLAST, during our search for ICBM5-like regions in the NR database described in the previous section.

### Clustering *Microviridae* proteins and genomes

The dataset used to classify phage ICBM5 was assembled by combining reference *Microviridae* genomes from publicly available sequence databases (NCBI), the ICBM5 phage genome, the newly detected ICBM5-like regions in bacterial genomes, as well as ICBM5 related EVGs. Here, we define as reference those microvirus genomes, be it from phage isolates or EVGs, that have been previously assigned to different *Microviridae* subfamilies. In addition, we have included in the analysis phage genomes from the *Obscuriviridae* family (Bartlau et al. 2021).

The genomes from all ssDNA phages in the above dataset were clustered hierarchically with VirClust, based on their protein super-supercluster content (Moraru 2021). Shortly, the genomes were translated into proteins, and the proteins were subjected to three clustering steps. In the first step, which grouped the proteins into protein clusters (PCs), the following parameters were used: BLASTp based similarity (evalue > 0.00001, bitscore >= 30 and coverage > 70) and clustering based on log evalues. In the second step, which grouped PCs into protein super-clusters (PSCs), the following parameters were used: Hiden Markow-Model (HMM) based similarity (probability >= 90, coverage >= 60, no threshold on alignment legth) and clustering based on log evalues. In the third step, which grouped PSCs into protein super-superclusters (PSSCs), the following parameters were used: HMM based similarity (probability >= 90, no threshold on coverage and alignment length). We refer from here on to PSSCs as “protein clusters”. Afterward, the phage genomes were clustered hierarchically with a clustering distance of 0.9.

To annotate the resulting protein clusters, we compared the individual protein sequences to various databases using VirClust. The NR database from NCBI was queried using BLASTp 2.6.0+ and the InterPro database v 66.0 (Finn et al. 2017) was queried using Inter-ProScan 5.27-66.0 tool (Jones et al. 2014). The prokaryotic Viruses Orthologous Groups (pVOGs) database (Grazziotin et al. 2017), the Virus Orthologous Group (VOGDB) database (Kiening et al. 2019), and the Prokaryotic virus Remote Homologous Groups (PHROGS) database were searched using hhsearch (Steinegger et al. 2019). Finally, the efam database (Zayed et al. 2021) was searched using hmmscan (Eddy 2011). At the end, the annotations of each protein were manually curated and compared with the annotations of the other proteins in the same protein cluster. The annotation of each protein cluster represented a consensus of the annotations of all proteins in the cluster (see SI file 3).

### Phylogenetic analysis of the ssDNA phages based on their MCP and Rep proteins

The recognizable MCP and Rep proteins from all phages in the dataset were aligned using Clustal Omega (Sievers et al. 2011) and then concatenated. A maximum-likelihood phylogeny was computed based with RAxML v8.2.12 (Stamatakis 2014), using an automatic determination of the best protein model (option -m PROTGAMMAAUTO) and 100 bootstrap replicates. The resulting phylogenetic tree was further visualized and refined with iTOL (Letunic and Bork 2021). This analysis did not include *Obscuriviridae* phages, because they have no recognizable MCP or Rep genes.

### Phylogenetic analysis of all host 16S rRNA genes and species assignment for *Sulfitobacter* sp. SH24-1b

A neighbor-joining tree of the 16S rRNA gene sequences from all phage hosts for which we could find the 16S rRNA gene (see SI file 4 Table 3) in this study was constructed with the ARB software package (Ludwig et al. 2004). Tree calculation was performed using the reference data set SSU Ref NR 111, with Jukes-Cantor correction, termini filter, and 1000 bootstrap replicates. Members of the genus *Acidobacterium* served as an outgroup. For species assignment of *Sulfitobacter* sp. SH24-1b, the ANI value between SH24-1b and the Sulfitobacter dubius type strain DSM 16472T was calculated with FastANI (Jain et al. 2018) and the dDDH value with GGDC (applying formula 2) (Meier-Kolthoff et al. 2022).

## Results

### “Ascunsovirus oldenburgi”ICBM5 – a novel *Microviridae* isolate

Phage ICBM5 was isolated from surface seawater, collected from the shoreline of the North Sea (53°42’09.8”N 7°41’58.9”E) during high tide in June 2015. The host, *Sulfitobacter* sp. SH24-1b, was isolated from a seawater sample taken on 12 May 2007 in the southern North Sea (54°42’ N, 06°48’ E) during a phytoplankton bloom (Hahnke et al. 2013). Its 16S rRNA has 99.8 % identity with *Sulfitobacter dubius* type strain DSM 16472^T^. According to its corresponding digital DNA-DNA hybridization value of 70 % and ANI of 96.9 %, strain SH24-1b belongs to the species *S. dubius*. A host range assessment performed on almost 100 bacterial strains showed that ICBM5 has a narrow host range, infecting only its original host *Sulfitobacter dubius* SH24-1b and *Sulfitobacter dubius* DSM 16472 ^T^ (SI file 1 Table S2). On *S. dubius* SH24-1b, ICBM5 formed turbid plaques.

ICBM5 has an icosahedral capsid and no tail, as revealed by TEM of uranyl acetate stained samples (Fig. 1A). The capsid diameter measured 28.68 +/- 1.95 nm (100 phages measured and three measurements per phage). Enzymatic digestion revealed that ICBM5 has an ssDNA genome (Fig. 1B). Sequencing and assembly resulted in a 5 581 bases long contig, circularly closed. Six protein-encoding genes were detected. Using BLASTp, only one ICBM5 protein was similar to proteins from previously classified *Microviridae*, namely the replication initiation protein (Rep). However, DELTA-BLAST, a remote homology tool, showed three more proteins distantly related to reference *Microviridae:* the major capsid protein (MCP), a pilot protein, and a lysis protein (see Fig. 1C). The presence of these *Microviridae* core genes, alongside its genome characteristics and virion morphology, clearly indicates that ICBM5 is a new member of this family. However, a first hint that ICBM5 is distantly related to known microviruses comes from the detection of MCP similarity to reference *Microviridae* only by a remote homology tool, although this protein is generally well conserved and often used to build *Microviridae* phylogenies. We consider thus that ICBM5 is the representative of a new phage species, which we tentatively named here “Ascunsovirus oldenburgi” ICBM5, using the binomial nomenclature recently adopted by ICTV.

**Figure 1.**
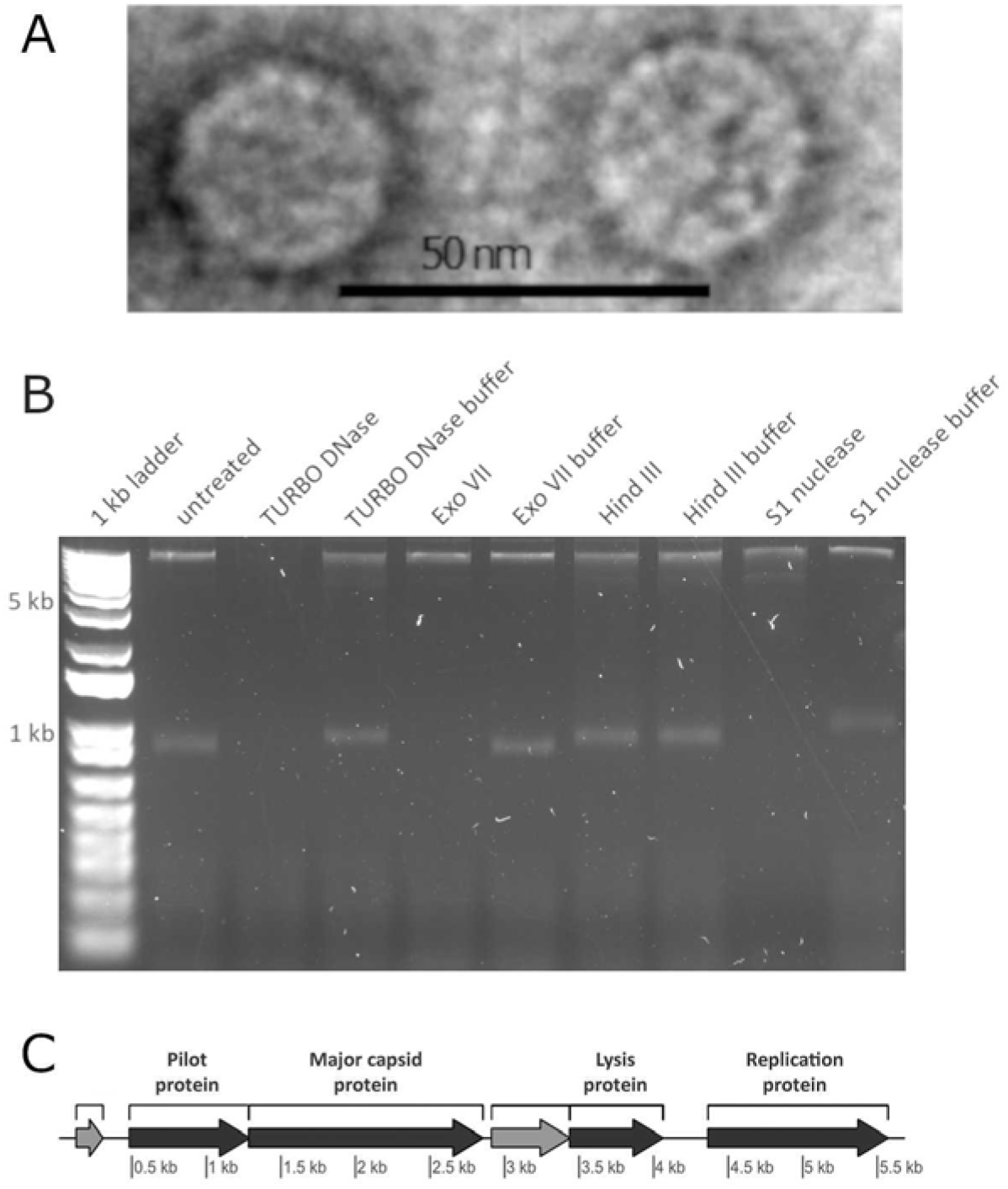
A. ICBM5 morphology determined by TEM of uranyl acetate stained virions. B. Agarose gel shows enzymatic digestion of ICBM5 ssDNA phage. The DNA was digested by TURBO DNase, Exo VII, and S1 nuclease, but was not affected by treating it via restriction enzyme Hind III, which only targets dsDNA, or the exclusion of nucleases (usage of only buffer). 1kb plus ladder was used as a molecular weight marker. C. Genome map of ICBM5. In dark grey - identified proteins, with labels on top of each gene. In light grey - hypothetical proteins.

### “Ascunsovirus oldenburgi” ICBM5 has both a lytic and a carrier state infection strategy on its *Sulfitobacter dubius* SH24-1b host

To characterize the infection cycle of the phage ICBM5, we performed one-step infection curves, in two separate experiments. The MOI was 6.5 in both experiments. Through the infections, samples were collected at different time points, in ^~^15 min increments. For each time point, we quantified: i) the free and total phages using plaque assays (see Fig. 2A); ii) the percentage of infected cells (see Fig. 2A) and iii) the variation of the amount of ICBM5 genomes per infected cell (see Fig. 2B for experiment 1, Fig. 3 and SI file 1 Fig. S3 for experiment 2). The latter two measurements were obtained by using ICBM5-targeted direct-geneFISH, a single cell method. We noticed a progressive increase of the per-cell ICBM5 genome numbers from ^~^50 min post-infection (p.i.) until ^~^110 min p.i.. At 50 min p.i., the median number of per cell ICBM5 genomes was 6.5x and 2x higher than at 35 min p.i., for experiment 1 and experiment 2, respectively. At 110 p.i, it was 61x and 81x higher, for experiment 1 and experiment 2, respectively. Therefore, ICBM5 was replicating its genome at least as early as 50 min post-infection. Cell lysis events were visually observed between 110 min and 140 min p.i. (see Fig. 3) in both experiments. In agreement, the free phage particles increased in numbers starting with ^~^110 min p.i.. This corresponded with the decline of the cell population with a high amount of ICBM5 genomes, and the emergence of a cell population with a low amount of ICBM5 genomes, as indicated by the progressive drop in the median number of ICBM5 genomes per cell (see Fig. 2B). The population with a low amount of ICBM5 genomes was stably maintained from 155 min to 185 min p.i., when the experiments ended. Therefore, the major lysis event took place starting at 110 min post-infection and newly released virions infected new cells, leading toward a second wave of infection. Overall, these results show that ICBM5 can undergo a complete lytic cycle on *S. dubius* SH24-1b.

**Figure 2.**
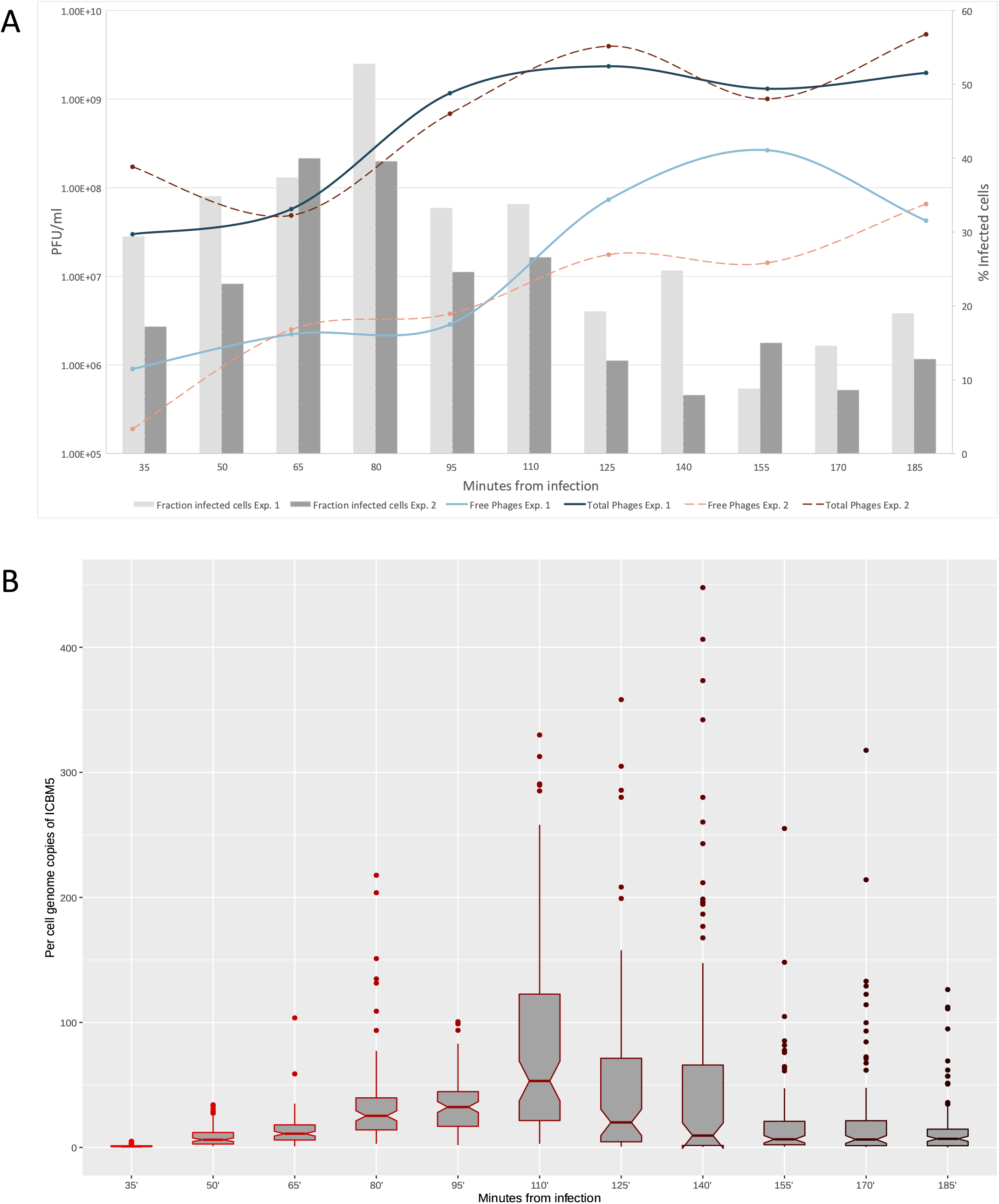
ICBM5 - *Sulfitobater dubius* SH24-1b infection dynamics. A. The fraction of ICBM5 infected cells (bars) and the abundance of free (extracellular) and total (intra- and extracellular) phages, given in PFU/ml (lines). The fraction of infected cells was calculated from the proportion of cells showing a phage signal, after subtraction of the false-positive signals detected in the negative control cultures (see SI file 1 text). B. The variation of the per-cell ICBM5 genome copies through the infection, as calculated from measuring the phage signal intensities using direct-geneFISH (see SI file 1 text). The box plot borders represent the 1^st^ and the 3^rd^ quartile, and the middle line represents the 2^nd^ quartile. The whiskers extend from the 1st or 3rd quartiles with 1.5 * IQR (distance between first and third quartiles). The data beyond the whiskers are plotted individually. The plot was generated using the ggplot2 R package (Hadley 2016). The different shades of red for the box plots are just indicating the progress of the infection time, from bright red at the beginning of the experiment, to dark red at the end of the experiment.

**Figure 3.**
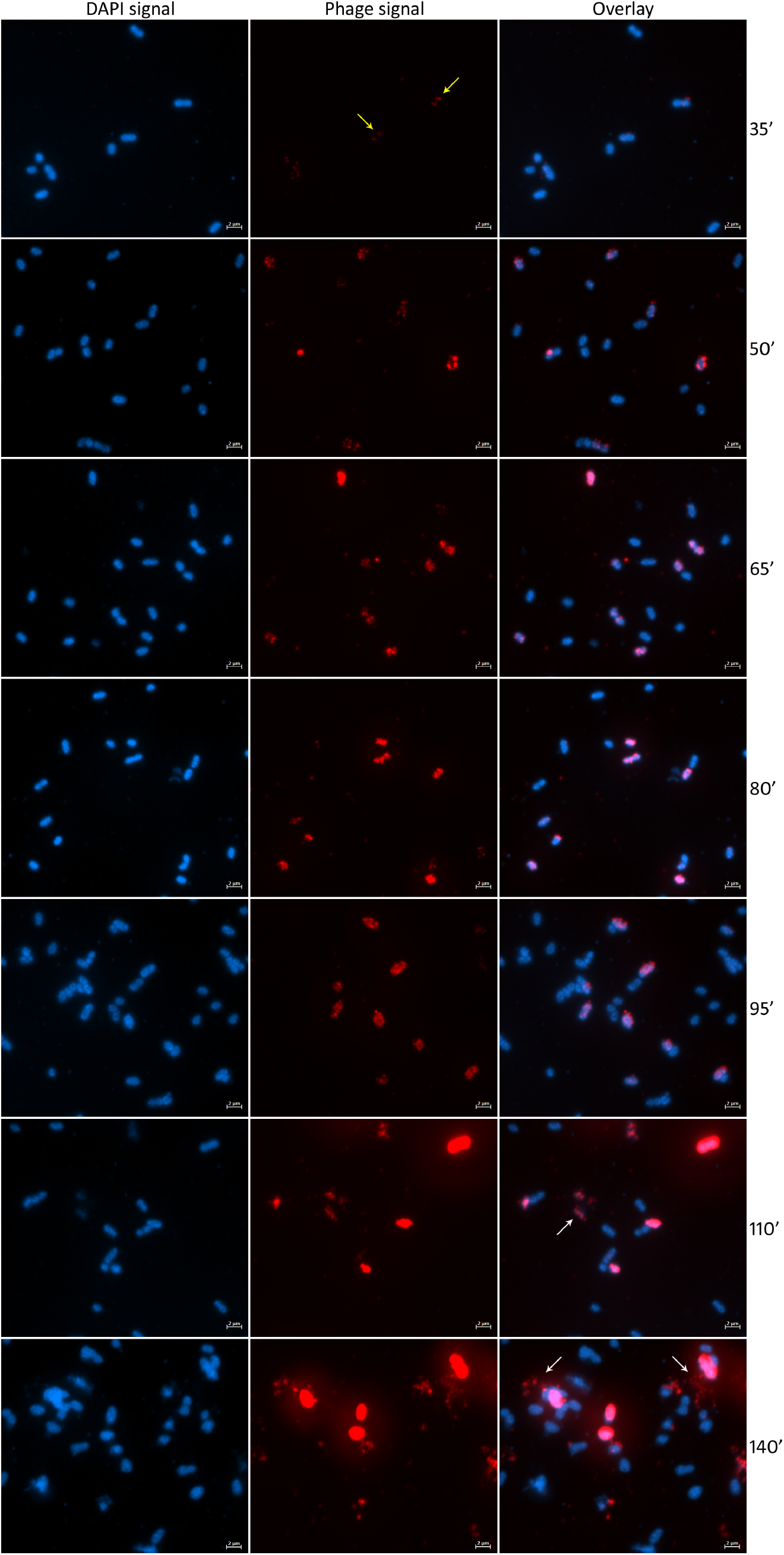
Visualization of the infection of *Sulfitobacter dubius* SH24-1b by ICBM5 using direct-geneFISH. First column: cells as visualized by DAPI staining. Second column: intracellular phages, as visualized by ICBM5 genome-targeted probes. Third column: overlay of DAPI and phage signal. The phage signal increases progressively through the infection. The very small dots in the first time point (yellow arrows) represent early infections, with just a few copies per cell. Larger, cell-wide signals in the following time points represent phages in the replication and maturation phase. And finally, at 110’ and 140’, cell lysis events can be noticed (white arrows) phages.

The fraction of ICBM5 infected cells reached a maximum at 80 min p.i., after which it progressively decreased (see Fig. 2A). This, together with the ability of ICBM5 to form turbid plaques, suggested the emergence of a resistant *S. dubius* SH24-1b sub-population. To test if the resistance was conferred by the integration of ICBM5 as prophage in the host cells, we collected surviving bacterial cells from turbid plaques and plated them to obtain single colonies that we screened by PCR for the presence of the phage ICBM5. When ICBM5-positive cultures derived from the phage-positive colonies were challenged with ICBM5 in a spot assay, no clearing zones were formed. Therefore, the new cultures were resistant to ICBM5. Using the Nanopore long-read sequencing technology on the entire genome, without any size-exclusion during library preparation, the phage ICBM5 was detected as an independent, circular contig. Using PacBio long-read technology, using a 7 kb size selection threshold for library preparation, ICBM5 specific sequences were not detected. Therefore, no evidence of integration in the bacterial chromosome was found, that is, no hybrid ICBM5 – *S. dubius* SH24-1b reads were present. ICBM5-targeted direct-geneFISH on this resistant culture showed that ICBM5 was present in about 4.5 % of the cells. The geneFISH signal varied in between cells. Some cells had small, dot-like signals, as characteristic of a low number of ICBM5 genome copies. Other cells had larger, diffuse signals, indicating the presence of a higher number of ICBM5 genomes. No cell lysis events were noticed. Together, these results showed that ICBM5 does not undergo lysogeny as integrated prophage in the tested conditions. However, ICBM5 can survive and replicate in a sub-population of sensitive *S. dubius* SH24-1b cells, which co-exists in parallel with a dominant ICBM5 resistant sub-population. This is indicative of a carrier state infection strategy.

### ICBM5 related genomes are widespread within bacterial genomes, both as prophages and episomes, and in environmental viromes

To better understand the spread of ICBM5-related phages in bacterial hosts and environmental samples, as well as their phylogenetic classification, we searched for similar genomes in two publicly available data sources: bacterial genomic data and viral metagenomes.

First, ICBM5 proteins were used to find potential prophages within prokaryotic genomes. Out of the 72 Microviridae-like genomic regions found in bacterial genomes, 7 have been previously described (Krupovic and Forterre 2011; Quaiser et al. 2015; Zheng et al. 2018) and 65 are new (see Fig. 4 and see SI file 4 Table 1 and SI file 5). For 39 of them, we were able to determine the borders, by comparing them with related bacterial strains free from these regions. For the rest, we narrowed down the borders by keeping only proteins with a clear phage origin (see Materials and methods). The length of these regions varied between 3.3 and 6.6 kbs for clear border regions, and between 3.5 and 8.2 kbs for unclear border regions. The majority of Microviridae-like genomic regions occurred in bacteria from *Alphaproteobacteria* (53.5%), followed by *Bacteroidia* (29.5%) and *Gammaproteobacteria* (5.6%). The rest was distributed among *Bacilli, Chlamydiae, Clostridia, Cyanophyceae, Erysipelotrichia, Flavobacteriia*, and *Negativicutes* (see Fig. 5). Most of the new Microviridae-like genomic regions had a chromosomal localization and three were localized on large plasmids (0,1 Mb to 1.6 Mb, see SI file 4 Table 1), indicating they are *bona fide* prophages. Seven genomic regions consisted of small contigs similar in size to *Microviridae* genomes, most likely representing episomes (see SI file 4 Table 3 and Fig. 4) from a carrier state life strategy.

**Figure 4.**
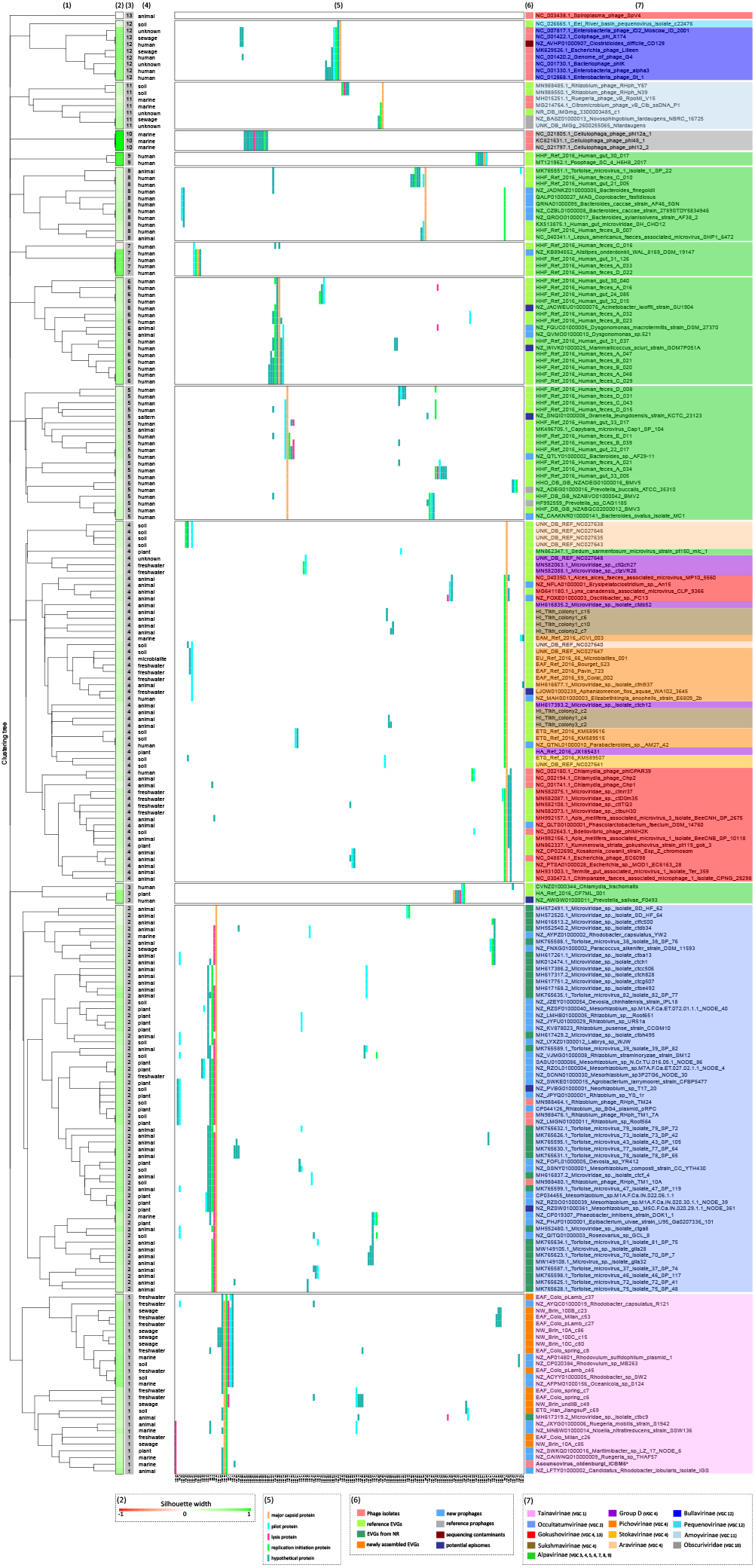
Hierarchical clustering of the ssDNA (pro)-phage genomes, based on their protein super-super clusters content. The annotations show the following: 1) Hierarchical clustering tree; 2) Silhouette width, measures on a scale from −1 to 1, how related is a viral genome with other genomes in the same genome cluster; 3) Viral genome cluster (VGC) number; 4) Habitat of the phage or host; 5) Distribution of the protein clusters (PSSCs) in each viral genome. Protein clusters not shared with any other phage genomes in this dataset are not shown. The color encodes different protein annotations; 6) Phage genome category. EVGs = environmental viral genomes; 7) The label given to each phage genome, consisting of accession numbers and names of the phage isolate, or the environmental contig or of the bacterial host in which a (pro)-phage was predicted.

**Figure 5.**
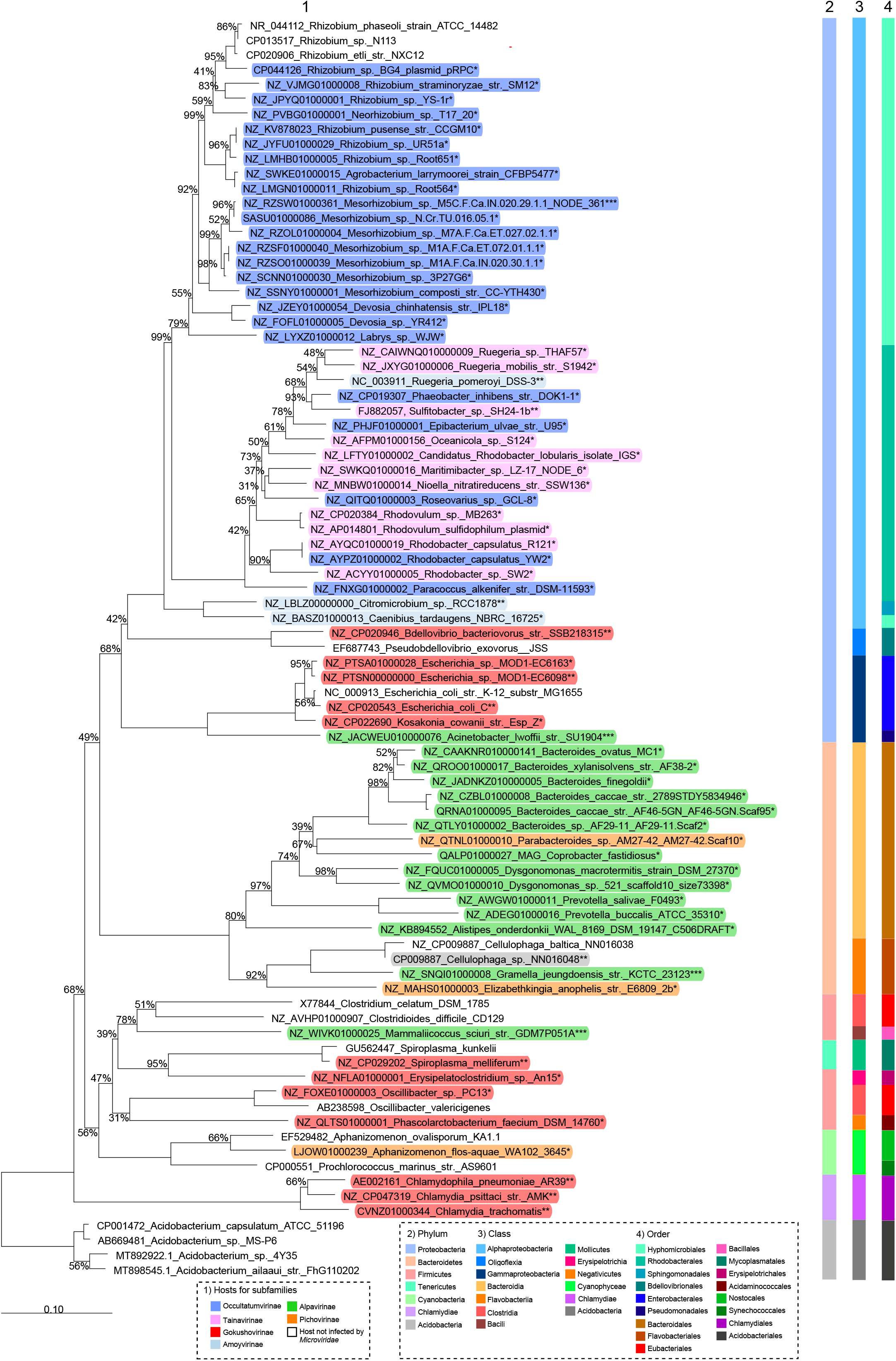
16S rRNA phylogenetic analysis of the (pro)-phage hosts. Neighbor-joining tree based on the 16S rRNA gene sequence similarity showing the position of *Sulfitobacter dubius* SH24-1b and other bacterial hosts for *Microviridae*-like (pro)-phages. The 16S rRNA gene was not detected for four bacterial strains in Figure 4 and therefore not included in this tree. Bootstrap values are derived from 1000 replicates. GenBank accession numbers are given as prefixes, followed by species and strain names. The bar represents 10 substitutions per nucleotide position. The stars encode the following: * hosts of predicted prophages, integrated into chromosomes or plasmids; ** hosts of isolated phages; *** hosts of predicted episomes, represented by short contigs.

In addition to these potential prophages, we also searched for ICBM5-related genomes amongst EVGs sequenced from virions from environmental samples, that can be found in NR and public viromes. We have retrieved 31 environmental phage genomes from our search of the NR NCBI database, alongside 23 EVGs already affiliated to know microvirus subfamlies. Their size ranged between 4.2 and 6.6 kbs. These NR sequences were generated from 10 viromes associated with humans, animals, or plants (see Fig. 4 and SI file 6). In addition, we used the MCP from ICBM5 to screen 2 944 previously published viral metagenomes yet only available as raw reads and that had to be newly assembled in this study. A total of 15 circular contigs representing potential full-length genomes were retrieved in 8 different viromes from fresh- or reclaimed water and soil (see Fig. 4 and SI file 2).

All newly found prophages, episomes, and EVGs were then pooled with ICBM5 and reference *Microviridae* genomes and compared in terms of gene content using VirClust. Their proteins are of course related and formed clusters with proteins from cultivated and uncultivated reference *Microviridae*. They shared no protein clusters with the *Obscuriviridae*, which are also ssDNA phages with icosahedral capsids. Most genomes newly detected here had the usual *Microviridae* proteins: pilot, MCP, lysis, and Rep proteins, with inter-spreading hypothetical proteins. Hierarchical clustering of the genomes based on their protein cluster content resulted in 13 major viral genome clusters (VGCs) (see Fig. 4). Each VGC had its own set of protein clusters, with few protein clusters being shared between the VGC. The phylogenetic analysis of the MCP and Rep proteins was generated to have a more precise idea of the relationships between these viruses and was coherent with the VirClust analysis as the 13 genome clusters mostly corresponded to major phylogenetic clades (see Figs. 4 and 6).

**Figure 6.**
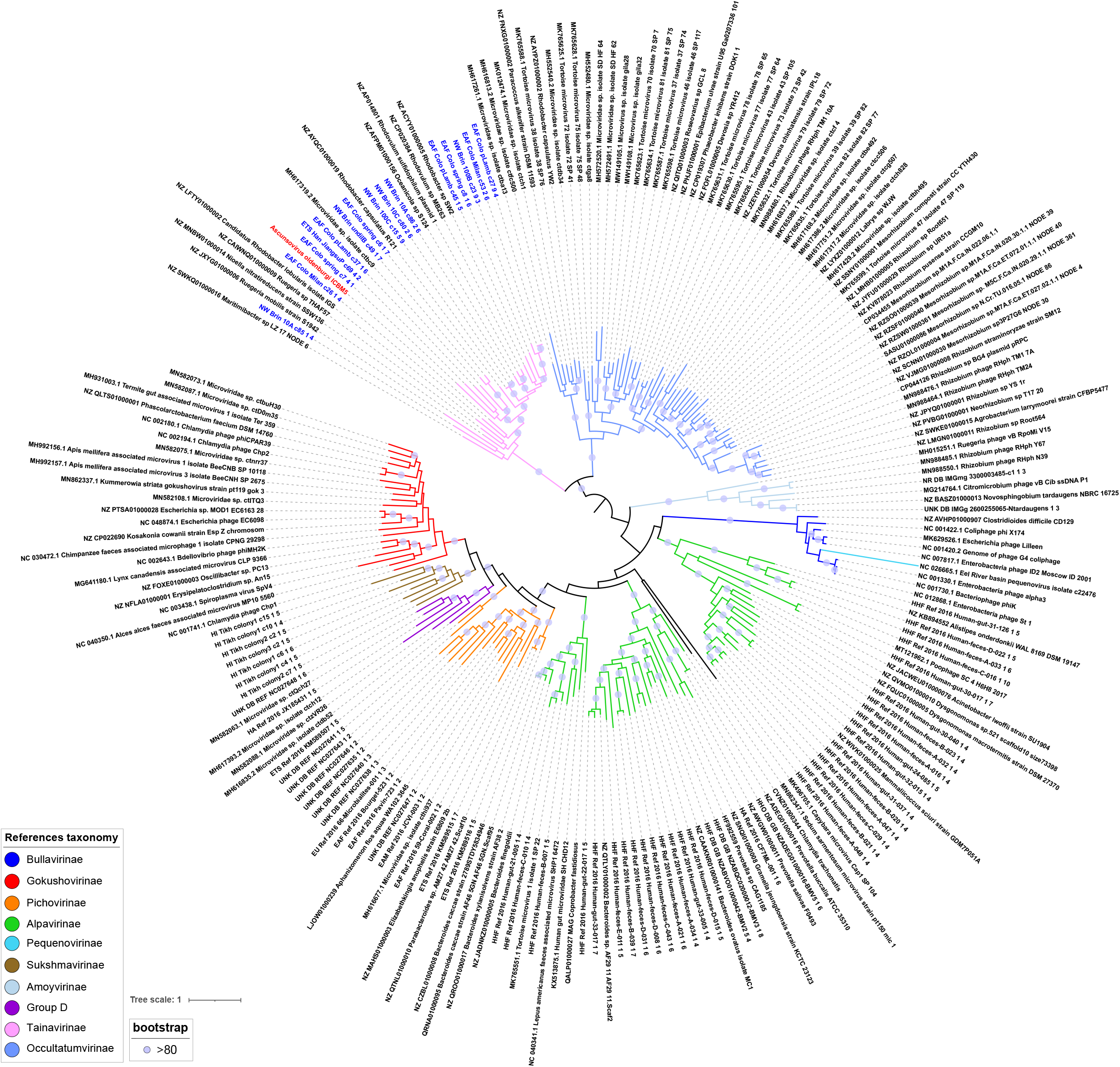
MCP and Rep based phylogeny. Phylogeny on the concatenated major capsid and replication proteins of the ssDNA (pro)-phage genomes. Branches are colored according to the taxonomic affiliation into subfamilies. Names are colored corresponding to the genome clusters from the VirClust analysis, see Fig. 3. Clear blue circles are placed on internal branches when bootstrap values are over 80. ICBM5 is highlighted in red, while closely related virome contigs are in blue (BLASTp on major capsid proteins, bit-score>300).

About half of the newly predicted prophages, episomes, and EVGs, as well as phage ICBM5, were gathered in a group separated from previously defined *Microviridae* subfamilies (Fig. 6). Viruses from this large group were clustered in two clades in the phylogenetic tree (Fig. 6) that correspond to VGC1 and VGC2 in the VirClust analysis (Fig. 4). The first cluster comprised ICBM5 as the only cultivated phage, alongside sequences newly identified. Indeed, VGC1 encompassed 10 newly found prophages and all the 15 EVGs newly assembled in this study from 8 viromes generated in 3 different previous studies (Colombo et al. 2017; Han et al. 2017; Brinkman et al. 2018). Only the contig MH617319.2_Microviridae_sp._isolate_ctbc9 was previously identified as *Microviridae*, but not further classified (Tisza et al. 2020). The second cluster was larger, with 21 new prophages and two episomes, 30 EVGs, and three recent isolated phages infecting *Rhizobium* (van Cauwenberghe et al. 2021). All the EVGs here were previously identified as *Microviridae* in different viromics studies (see SI file 6) (*Creasy et al. 2018; Orton et al. 2020; Tisza et al. 2020; Collins et al. 2021)*. In two of the studies, the respective EVGs were recognized already to represent a separate group from the known *Microviridae* subfamilies (Creasy et al. 2018; Orton et al. 2020). However, no further classification of these phages was performed.

Considering that two VGCs are generated, clearly separated from each other and even more distantly related to known Microviridae subfamilies in the phylogeny, we tentatively propose here two new subfamilies: i) “Tainavirinae” (from the Romanian word “taina”, which means secret), representing the clade with phage ICBM5; and ii) “Occultatumvirinae” (from the Latin word “occultatum”, which means hidden), representing the clade with the *Rhizobium* phages. The genomic diversity within the two new subfamilies was high, with the nucleotide-based intergenomic identity ranging between 0.0 % and 99.9%. Most of the phage pairs had an intergenomic identity lower than 40%. Few phages had an intergenomic identity above 95%, which would place them into the same species: five EVGs into two species in the “Tainavirinae”, and two *Rhizobium* phages into one species in the “Occultatumvirinae” (see SI file 7 and SI file 8).

The two subfamilies comprise phages present in different environments (Fig. 4) and spread worldwide (Fig. 7), with only *Alphaproteobacteriaas* known hosts (Fig. 5). Tainavirus prophages and ICBM5 were only found in *Rhodobacterales* hosts. Most occultatumvirus prophages, episomes, and cultivated phages infected *Hyphomicrobiales*, the rest infecting the related bacterial order *Rhodobacterales*. A habitat overview showed that these phages, including the EVGs, and their hosts are usually found in the terrestrial and marine environment, often in association with plants and animals (see Fig. 4, SI file 4, Table 2, and SI file 6). We found occultatumvirus EVGs in viromes from the lizard *Heloderma suspectum* (Collins et al. 2021), the fishes *Carassius carassius, Lutjanus campechanus* and *Pimephales* sp., the snail *Haliotis sp*. (Tisza et al. 2020), the tortoise *Gopherus morafkai* (Orton et al. 2020) and the sea squirt *Ciona robusta* (Creasy et al. 2018). All *Hyphomicrobiales* infected by *Occultatumvirinae* were isolated from soil or plant root nodules. The *Rhodobacteraceae* infected by occultatumviruses were isolated either from the soil, or marine algae and seawater (see SI file 4, Table 2). We found tainavirus EVGs in viromes from paddy soils (Han et al. 2017), freshwater rivers (Colombo et al. 2017), and wastewater (Brinkman et al. 2018). The *Rhodobacteraceae* hosts infected by tainaviruses were isolated from terrestrial environments, including ponds, sediments, and soil, and marine environments, including sediments, water column, sponges, copepods, and dinoflagellate (see SI file 4, Table 2).

**Figure 7.**
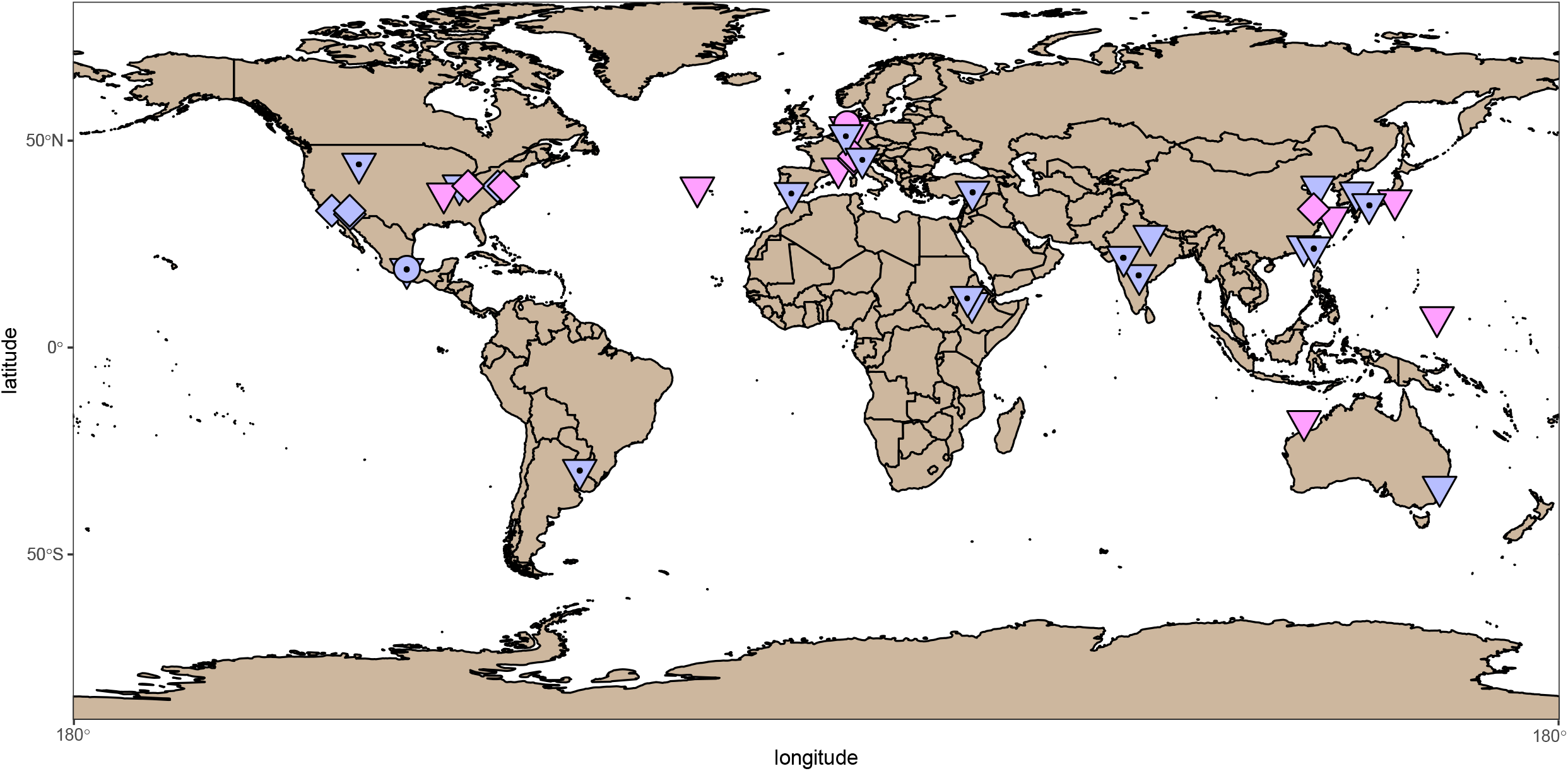
Biogeographical distribution of tainaviruses (pink) and occultatumviruses (lavender). The points on the map represent the sampling place for phage isolates (circles), for bacterial hosts harboring prophages and episomes (triangles), and for the viromes from which the EVGs have been assembled (squares). The triangles with a central circle represent locations for *Rhizobium* and *Mesorhizobium* infected by occultatumviruses.

## Discussion

Knowledge of the diversity of tailless, icosahedral ssDNA phages is still in progress, as evidenced by the constant sequencing of new viral genomes. As a result, their classification is under regular revision, as it is clear by now that there are several major clusters of icosahedral ssDNA viruses. These are defined so far as subfamilies – *Bullavirinae, Gokushovirinae, Pichovirinae, Aravirinae, Alpavirinae, Stockavirinae, Pequenovirinae, Sukshmavirinae*, and *Amoyvirinae* –, and grouped under the umbrella of the *Microviridae* family (Creasy et al. 2018; Zheng et al. 2018). This study illustrates such a process, because (i) the new phage isolated here is far from references, and (ii) the use of this phage as a stepping-stone led to lifting the veil on two new major groups of *Microviridae* – “Tainavirinae” and “Occultatumvirinae”. These proposed new subfamilies represent a major contribution to the known *Microviridae* diversity, as indicated by the MCP-Rep phylogeny, where the two groups are distant from all current subfamilies (see Fig. 6).

In addition, the data collected here extends our understanding of *Microviridae* lifestyle. So far, only a handful of microvirus-like prophages were predicted bioinformatically, contrarily to inoviruses, the other major group of ssDNA phages. Indeed, inoviruses have been a long time known to integrate into their host genomes, as part of their chronic life cycle (Mai-Prochnow et al. 2015) and a recent bioinformatics study found a huge diversity of inovirus-like prophages, spread throughout many bacterial and archaeal taxa (Roux et al. 2019). Here, the cultivation and sequencing of ICBM5 enabled the prediction of many more microviridae-like (pro)-phages. Furthermore, the known host clades, so far restricted to the *Bacteroidia* class, *Enterobacteraceae* family (*Gammaproteobacteria* class), and one *Hyphomicrobiales* family (*Alphaproteobacteria* class) (Krupovic and Forterre 2011; Roux et al. 2012; Quaiser et al. 2015; Zheng et al. 2018), were here expanded to include several *Firmicutes* classes*, Flavobacteriia*, and new *Alphaproteobacteria* and *Gammaproteobacteria*. Within the *Alphaproteobacteria*, this is the first report of prophages in genomes of the *Rhodobacteraceae* family, an environmentally significant clade in the marine environment. *Microviridae* prophages have been recently reported in a few genomes from *Rhodobacteraceae (Forcone et al. 2021)*, however, a closer look revealed that they are phiX174, most likely an unremoved addition from the sequencing process. The many (pro)-phages we found both in *Hyphomicrobiales* and *Rhodobacterales* are unrelated with the *Amoyvirinae* prophage previously found in *Caenibius tardaugens (Hyphomicrobiales)* (Zheng et al. 2018). Similarly, they are not related to the previously isolated *Ruegeria pomeroyi* phages, which according to our analysis belong to *Amoyvirinae*.

Occultatumviruses and tainaviruses with known hosts infect *Hyphomicrobiales* and *Rhodobacterales*, but of course, hosts are not known for the EVGs. Yet, their phylogenetic relatedness to phage isolates and prophages, and the homogeneity of the known bacterial hosts suggest that these EVGs also infect *Rhodobacterales* or *Hyphomicrobiales*. Indeed, representatives of these two bacterial orders have been found in habitats similar to those in which the EVGs were found, either in diverse aquatic systems (sewage, freshwater, marine), soil or associated with eukaryotes (animals, plants, or microalgae). For example, *Rhodobacteraceae* were found in the gut of a *Ciona* species, suggesting that the occultatumvirus EVGs found in *Ciona robusta* could also infect bacteria in this family. Similarly, *Rhodobacteraceae* and *Hyphomicrobiales* were found in the gut or skin of similar fish species from which occultatumvirus EVGs were retrieved (DeBofsky et al. 2020; Nielsen et al. 2018; Tarnecki et al. 2022). In the same vein, the tortoise *Gopherus morafkai* harbors in its nasal microbiome both *Hyphomicrobiales* and *Rhodobacterales (Weitzman et al. 2018)*, making members of the two orders very likely the hosts of the occultatumviruses found in the fecal viromes from similar tortoises. Furthermore, *Rhodobacteraceae* are common in wastewater (Numberger et al. 2019), freshwater rivers (Liu et al. 2019), paddy soils (Wang et al. 2018), and in marine sediments and water column (Hahnke et al. 2013). For example, here, *Rhodovulum sp* MB263 and *Rhodovulum sulfidophilum*, which belong to the same species (Fig 5) are respectively found in soil and marine systems. Accordingly, their tainaviruses appear as closely related (Fig 4 and 6). *Rhodobacteraceae* are also known to have a free or associated lifestyle. For example, the tainavirus host *Epibacterium ulvae (Breider et al. 2019)* was isolated from macroalgae. It is thus very likely that tainavirus EVGs also infect *Rhodobacteraceae* living in diverse aquatic systems (sewage, freshwater, marine) and soil. Concerning occultatumviruses, which infect two bacterial orders, *Rhodobacterales* and *Hyphomicrobiales*, it has to be noted that these orders are closely related in the bacterial tree. Furthermore, all occultatumviruses infecting *Hyphomicrobiales* form a monophyletic group (Fig 6). The *Hyphomicrobiales* order, containing *Rhizobium* and *Mesorhizobium* strains, is particularly interesting. These bacteria are commonly found in soil and, in conditions of nitrogen starvation, they migrate in the root hairs of legumes, where they transform into nitrogen-fixing endosymbionts (Poole et al. 2018; Clúa et al. 2018). Considering the worldwide spread of both wild and cultivable, economically important legumes (Sprent et al. 2017), it can well be that occultatumviruses are cosmopolitan. Certainly, the varied geographical locations in which occultatumvirus harboring *Rhizobium, Neorhizobium and Mesorhizobium* were found (see Fig. 7) support this hypothesis. To summarize, the two related groups “Tainavirinae” and “Occultatumvirinae” infect respectively only *Rhodobacterales* or *Hyphomicrobiales* and *Rhodobacterales*, two related orders of *Alphaproteobacteria*, reinforcing our belief that these viral groups are evolutionnary linked. Either the ancestor of these viruses was already infecting the ancestor of this branch of *Alphaproteobacteria* or the viral ancestor infected only *Rhodobacterales* and during the evolution it infected *Hyphomicrobiales*. Concerning the rest of the *Microviridae*, some subfamilies proposed earlier, like Pichovirinae and Alpavirinae, appear to be polyphyletic when adding new prophages and EVGs. In addition, these two subfamilies infect respectively 2 and 3 bacterial phyla, suggesting that these subfamilies might be too large and need to be revised.

Phage infections are usually described in terms of lytic, lysogenic, or chronic lifestyles. Alternative lifestyles, such as pseudolysogeny or carrier state have been observed among different phage groups, although their definition is not always consistent and the two terms have been used interchangeably (reviewed in (Mäntynen et al. 2021)). The same phage can exhibit different lifestyles on the same host. For example, the P22 phage displayed the following lifestyles when infecting its host, *Salmonella* Typhimurium: i) a lytic strategy resulting in cell lysis, ii) a pseudolysogenic strategy characterized by the existence of a P22 episome, which, after cell division, was transmitted only to one of the daughter cells, and iii) a lysogenic strategy, arising from the cell which inherited the episomal phage (Cenens et al. 2015). The discovery of prophages in other *Rhodobacteracea* has prompted us to ask if ICBM5, in addition to its lytic strategy, also has a lysogenic lifestyle. When investigating an ICBM5 resistant strain isolated from turbid plaques, we found no evidence that ICBM5 integrates into the genome of *S. dubius* SH24-1b. However, we found that ICBM5 can undergo a carrier state life strategy. A sub-population of resistant strain cells carried the ICBM5 genome as an episome present in variable numbers. A second, numerically dominant sub-population carried no ICBM5, likely being resistant to this phage and conferring to the host strain the resistance to ICBM5 observed in spot assays. Considering that this strain was isolated from a single colony, there are two possible mechanisms by which the two sub-populations were produced. In the first scenario, the single cell from which the colony arose harbored the ICBM5 genome intracellularly. Asymmetrical cell division would have resulted in transmission of ICBM5 only to one of the daughter cells. In a second scenario, the ICBM5 phage particle somehow became attached extracellularly to the initial colony-forming cell. Upon subsequent cell divisions, sensitive cells would have arisen and would have become infected by ICBM5. What factors confer resistance to the ICBM5 is for now unknown. During P22 infection of *Salmonella* Typhimurium, P22-free daughter cells resulting from the asymmetric division of pseudolysogenic cells were transiently immune to P22. The resistance was conferred by immunity factors cytoplasmically transmitted from the mother cell and thus, inevitably diluted by subsequent cell divisions (Cenens et al. 2015). Considering the high proportion of not-infected cells in our ICBM5 resistant strain, it is unlikely that a similar mechanism is responsible for conferring resistance. Further experiments are required to characterize the ICBM5 carrier state life strategy and to elucidate the host resistance mechanism. The carrier state does not seem to be confined to infection of *S. dubius* SH24-1b by ICBM5. We predicted *Microviridae-like* episomes in *Mesorhizobium, Neorizobium, Prevotella, Aphanizomenon, Gramella, Mammalicoccus*, and *Acinetobacter*, hosts belonging to various phyla. Likely, these phages used a carrier state life strategy to survive in their host cultures, without having a dramatic effect on the culture’s growth. Furthermore, a similar carrier state was recently shown for a gokushovirus “revived” from its prophage state in the host genome by molecular cloning (Kirchberger and Ochman 2020). Together, this indicates that such a carrier state is spread amongst *Microviridae* phages, very likely enabling co-existence with their hosts in environmental samples.

Fluorescence *in situ* hybridization targeting phage genes was previously used to characterize the lytic life cycle of PSA-HP1, a *Pseudoalteromonas* infecting dsDNA phage (Allers et al. 2013). The method used multiple probes labeled with digoxygenin, followed by a signal amplification step mediated by antibody binding and enzymatic tyramide deposition. Here, we have applied the relatively new direct-geneFISH method (Barrero-Canosa et al. 2017; Barrero-Canosa and Moraru 2021b), which uses multiple probes directly labeled with fluorochromes and thus avoids signal amplification steps. This method was used recently for intracellular virus detection of dsDNA archaeal and picoeukaryotic viruses in environmental samples and pure cultures (Castillo et al. 2020; Castillo et al. 2021; Rahlff et al. 2021). In our study, using phage-targeted genome-wide probes, we were able to characterize the lytic cycle of an ssDNA phage. The time until lysis for ICBM5, in the tested conditions, was 110 min. This is shorter than the ^~^3 h reported for the vB_Cib_ssDNA_P1 phage infecting *Citromicrobium* sp. (Zheng et al. 2018) and the vB_RpoMi_Mini infecting *Ruegeria pomeroyi* DSS-3 (Zhan and Chen 2019b). In contrast, the duration of the lytic cycle for phiX174, the best characterized to date *Microviridae*, was only 20 min on *Escherichia coli* (Hutchison and Sinsheimer 1963). The difference comes most likely not only from the genetic differences between the phages, but also from the differences in the physiology of the two hosts. Furthermore, by measuring the total phage signal intensity values per cell and normalizing them to the intensity of a single phage, we were able to quantify the per cell genome numbers of ICM5. In the last phases of the lytic cycle, most of the cells had up to 125 ICBM5 genome copies. However, some cells reached more than 300 ICBM5 genome copies. Our measurements do not show how many of these genomes were packed into mature virions and released from the cells. However, these values are similar to the burst size of 250 phages per cell calculated from the PFU measurements. In comparison, phiX174 and vB_Cib_ssDNA_P1 have a burst size of ^~^170 phages per cell (Hutchison and Sinsheimer 1966), and vB_RpoMi_Mini of only ^~^8 phages/cell.

The discovery of so many new microviridae-like prophages and episomes raises the question of whether they are more widespread in bacterial genomes than previously recognized. Our prophage prediction approach using ICBM5 proteins to search the NCBI-Blast database with DELTA-BLAST, followed by several rounds of PSI Blast, is a relatively unsophisticated procedure. It was able to find not only ICBM5-related prophages but also distant relatives, for example, prophages of *Bacteroidetes* which group with Alpavirinae or Gokushovirinae. This suggests that most of the findable prophage and episome diversity at the date of our search has been recovered. The addition of new bacterial genomes or the discovery of new microviridae-like sequence diversity, either by phage cultivation or metagenomics, could reveal further microvirus diversity in bacterial genomes.

## Acknowledgments

This work was supported by the Deutsche Forschungsgemeinschaft (DFG) within the Transregional Collaborative Research Centre Roseobacter (TRR51). We thank Cathrin Spröer for sequencing support. The work conducted by the U.S. Department of Energy Joint Genome Institute (SR), a DOE Office of Science User Facility, is supported by the Office of Science of the U.S. Department of Energy under contract no. DE-AC02-05CH11231.

## Author contribution

CM designed the study, isolated the phage ICBM5, carried out part of the prophage prediction and phage classification using VirClust, analyzed the data, and wrote the manuscript. FZ performed out the direct-geneFISH, the experiments proving the ssDNA nature of ICBM5 and the Nanopore sequencing, carried out part of the prophage prediction and phage classification using VirClust, analyzed the data, and wrote the manuscript. VB performed TEM and host range assays for ICBM5, calculated the 16S tree, and wrote the manuscript. Genomes of *S. dubius* SH24-1b strains based on PacBio sequencing were generated by HF. BH isolated the ICBM5 resistant strain. AP sequenced the ICBM5 phage. SR, EON, and FE constructed the MCP-REP tree and participated in the phage classification. All authors revised the manuscript.

