## Supplementary material for "New Microviridae isolated from Sulfitobacter reveals two cosmopolitan subfamilies of ssDNA phages infecting marine and terrestrial Alphaproteobacteria": SI_file_1

### Title:

The authors declare no conflict of interest.

Keywords: ssDNA phages, *Microviridae*, *Tainavirinae*, *Occultatumvirinae*, Ascunsovirus oldenburgi

Running title: ICBM5 ssDNA phage and like prophages

#### Materials and methods

##### Growth media

Marine broth (MB) was used both for the liquid cultures and for the plaque and spot assays. This media had the following recipe. 5.0 g/l peptone, 1.0 g/l yeast extract, 0.1 g/l C_6_H_8_FeO_7_, 12.6 g/l MgCl_2_x6H_2_O, 3.24 g/l Na_2_SO_4_, 19.45 g/l NaCl, 2.38 g/l CaCl_2_x2H_2_O, 0.55 g/l KCl, 0.16 g/l NaHCO_3_, 0.01 g/l Na_2_HPO_4_x2H_2_O, 0.08 g/l KBr, 0.034 g/l SrCl_2_x6H_2_O, 0.022 g/l H_3_BO_3_, 0.004 g/l Na_2_SiO_3_x3H_2_O, 0.0024 g/l NaF, 0.0016 g/l NH_4_NO_3_. After autoclavation, the media was completed by adding 1 ml/l of a multi-vitamin solution (Balch *et al.* 1979). Artificial sea water (ASW) base medium was used for plaque assays or one-step infection experiments. This media had the following recipe. 24.32 g/l NaCl, 10 g/l MgCl_2_x6H_2_O, 1.5 g/l CaCl_2_x6H_2_O, 0.66 g/l KCl, 4 g/l Na_2_SO_4_, 2.38 g/l HEPES, 1 ml KBr (0.84 M), 1 ml H_3_BO_3_ (0.4 M), 1 ml SrCl_2_ (0.15 M), 1 ml NH_4_Cl (0.4 M), 1 ml KH_2_PO_4_ (0.04 M), 1 ml NaF (0.07 M).

##### Phage plaque assays using the double-agar layer method

To obtain single plaques, serial dilutions (10^0^, 10^-1^, etc.) were prepared from the phage fractions by mixing with MB medium. Further, 100 µl of phage dilution were mixed with 280 µl of exponentially growing host culture (OD = 0.2 - 0.3) and incubated for 15 min on ice. The mixture was transferred to 3 ml MB-soft agar (0.6 % low melting point Biozym Plaque GeneticPure agarose, Biozym, kept warm at 37°C), mixed by brief vortexing and poured onto the bottom MB agar layer (1.8 % agar). After drying of the top layer, the plates were incubated at 20 °C. For isolation, when phage plaques were observed as clearing zones within the grown bacterial lawn, they were picked with sterile Pasteur pipettes and incubated overnight in 500 µl ASW base at 4 °C. After subsequent centrifugation (10 min, 10000 x g, 4 °C), the supernatant was used for a next round of plaque assays. The procedure of plaque assay, picking of plaques and re-plating was repeated three times to ensure purity of the newly isolated phages.

##### Preparation of phage ICBM5 stocks

To obtain a larger volume of phage lysate, plaque assay was performed with 20 plates, using a dilution of an ICBM5 phage plaque resulting in confluent plaques.

Liquid infection culture was set up, by inoculation of 30 ml MB medium in an Erlenmeyer flask with an exponentially growing *Sulfitobacter* sp. SH24-1b culture (final OD = 0.006) and infected with a single ICBM5 phage plaque, picked with a sterile Pasteur pipette. A second culture was not infected and regarded as control. After overnight incubation at 20 °C and 100 rpm, the phage lysate was obtained by removing cells and debris by centrifugation (15 min, 4000 x g and 4°C) and 0.2 µm filtration. The phage lysate was stored at 4 °C. For long term storage, two types of glycerol stocks were prepared: i) stock of free phage particles (1 part phage fraction and 1 part MB media with 50% glycerol) and ii) stock of infected host cells (1 part infected cells - 375 µl phage fraction added to 375 µl host culture, 15 min on ice for absorption - and 1 part MB media with 50% glycerol).

##### Host range determination by spot assay

94 different strains of the *Rhodobacteraceae* family (see Table S2) were challenged with the purified ICBM5 phage by spot assay. For the spot assay, 280 µl of exponentially growing host culture (OD = 0.2 ‑ 0.3) were mixed with 3 ml MB-soft agar and poured onto the bottom MB agar layer (1.8 % agar). After drying of the top layer, 15 µl of phage fraction, obtained from a liquid infection as described above, were spotted in triplicates onto the top layer. For each strain, three plates were prepared this way and incubated at 15 °C, 20°C or 28 °C. Upon observation of lysis, the respective bacterial strains were challenged once again with the ICBM5 phage by plaque assay. Dilutions of the phage fraction (10^0^, 10^-2^, 10^-4^ and 10^-6^) were mixed with the host and plated as described above. Incubation was again done at 15 °C, 20°C or 28 °C.

##### ICBM5 purification via CSCl gradient ultracentrifugation

Phages were concentrated from the ICBM5 phage lysate by polyethylene glycol (PEG) precipitation. The soft layer from 20 double agar layer plates with confluent plaques was scraped off, resuspended in SM buffer (100 mM NaCl, 8 mM MgSO4, 50 mM Tris-HCl pH 7.4) and incubated overnight at 4°C. After centrifugation (15 min, 4000 x g, 4°C), the 75 ml supernatant were collected and incubated for 2 h at 4°C with PEG (final concentration 10%) and NaCl (final concentration 0.6 mM). After centrifugation for 2 h at 7197 x g and 4 °C, the supernatant was discarded and the pellet resuspended in 500 µl SM buffer (100 mM NaCl, 8 mM MgSO4, 50 mM Tris-HCl pH 7.4) in total. For resuspension of the phages, 30 min incubation at 4°C followed.

Further concentration and purification of the phages was done by caesium chloride gradient ultracentrifugation. In UltraClear^TM^ centrifuge tubes (Beckman Coulter), a density gradient was set up from caesium chloride solutions with different densities (from bottom up): 1.5 ml of 1.65 g/ml, 2 ml of 1.5 g/ml, 2 ml of 1.4 g/ml, 1 ml of 1.2 g/ml. The PEG concentrated phage fraction was transferred on top. Ultracentrifugation was run for 4 h at 20 °C and 25000 rpm (Beckman, SW 41 Ti). Afterwards, the visible band corresponding to the phages was collected with syringe and needle through the side wall of the ultracentrifuge tube (~500 µl). Removal of caesium chloride was done by dialysis with Slide-A-Lyzer® G2 Dialysis Cassettes 10K MWCO (Thermo Fisher Scientific) against ASW base for 21 h with buffer exchange after 3 h and 18 h. The selected phage band was tested for lysis by spot assay, spotting dilutions (10^0^-10^-11^) of the dialysed band on double agar layer plates containing exponentially growing *Sulfitobacter* sp. SH24-1b and incubating at 20°C.

##### Transmission electron microscopy

30 µl of phage ICBM5 concentrate were pipetted on top of a carbon coated grid (Formvar 162, 200 mesh) and phages were allowed to absorb for 3 min, followed by staining with 30 µl uranylacetate (2 %) for 45 sec and gentle removal of the liquid with filter paper. After air drying for 15 min, the grids were visualized with the transmission electron microscope Zeiss EM902A. Images were documented with the Proscan High Speed SSCCD camera and analyzed using the software ImageSP viewer (Version 1.2.5.16).

##### ICBM5 genome extraction using phenol-chloroform

Concentrated ICBM5 phage from CsCl gradient ultracentrifugation was mixed with the same amount of phenol:chloroform:isoamyl (Roth) solution, gently inverted and centrifuged (15 min, 12.000 x g and 4°C). The aqueous phase was mixed with an equal amount of ice cold absolute ethanol (Th.Geyer) and DNA was precipitated at -80°C for 30 minutes. Another centrifugation was done (20 minutes, 12.000 x g and 4°C) to pellet the phage DNA. Remaining liquid was carefully decanted and the pelleted DNA was resuspended in nuclease free water (Thermo Fisher Scientific). Afterwards, the DNA was purified with NucAway spin column kit (Thermo Fisher Scientific) and quantified via Nanodrop 2000 spectrophotometer.

##### Enzymatic digestions for testing the ssDNA nature of the ICBM5 phage

1 µl of S1 nuclease (Thermo Fisher Scientific), TURBO DNAse (Thermo Fisher Scientific), Exonuclease VII (New England Biolabs) and Hind III (New England Biolabs) were added to separate Eppendorf tubes containing 1 µg of extracted phage DNA. To the one containing TURBO Dnase, 5 µl 10x Dnase buffer was added, the one containing S1 nuclease, 10 µl 5x S1 buffer was added. To the tube with Exo VII, 2.5 µl 1M Tris-HCl pH 8.0 was added. Each tube was filled up to 50 µl with nuclease free water and incubated for 30 min at 37°C. Enzymes were deactivated by increasing the temperature to 95°C for 10 minutes.
2 µl of digest product were mixed with 5 µl of loading buffer (BlueJuice Gel Loading Buffer, Thermo Fisher Scientific) and loaded to a 0.9% agarose gel. The gel was run for 30 min at 80 V and pre-stained with SYBR gold (Thermo Fisher Scientific). The stained gel was analyzed with FAS Digi Gel Documentation System (NIPPON Genetics Europe) and evaluated using BioDocAnalyze software (Biometra GmbH).

ICBM5 concentration using AMICON

A total volume of 15 ml phage stock solution, which was collected from plaque assays, was carefully added into the AMICON ultracentrifugal filter columns (Merck Millipore). During this process, the same column has been used repeatedly. The upper section of the column was filled and centrifuged (5 min, 1.000 x g and 20°C). This was followed by another centrifugation step (12 min, 2.900 x g and 20°C). The concentrate was collected afterwards using a sterile pipette and filtered through a 0.2 µm syringe filter. The filtrate was stored at 4°C in sterile Eppendorf tubes until further processing.

##### ICBM5 purification via OptiPrep™ gradient ultracentrifugation

Phages were purified using an OptiPrep™ density gradient kit (Sigma Aldrich). 1 ml of each concentration (10%, 15%, 20%, 25%, 30%, 35%, 40%, 45% and 50%) was carefully dispensed into two sterile glass gradient tubes. Then, tubes were placed to settle at RT for 2h. 1 ml of the concentrated phage solution was slowly added into the tubes and sub sequentially treated by ultracentrifugation (12 h, 40.000 x g and 20°C) (Beckman, SW 41 Ti). 1 ml of each gradient was collected into sterile 1.5 ml Eppendorf tubes and subjected to spot assays using two different dilutions (10^0^ and 10^-2^). Phages have been found in the 30%, 35% and 40% gradients. Those have been washed and concentrated via AMICON (Merck Millipore). Briefly, 3 AMICON ultra 0.5 ml centrifugal units were filled with 400 µl SM buffer (100 mM NaCl, 8 mM MgSO4, 50 mM Tris-HCl pH 7.4) and 100 µl phage gradient solution and then centrifuged (15 min, 1.000 x g and 4°C). The washing procedure was repeated three times to clean the samples from the gradient solution. 100 µl of freshly washed and concentrated phage solution was collected into sterile 1.5 ml Eppendorf tubes and stored at 4°C

##### ICBM5 genome extraction using the Charge Switch kit

Extracellular DNA was removed by incubating the phage concentrates with 0.043 units/µl of Turbo DNase (Thermo Fisher Scientific) for 30 min at 37°C, followed by enzyme inactivation by incubating for 10 min at 75 °C with 15 mM EDTA. Further, the phage DNA was extracted using the ChargeSwitch gDNA Mini Bacteria Kit (Thermo Fisher Scientific), according to the instructions manual. The first step included only RNAse, no lysozyme treatment. The lysis step with Proteinase K was performed for 30 min at 55 °C. The DNA was finally eluted in 200 µl elution buffer, incubating for 5 min at 55 °C for a higher yield. The concentration and quality of the obtained DNA was checked fluorometrically with Qubit 2.0 and the Qubit® dsDNA HS Assay, spectrophotometrically with Nanodrop 2000 spectrophotometer and visually by regular gel electrophoresis (0.7 % agarose gel, 50 V, SYBR Gold staining).

##### Isolation of a host strain lysogenic for ICBM5 phage – phage ICBM5 specific PCR

ICBM5 specific primers, 304 F, R and 3092 F, R were used. A master mix for PCR was prepared according to (Table S2). DNA of *Sulfitobacter* sp. strain SH24-1b was used as a negative control. Phage ICBM5 DNA was used as positive control.

Extracted DNA was mixed with the master mix 2 µl:48 µl for a final volume of 50 µl. The PCR-cycle (Eppendorf Mastercycler pro S) was 4 min at 96°C, 32x 1 min at 94°C, 30 sec at 55°C and 30 sec at 72°C. Cycling ended with a 10 min period at 72°C and cooled down to 4°C until further downstream processing. Amplified DNA was mixed 5 µl : 2 µl with loading buffer (BlueJuice Gel Loading Buffer, Thermo Fisher Scientific) and loaded into a 1.5% agarose gel. The gel was run in an electrophoresis chamber for 30 min at 85V. Gels were stained using Midori staining solution.

##### Genome of original and infected host strain SH24-1b via PacBio sequencing

Genomic DNA was extracted with the Genomic‐tip 100/G kit (Qiagen, Hilden, Germany). A SMRTbell™ (PacificBiosciences, Menlo Park, CA, USA) template library was prepared according to the manufacturer’s “Procedure & Checklist - 20 kb Template Preparation Using BluePippin™ Size-Selection System” protocol. Shortly, sheared genomic DNA was end-repaired and ligated to hairpin adapters applying components from the DNA/Polymerase Binding Kit P6 (Pacific Biosciences, Menlo Park, CA, USA). BluePippin™ Size-Selection to 7 kb was performed as recommended by the manufacturer (Sage Science, Beverly, MA, USA). SMRT sequencing was carried out on the PacBio RSII (Pacific Biosciences) taking 240-minutes movies, which resulted in 166,457 and 90,339 post-filtered reads with a mean read length of 14,035 bp and 14,014 bp, respectively. Illumina libraries were prepared with the Nextera XT DNA Sample Preparation Kit (lllumina Inc., San Diego, CA, USA) modified after (Baym *et al.* 2015) and paired-end sequencing was performed on the NextSeq 500 (PE75).

Long read genome assembly was performed with the “RS_HGAP_Assembly.3“ protocol in SMRTPortal version 2.3.0. The assembled contigs were error-corrected by mapping of Illumina short reads using the Burrows-Wheeler Aligner (BWA 0.6.2) (Li and Durbin 2009) and subsequent variant and consensus calling using VarScan 2.3.6 (Koboldt *et al.* 2012). The final assembly was trimmed, circularized and adjusted to the replication system as start point (https://github.com/boykebunk/genomefinish) and checked via the mapping of Illumina reads (BWA) and PacBio reads (Bridgemapper). The genome was annotated with Prokka 1.13 (Seemann 2014) with subsequent manual curation of the replication systems. For the infected strain, PacBio reads were not only assembled but also mapped on the original genome including the phage ICBM5 genome. PacBio reads were also compared with blastn against the genome of phage ICBM5 but no hit was detected.

##### Sequencing of Ascunsovirus oldenburgi ICBM5 via Illumina sequencing

Exact concentration of double stranded viral DNA was determined using the Qubit® dsDNA HS Assay Kit as recommended by the manufacturer (Life Technologies GmbH, Darmstadt, Germany). Illumina shotgun libraries were prepared using the Nextera XT DNA Sample Preparation Kit (Illumina, San Diego, CA, USA). To assess quality and size of the libraries, the samples was run on an Agilent Bioanalyzer 2100 using an Agilent High Sensitivity DNA Kit as recommended by the manufacturer (Agilent Technologies, Waldbronn, Germany). Concentration of the libraries were determined using the Qubit® dsDNA HS Assay Kit as recommended by the manufacturer (Life Technologies GmbH, Darmstadt, Germany).Sequencing was performed on a MiSeq system with the reagent kit v3 with 600 cycles (Illumina, San Diego, CA, USA) as recommended by the manufacturer resulting in 785.119 paired end reads.

##### Sequencing of the host strain lysogenic for ICBM5 phage via Nanopore

Phenol chloroform extracted DNA was checked for purity via Nanodrop before library preparation was done. Absorbance ratio of 260/280 has to be between 1.8 and 2.0, and absorbance ratio of 260/230 has to be between 2.0 and 2.2 to make sure that the pores are not clocked during the run. Once purity had been validated, library preparation was performed as described by the manufacturers (Oxford Nanopore Technologies, SQK-RBK004). 7.5 µl of host DNA was combined with 2.5 µl fragmentation (FRA) mix in a small PCR LoBind tube (Eppendorf, Germany) and incubated for 1 min at 30°C and then for another 1 min at 80°C. Afterwards the tube was spun down and stored on ice, 1 µl of rapid mix (RAP) was added and the solution was incubated for 5 minutes at room temperature. The prepared library was stored on ice. To perform the sequencing run, the flow cell needed to be primed. Therefore, 30 µl of flushing tether (FLT) was mixed with the flushing buffer (FLB). The flow cell was placed into the MinIon sequencer and the priming port was opened. The pipette tip was placed on top of the port and a small amount of liquid was sucked in. Afterwards, 800 µl of the prepared priming mix (FLB and FLT) were slowly loaded into the priming port while taking care to not introduce air bubbles. Then, the flow cell needed to incubate for 5 min. Next, the pre-sequencing mix was prepared. 34 µl of sequencing buffer (SEQ) was mixed with 4.5 µl nuclease free water (Thermo Fisher Scientific). The loading beads (LB) were gently resuspended and added together with 11 µl of the DNA library to the diluted sequencing buffer. The solution was gently mixed and spun down to make sure everything is at the bottom of the tube. To load the flow cell, the sample port was opened and 200 µl of the priming mix were added slowly. The pre-sequencing mix was resuspended again and loaded drop by drop into the sample port. All ports were closed and the run could be started. Therefore, the MinIon software was started and ensured that the device was connected and found within. A ‘new experiment’ was selected and the default parameters have been used. Base calling was performed simultaneously during sequencing using the MinIon software.

After sequencing was done, all .fastq files were merged (cat *.fastq > 01_reads.fastq). First quality control was done by FastQC (https://www.bioinformatics.babraham.ac.uk/projects/fastqc/) (fastqc -t 16 $input"01_reads.fastq" -o $Step2_out --nano) and adapters were removed using PoreChop (https://github.com/rrwick/Porechop) (porechop -i $input"01_reads.fastq" -t 16 -v 2 -o $output"03_reads_trimmed.fastq" > $output"03_porechop.log"). Assembly was performed using a set of tools from the pomoxis suite (https://github.com/nanoporetech/pomoxis). Minimap2 (minimap2 -x ava-ont -t $threads $output"01_reads.fastq" $output"01_reads.fastq" | gzip -1 > $mapping"08_mapping.paf.gz") and miniasm (miniasm -f $output"01_reads.fastq" $mapping"08_mapping.paf.gz" > $mapping"08_miniasm_reads.gfa") were used to map the .fastq files onto each other to find overlaps. Afterwards, the .gfa file was converted to .fasta (awk '$1 ~/S/ {print ">"$2"\n"$3}' $mapping"08_miniasm_reads.gfa" > $assembly"09_miniasm_reads.fasta"). Assembly was done by minimap2 (minimap2 $assembly"09_miniasm_reads.fasta" $output"01_reads.fastq" > $assembly"09_minimap_reads.paf") and polishing was done by racon (racon -t $threads $output"01_reads.fastq" $assembly"09_minimap_reads.paf" $assembly"09_miniasm_reads.fasta" > $polished"10_racon.fasta"). A last quality control was performed by quast (quast.py -t $threads -o $Step10_QC $polished"10_racon.fasta" $assembly"09_miniasm_reads.fasta") (Gurevich *et al.* 2013).

##### One step infection experiment with ICBM5 phage

For one-step infection curves, the ICBM5 phage was added at a multiplicity of infection (MOI) of 6.5 to a culture of *Sulfitobacter* SH24-1b growing exponentially in MB media. To enable the MOI calculation, the phage and bacteria concentrations were determined as follows.

The concentration of the phage stock was determined in the same day with the one step infection experiment, by performing plaque assays for serial dilutions of the phage stock. The following formula was applied to calculate number of plaque forming units (PFUs) per ml:

PFU/ml = number of single plaques per plate / (dilution x volume of phage stock dilution in ml)

The concentration of the bacterial cells was determined right before setting up the one step infection experiment. For this, the optical density at 600 nm was measured and then correlated with the corresponding number of cells per ml, as previously determined by plating and counting of colony forming units (CFUs), see Fig. S1.

Through the infection experiment, plaques assays (see section “Phage plaque assays using the double-agar layer method“) were used to quantify both the free and total (free and cell bound) phages. The free fraction was obtained by filtering the infected culture through a 0.22 µm syringe filter.
For phage-targeted genomeFISH, to the culture samples paraformaldehyde (Electron Microscopy Sciences) was added to a final concentration of 4%. The samples were incubated at RT for 1 h. Afterwards, the fixed cells were pelleted by centrifugation at 5.000 x g for 10 minutes, resuspended in 1 x PBS (137 mM NaCl, 2.7 mM KCl, 8 mM Na_2_HPO_4_ and 2 mM KH_2_PO_4_) (Invitrogen) and again pelleted by centrifugation at 5.000 x g for 10 minutes. The supernatant was discarded and the cell pellet was resuspended in a 1:1 ratio in 1 x PBS and absolute ethanol (Th. Geyer). The fixed cells were then stored at -20°C until further processing.

##### Calculating burst size from PFU counts

To calculate the burst size from the one step growth infection cycle experiment, first the amount of free phage average of the time points on the plateau before the burst was determined (A). Then the free phage average of the time points on the plateau after the burst was determined (B). Afterwards, A was subtracted from B, which gives the total burst or new phages released (C). This number is divided by the number of infecting phages (free phage average at T1) which gives the final burst size.

T1: 544.500

A: 3.321.666.67

B: 139.600.000

C: (139.600.000 - 3.321.666.67) = 136.278.333

Burst size: 136.278.333 / 544.500 = **250.28**

##### Labeling of ICBM5 genome probes

The dry dsDNA polynucleotides delivered by Integrated DNA Technologies (IDT) were resuspended in 5 µl labelling buffer (5 mM Tris-HCl and 1 mM EDTA), to a final concentration of 100 ng/µl. 2 µl of each probe were collect in a 0.2 ml thin walled PCR tube (Thermo Fisher Scientific) and filled up to 20 µl with labelling buffer. Further, probes have been denatured at 95°C for 5 minutes and placed on ice. Immediately afterwards, 30 µl of Alexa Flour 594 labeling reagent (Invitrogen) was added while still on ice and then incubated at 80°C for 30 minutes. The probes were purified using the NucAway Spin column kit (Thermo Fisher Scientific). The concentration and labeling efficiency were measured using a Nanodrop 2000 spectrophotometer. Base to dye ratios were calculated according to the instructions given with the ULYSIS Nucleic Acid Labeling Kit (Thermo Fisher Scientific). 1 dye every 20 bases has been calculated and 22 ng/µl DNA concentration was quantified.

##### Image Analysis

Cells that have been visualized via microscopy and were quantified using CellProfiler Image analysis software (version 3.1.9). An automated quantification pipeline was developed which works as follows: every picture which needed to be analyzed was imported into the program and designated to different groups (e.g. DAPI pictures to DAPI group and Alexa pictures to Alexa group). For this, the lowest exposure settings were used, to allow for the detection of single phage signals and avoid analyzing pictures that have been overexposed. In the first step of the pipeline, primary objects, in this case bacterial cells, were identified within the DAPI group by the algorithm of the software and edited manually, so all cells in one picture were selected. In the second step, an empty spot was selected manually within the Alexa group and defined as background. This background was then subtracted from the Alexa image. The data for every analyzed picture was exported as an excel sheet, containing the image number, cell number, area size for cells and the mean intensity for each area after background correction. Within the negative control images (samples without an infection), the mean signal intensity was calculated and used to produce a new column, the corrected intensity in which negative control mean signal intensity was subtracted from the mean signal intensity after background correction. This new column was multiplied with the area size to determine the total intensity for each cell.

To quantify the fraction of infected cells, false positives have been deleted from the dataset by removing cells with a strong adjacent signal and those that occur in dense cell clusters. Within these reduced datasets, infected cells have been quantified manually. The amount of infected cells per time point was then divided by the total amount of cells within the reduced dataset to get the percentage of infected cells.

To calculate the amount of phage genome copy per cell an average value for one phage genome signal was determined. This was done by selecting 100 single phage signals at low exposure times from T1 and calculating an average value. This value was multiplied with the total intensity determined from previous steps to determine the amount of phage genome copies for each cell in each picture and each time point, allowing for a precise quantification of infected cells.

#### Results

Tab S1: PCR reagent master mix composition.

| µl/50 µl end volume | Reagent | Final concentration |
| --- | --- | --- |
| 5 | 10x Rxn Buffer |  |
| 5 | dNTPs (2.5 mM) | 250 µM |
| 2.1 | MgCl_2_ (50 mM) | 2.1 mM |
| 2.5 | BSA (30 mg/ml) | 1.5 mg/ml |
| 32.2 | PCR H_2_O |  |
| 0.5 | Primer F (20 pmol/µl) | 10 pM |
| 0.5 | Primer R (20 pmol/µl) | 10 pM |
| 0.2 | Taq Polymerase (5U/µl) | 1U |

Table S2: List of *Rhodobacteraceae* strains used for the host range assay.

| **Name** | **Strain designation** | **Strain** | **Infected (- no, + yes)** |
| --- | --- | --- | --- |
| *Aliiroseovarius crassostreae* | CV919-312, CVSP | DSM 16950^T^ | - |
| *Aliiroseovarius halocynthiae* | MA1-10 | DSM 27840^T^ | - |
| *Antarctobacter heliothermus* | EL-219 | DSM 11445^T^ | - |
| *Celeribacter baekdonensis* | L-6 | DSM 27375^T^ | - |
| *Celeribacter halophila* | ZXM137 | DSM 26270^T^ | - |
| *Celeribacter indicus* | P73 | DSM 27257^T^ | - |
| *Celeribacter marinus* | IMCC12053 | DSM 100036^T^ | - |
| *Celeribacter neptunius* | H 14 | DSM 26471^T^ | - |
| *Cognatishimia maritimus* | GSW-M6 | DSM 28223^T^ | - |
| *Cognatiyoonia koreensis* | GA2-M3 | DSM 17925^T^ | - |
| *Dinoroseobacter shibae* | DFL 12 | DSM 16493^T^ | - |
| *Hwanghaeicola aestuarii* | Y26 | DSM 22009^T^ | - |
| *Jannaschia donghaensis* | DSW-17 | DSM 102233^T^ | - |
| *Jannaschia helgolandensis* | Hel10 | DSM 14858^T^ | - |
| *Jannaschia pohangensis* | H1-M8 | DSM 19073^T^ | - |
| *Jannaschia rubra* | 4SM3 | DSM 16279^T^ | - |
| *Leisingera aquimarina* | R-26159 | DSM 24565^T^ | - |
| *Leisingera caerulea* | 13 | DSM 24564^T^ | - |
| *Leisingera daeponensis* | TF-218 | DSM 23529^T^ | - |
| *Leisingera methylohalidivorans* | MB2 | DSM 14336^T^ | - |
| *Limimaricola cinnabarinus* | LL-001 | DSM 29954^T^ | - |
| *Limimaricola hongkongensis* | UST950701-009P | DSM 17492^T^ | - |
| *Limimaricola pyoseonensis* | JJM85 | DSM 21424^T^ | - |
| *Litoreibacter albidus* | KMM 3851 | DSM 26922^T^ | - |
| *Litoreibacter arenae* | GA2-M15 | DSM 19593^T^ | - |
| *Litoreibacter janthinus* | KMM 3842 | DSM 26921^T^ | - |
| *Loktanella fryxellensis* | R-7670 | DSM 16213^T^ | - |
| *Loktanella salsilacus* | R-8904 | DSM 16199^T^ | - |
| *Maribius pelagius* | B5-6 | DSM 26893^T^ | - |
| *Maribius salinus* | CL-SP27 | DSM 26892^T^ | - |
| *Marinovum algicola* | FF3 | DSM 10251^T^ | - |
| *Marinovum algicola* | DG898 | DSM 27768 | - |
| *Maritimibacter alkaliphilus* | HTCC2654 | DSM 100037^T^ | - |
| *Oceanicola granulosus* | HTCC2516 | DSM 15982^T^ | - |
| *Octadecabacter temperatus* | SB1 | DSM 26878^T^ | - |
| *Pacificibacter marinus* | HDW-9 | DSM 25228^T^ | - |
| *Palleronia marisminoris* | B33 | DSM 26347^T^ | - |
| *Phaeobacter gallaeciensis* | BS 107 | DSM 26640^T^ | - |
| *Phaeobacter inhibens* |  | DSM 17395 | - |
| *Phaeobacter inhibens* | T5 | DSM 16374^T^ | - |
| *Phaeobacter inhibens* | 2.10 | DSM 24588 | - |
| *Phaeobacter italicus* | R11 | DSM 26436^T^ | - |
| *Ponticoccus litoralis* | CL-GR66 | DSM 18986^T^ | - |
| *Pseudooceanicola batsensis* | HTCC2597 | DSM 15984^T^ | - |
| *Pseudooceanicola nanhaiensis* | SS011B1-20 | DSM 18065^T^ | - |
| *Pseudophaeobacter arcticus* | 20188 | DSM 23566^T^ | - |
| *Pseudoruegeria lutimaris* | HD-43 | DSM 25294^T^ | - |
| *Roseibacterium elongatum* | Och 323 | DSM 19469^T^ | - |
| *Roseivivax isoporae* | sw2 | DSM 22223^T^ | - |
| *Roseobacter denitrificans* | Och 114 | DSM 7001^T^ | - |
| *Roseobacter litoralis* | Och 149 | DSM 6996^T^ | - |
| *Roseovarius indicus* | B108 | DSM 26383^T^ | - |
| *Roseovarius lutimaris* | 112 | DSM 28463^T^ | - |
| *Roseovarius mucosus* | DFL-24 | DSM 17069^T^ | - |
| *Roseovarius nubinhibens* | ISM | DSM 15170^T^ | - |
| *Ruegeria atlantica* | 1480 | DSM 5823^T^ | - |
| *Ruegeria conchae* | TW15 | DSM 29317^T^ | - |
| *Ruegeria marina* | ZH17 | DSM 24837^T^ | - |
| *Ruegeria pomeroyi* | DSS-3 | DSM 15171^T^ | - |
| *Sagittula stellata* | EE-37 | DSM 11524^T^ | - |
| *Salinihabitans flavidus* | ISL-46 | DSM 27842^T^ | - |
| *Salipiger bermudensis* | HTCC2601 | DSM 26914^T^ | - |
| *Salipiger aestuarii* | AD8 | DSM 22011^T^ | - |
| *Salipiger marinus* | CK-I3-6 | DSM 26424^T^ | - |
| *Salipiger mucosus* | A3 | DSM 16094^T^ | - |
| *Salipiger pacificus* | DX5-10 | DSM 26894^T^ | - |
| *Sedimentitalea nanhaiensis* | NH52F | DSM 24252^T^ | - |
| *Sediminimonas qiaohouensis* | YIM B024 | DSM 21189^T^ | - |
| *Shimia aestuarii* | JC2049 | DSM 15283^T^ | - |
| *Shimia haliotis* | WM35 | DSM 28453^T^ | - |
| *Shimia marina* | CL-TA03 | DSM 26895^T^ | - |
| *Sulfitobacter delicatus* | KMM 3584 | DSM 16477^T^ | - |
| *Sulfitobacter dubius* | KMM 3554 | DSM 16472^T^ | + |
| *Sulfitobacter indolifex* | HEL-45 | DSM 14862^T^ | - |
| *Sulfitobacter litoralis* | Iso 3 | DSM 17584^T^ | - |
| *Sulfitobacter marinus* | SW-265 | DSM 23422^T^ | - |
| *Sulfitobacter mediterraneus* | CH-B427 | DSM 12244^T^ | - |
| *Sulfitobacter noctilucae* | NB-68 | DSM 100978^T^ | - |
| *Sulfitobacter noctilucicola* | NB-77 | DSM 101015^T^ | - |
| *Sulfitobacter pseudonitzschiae* | H3 | DSM 26824^T^ | - |
| *Sulfitobacter sp.* | EE-36 | DSM 11700 | - |
| *Sulfitobacter dubius* | SH24-1b |  | + |
| *Thalassobius taeanensis* | G4 | DSM 22007^T^ | - |
| *Thalassococcus halodurans* | UST050418-052 | DSM 26915^T^ | - |
| *Thioclava dalianensis* | DLFJ1-1 | DSM 29618^T^ | - |
| *Thioclava pacifica* | TL 2 | DSM 10166^T^ | - |
| *Tranquillimonas alkanivorans* | A34 | DSM 19547^T^ | - |
| *Tranquillimonas rosea* | BH87090 | DSM 23042^T^ | - |
| *Tritonibacter multivorans* | MD5 | DSM 26470^T^ | - |
| *Tropicibacter naphthalenivorans* | C02 | DSM 19561^T^ | - |
| *Tropicimonas isoalkanivorans* | B51 | DSM 19548^T^ | - |
| *Wenxinia marina* | HY34 | DSM 24838^T^ | - |
| *Yoonia tamlensis* | SSW-35 | DSM 26879^T^ | - |
| *Yoonia vestfoldensis* | R-9477 | DSM 16212^T^ | - |

Tab. S3: Overview of designed polynucleotides showing names, sequences, start and end position as well as GC content

| **names** | **sequences** | **start** | **end** | **length** | **GC%** |
| --- | --- | --- | --- | --- | --- |
| Poly_11 | aggatgcgcgacgataaaattagctcttgcaaaccccgtccaaattcgactaaggtcaatttagggaatggttcccggcgcgggtttatcccgtggtgctaacaaaggaaaaacacatggctgataaagcaacttccccgactaccgtctctaaaaaagatcgtgatgattttgagcttcatgttcggctcttgagccgggcaggttacaaaactgccgaagctcggacaattgcatggctgcgcggccctgctgggctgcaagaaatgcttgcgggcgtcaaggtcgcaagctaacata | 11 | 310 | 300 | 50 |
| Poly_1891 | tcgaaggcgtcggcgttgaatacaatgccgttccagtgactacccccggttggaacgatacagagaatgaccgtgctgctgatcctcactatttctcggaccacgccgcaggcattcgccttgcgattgcctcggagaatggtcgtccagcggtctacgctgttggcgagggtcaaaatgccggcggcatttcccttactgacttctacaacgccgaactgatggacagtcttgtgcgccagatgcggcagattgtcgatgataaccctgagtatggcgaagaaatggtcacacgctggg | 1891 | 2190 | 300 | 55 |
| Poly_2229 | attttgcaccaaagccagcaaatgttcggcaatcaattccgccgtgccatggatggtgcaaacctcgatgttgctcaatccgacatgatgcaaagccttgaattcactgtccccgtgcctccaacggaattgggcggggtcgtgattacctttgcatctgtgaagcctgacgaaacactaggctctcagccgcacccgttcctgtcgacaacatgggaggcgactaactacgtctccgacgagttggcacgcgaccctgaggccgtgaccatgcgtcacctgaacagtgacgttcctatg | 2229 | 2528 | 300 | 54 |
| Poly_2538 | gacactcgtgctctctacatcggtcacaacggtctgaaaaaagcctacatctcatatggctttaaccggcatctcgaccctactacggtcgaagcgaaaactgccatctggcagcttgaggtgccaatgtccgtaacgcctgaaagcgttatctatcccgaggacttggaccattatccgtttgcggatcagcttgcagaggtctgcacctatcaggttagttcgaccgctacagtgcggacgcctatggtcttcggtccgacaccagttgaggaactggcccagattgagacagacaac | 2538 | 2837 | 300 | 52 |
| Poly_2868 | ggttatgccgcccctcaataaaccttaaaaaactggaaagatgaaatgcaagttaatgatgcaaaacgctccgttcttctcgtcggccttaatggcggcgaagtttctgtaatctccgcagatggtgagattatcgcgaccgaaggtgtaaccgctggtcggcataaatgctcctcttgggtgccatttatgtccaatgagggcgatgaattgagcttctccggcgatgtggtcccaatggtgccaaatggcggtcgtgtccggcctatggcctacggccccggtcaatttgaaagcggt | 2868 | 3167 | 300 | 51 |
| Poly_3876 | ctatagacctcgtgcatggttatgattggggatgggacagtgcccatattgagattgctttgtggcgtcaagttccaaagcgtcaaattggcaaaactggctctaattatcccccgtcaccttgggaacgtgaacagcgtttcaaggagttgcttcccgctgtttatgcggcgcaacatcggccatagacttatggcaatctatgcaaactaacaagcgggcggcctctatgcgagagccgcccgttttgcatccctgcgggccagtcggagactggccagcggcaactatattccgggc | 3876 | 4175 | 300 | 53 |
| Poly_4400 | tttgcctgtcgtaagtgtaatgagtgcatcaccgctaggaaaaatggttgggttgctcgtgcgatggctgaaaaggccgtcactgctgaaactttcagcgtaaccctaacttataatgatgctactcaggaaagccgcgatgcggcaaagacgttcgagtatcgacacgtcaaaaactggattaagaaccttaggcgtcaaatagagtacaccaccggccagactggccttctccgctaccttgtagcgggtgagcgcggttccgataagggccgctgccactggcatgtaatcttgttc | 4400 | 4699 | 300 | 51 |
| Poly_4700 | Tgtaacgccgatattttaacgctaggaaaaatgacgcactggccctctggcaaacttgctgccgatagggccgaaattataactacaggaaaaaggaaaaaacgtattaactggtctctctggccgtatggcttcgtcagctttcaagagcctgaccagtggggcatggagtacgccttagcgtatgccctcaaagatcagtttaacattgtctccgctgccggtacggcgcgggaggctcacgtttcccgcacttctgcgggtatgttccgcatgtccaaaaaacccccaatcggtttc | 4700 | 4999 | 300 | 51 |

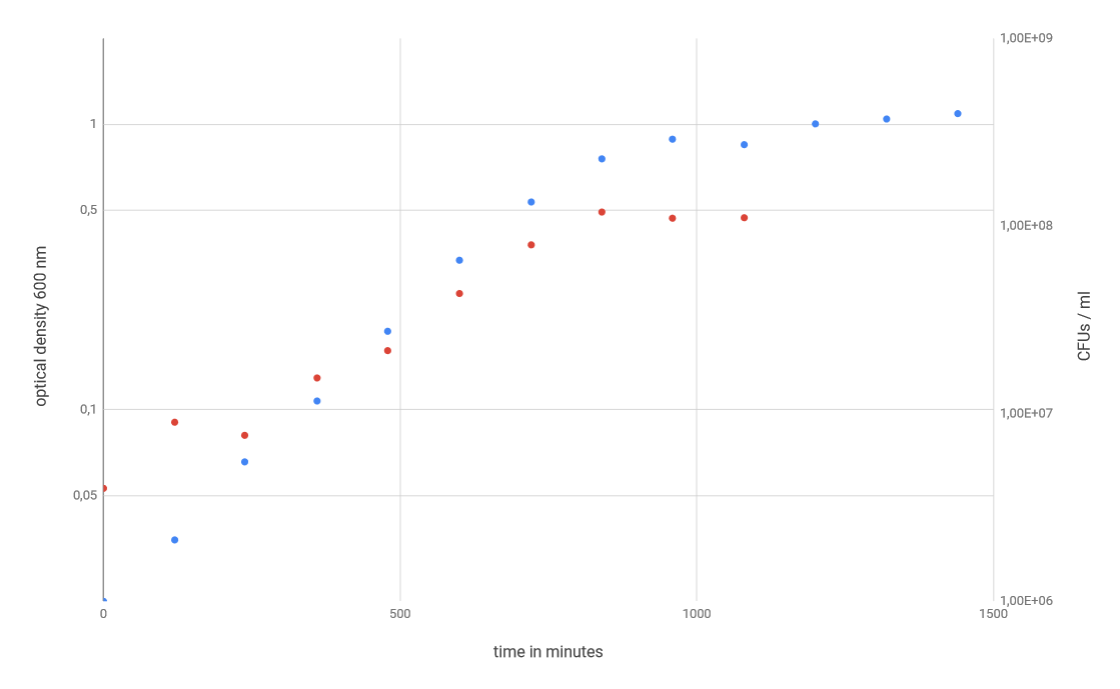

Fig. S1: CFU assay of *Sulfitobacter* sp. SH24-1b, blue dots representing the optical density values over time, while the red dots show the development of CFUs/ml over time. An OD_600_ of 0.5 roughly equals 1*10^8^ CFUs/ml.

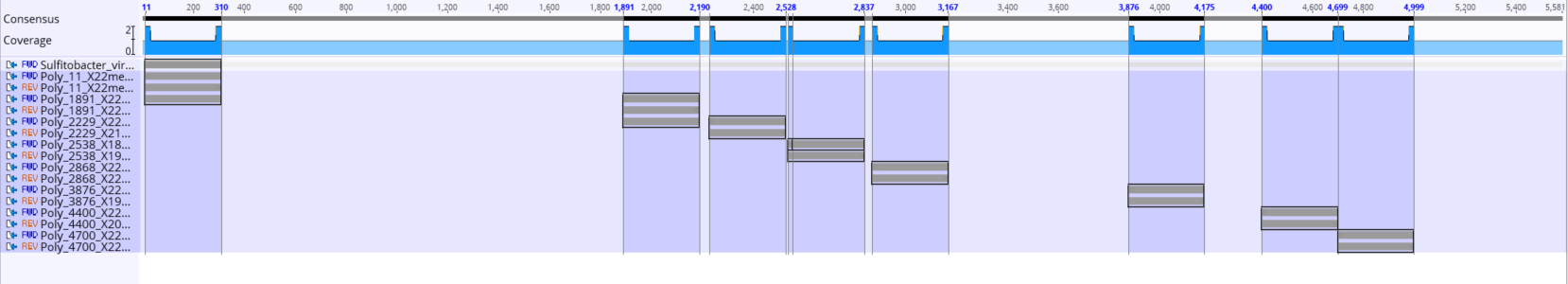

Fig. S2: Overview of designed polynucleotides on the entire genome of ICBM5 phage, the selected regions are each 300 bps in length.

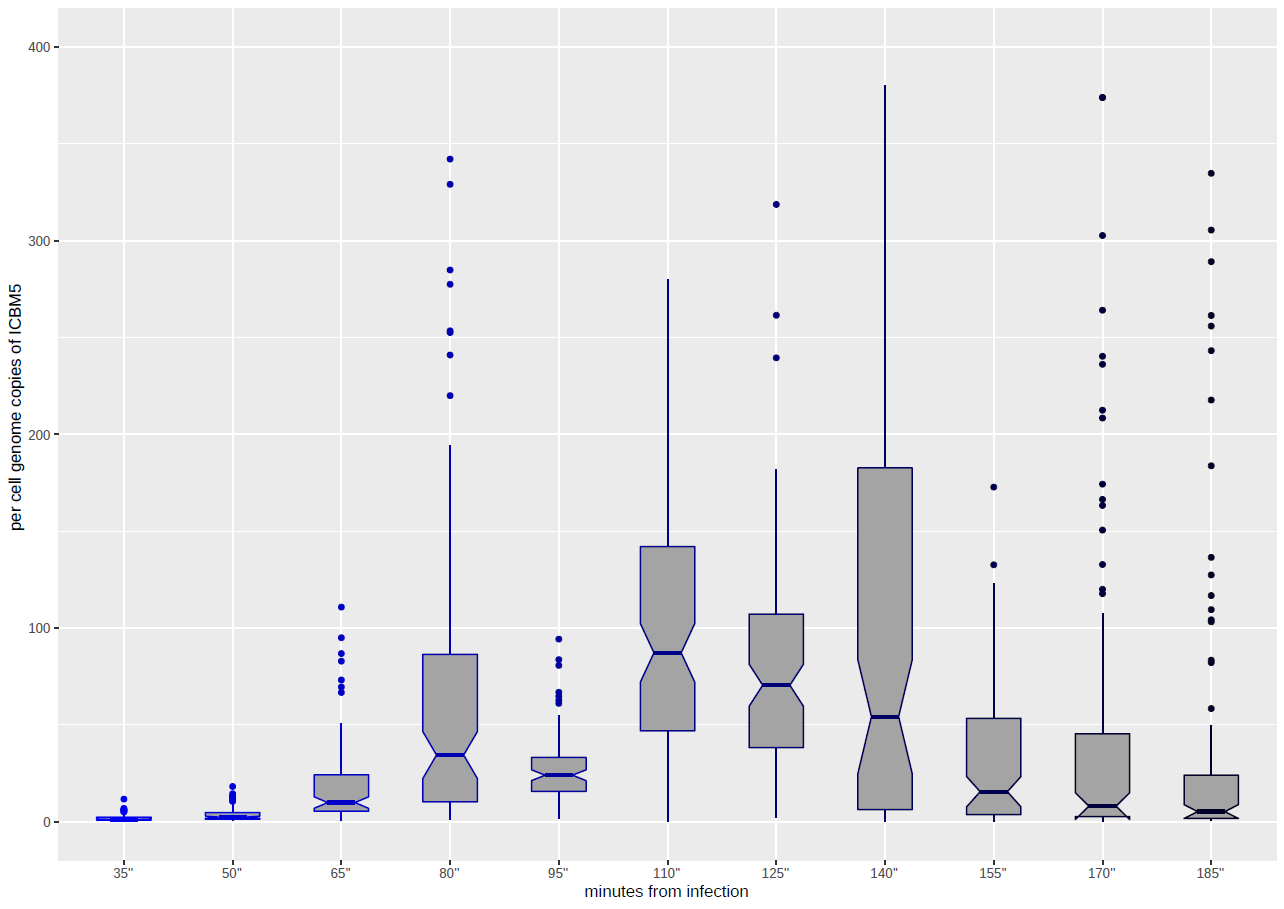

Fig. S3: The per cell genome copies of ICBM5 through the infection, presented as box plot. The box plot borders represent the 25th and the 75th percentile, and the middle line represents the 50th percentile. The plot was generated using the ggplot2 R package v. 3.3.0

Genomic information

>Ascunsovirus_oldenburgi_ICBM5
TTCAGACGAAAGGATGCGCGACGATAAAATTAGCTCTTGCAAACCCCGTCCAAATTCGACTAAGGTCAATTTAGGGAATGGTTCCCGGCGCGGGTTTATCCCGTGGTGCTAACAAAGGAAAAACACATGGCTGATAAAGCAACTTCCCCGACTACCGTCTCTAAAAAAGATCGTGATGATTTTGAGCTTCATGTTCGGCTCTTGAGCCGGGCAGGTTACAAAACTGCCGAAGCTCGGACAATTGCATGGCTGCGCGGCCCTGCTGGGCTGCAAGAAATGCTTGCGGGCGTCAAGGTCGCAAGCTAACATAGGTTGAGCGGTTCGCCGCTCAACATTAACCGCAAAAAAGGAGAGAAATGTGAGTGCATTGAAACACTTTCTTACAAACCGTTTGACAGTCGTGGTTCTTACCTCGCTGGCAACGGCAGTAGGTACTGCGCTTGCGACGGAGTTTCCGTCCATTTATAGCGCAGTCTGTGCCTAATGTCGGCTATCGCTGGTGCTCTTATCTCAGGTGGCGCAAGCCTCCTAGGTGGTCTGTTCGGGCGTTCTTCTGCTAGCAAGCAGCAAGCCCGACAGAATGAGTACAACAAGCCGATTAACATTCGCAAGCGGGCCGAGGAGGGGGGGTTCAACCCCCTGCTCTGGGCCGGTCAAGGCAACATCCAAATGCAGCCGGGTCCGTCCGGCATCATGGGTTCCGCTATTGCGAATGCTGGCCTAGCTCTTGCCGATGGCATGAGCGAACAACGCCAGCTCGACCTTGAGCGTACCAAGCTCAAGCAAGATCAAGAGCGTCTCGACGCTCTGATCGAAAAACAGACCATCCGGCCAAAGGTCGGCGGCATTTATGCCGGGTCGCAACAAACGCCTTCTGTAGCGCGCGCTCCCGGTCGCCCGCTTATGAATGGCGCTCCTCAACCCGGCTCTGCGCCGGTCTTTAACCCGCCAACGGAGTACAACCCAATCCCCGATGATGGCCCTCGCCTGCAAACGAAAGTGATGCGTAGCGATGGCATGACCTCGGCGGATCCTGAAAATCCCGCCGAAATGGAGGGCGATTGGTGGACGTGGGCCAGAGAGGGAACTTTCTGGCAAAACAACAACGAAATTCTGCGGCGTAATACGCCGGAGACGTTGCACTACAAGGGTCGCGATGCCTTGTTCCCGAAAATGATTGACGGGGCAAGGAAAGCGCATAAGAAGGCTCAAGAGGACTTCGAGAAAAACCCGCCAAAACGCCGCAAGCTTAAAGGCGTCAACCCTAACCTAAACGACAAGAAATGGTAATGAACATGTCAAAGTATCAACGTCCTACAAACACACGCCGCGAAAGCCGGACCATCGCTGGCCGGTTCCGTGGCGGCAAGTTGGCTCCTGTTATGGCGTCCGCGTTCCGTGAGAGCGAAAGTGCAATCCTTTCGCAACAAGTTACCTATGAACTTGACCCAATCGCGGGCCGTATGATTACGCCGATCATGGCGGAACTTATCTCTGTATATGTTCCGGTCCAAGCGATCGACGCCCTAAAAAACCCTGAGGAGGCTTATGCCGGTAACACTGAGGTTGTCCGTGACAAGCTCCTCTCAGGAACGCCGCTGTTCGGCCTCGAAGACGAAAGCGAGATTTCGAAGCGTCTAGGCGTTAACCCTATCTCTGTGGGTGGTGTCAAAAAAGTGAACGAGGCGGCACGCCTTGCGCATAACTGCGCCGTGAACTTCCTACGTCAGCGCAAGTACGTCAACACGGTCAAGCTATTGGCCGATAATATGAACGTAACGCCTGCGCTGATCTCTCAAACGGTTCTCGACCGGCTGAACGCCGTGCTCGATCCTGAGGATCGTGTCAACGGTGCTGTGCAGCTTGACCTTGGGAACGTGCGTATGCCGGTCGAAGGCGTCGGCGTTGAATACAATGCCGTTCCAGTGACTACCCCCGGTTGGAACGATACAGAGAATGACCGTGCTGCTGATCCTCACTATTTCTCGGACCACGCCGCAGGCATTCGCCTTGCGATTGCCTCGGAGAATGGTCGTCCAGCGGTCTACGCTGTTGGCGAGGGTCAAAATGCCGGCGGCATTTCCCTTACTGACTTCTACAACGCCGAACTGATGGACAGTCTTGTGCGCCAGATGCGGCAGATTGTCGATGATAACCCTGAGTATGGCGAAGAAATGGTCACACGCTGGGCGCATGGTCTTTCTGTTGATAACGGCAAAACTCCTTGGATTTTGCACCAAAGCCAGCAAATGTTCGGCAATCAATTCCGCCGTGCCATGGATGGTGCAAACCTCGATGTTGCTCAATCCGACATGATGCAAAGCCTTGAATTCACTGTCCCCGTGCCTCCAACGGAATTGGGCGGGGTCGTGATTACCTTTGCATCTGTGAAGCCTGACGAAACACTAGGCTCTCAGCCGCACCCGTTCCTGTCGACAACATGGGAGGCGACTAACTACGTCTCCGACGAGTTGGCACGCGACCCTGAGGCCGTGACCATGCGTCACCTGAACAGTGACGTTCCTATGGTCGATGAGGACACTCGTGCTCTCTACATCGGTCACAACGGTCTGAAAAAAGCCTACATCTCATATGGCTTTAACCGGCATCTCGACCCTACTACGGTCGAAGCGAAAACTGCCATCTGGCAGCTTGAGGTGCCAATGTCCGTAACGCCTGAAAGCGTTATCTATCCCGAGGACTTGGACCATTATCCGTTTGCGGATCAGCTTGCAGAGGTCTGCACCTATCAGGTTAGTTCGACCGCTACAGTGCGGACGCCTATGGTCTTCGGTCCGACACCAGTTGAGGAACTGGCCCAGATTGAGACAGACAACGTCTTCGAAGACGTATAAAAAAAGGGGGGCGGTTATGCCGCCCCTCAATAAACCTTAAAAAACTGGAAAGATGAAATGCAAGTTAATGATGCAAAACGCTCCGTTCTTCTCGTCGGCCTTAATGGCGGCGAAGTTTCTGTAATCTCCGCAGATGGTGAGATTATCGCGACCGAAGGTGTAACCGCTGGTCGGCATAAATGCTCCTCTTGGGTGCCATTTATGTCCAATGAGGGCGATGAATTGAGCTTCTCCGGCGATGTGGTCCCAATGGTGCCAAATGGCGGTCGTGTCCGGCCTATGGCCTACGGCCCCGGTCAATTTGAAAGCGGTGCCAATCCCGATTTCGTCGTTACTTCGGCGGATCGGATGGCTCGTGAGCTTGACCATAAAATTAGGGGTCTTGACCAGACCGCTAAAAAAGTTGAGGCCCGTATTGCGCAGTTGAACAATCTAGCAGAACGCGCTAAAACACCTGTAGAAGTAGCAAAGGAGGAAAAAGTCGATGTTATTGATGATGACGATGTTTCTGACGTTCGTAGCTCTGGCGACGATAGTGACGATACGGGACCTGATTTTGGCGCTGAAAAAGTGGGGGCAGAATGAATGATGCCCCGAAAAAAAGCAATTCGGGTCGCAAACAAAGCTGGCTCCGCGCGTATGCGCGGGGCCTCAAAGGCGGCACCCCGAAAACGCTCTGCGAAAGCCTGCGCCTGTGGTCAATCCCACAAGAGCAGGCGCAAGACGTGACAAAAAAAACTTACCTCAACGCCGCAGATGGCCCCATGATCTGGCTCGGTGAGCAGATTGTCAAACGCATGGAAAAGGCGGGTTATCCCTCCCGCATTTTCTGCGGCTATCGTTCCCCAGAACAGCAAGACAAGGAGTTTGCAGAGGGCGACAGCAAGGCTAGGGCCTATCAAAGCCCACATCAGTTTTATGAAGCGGTAGACATAATTCATAAGACAAAAGCATGGAACGTCTCTCAAGATTATTGGGATACACTCGCCGCAATTGTGCGAGTTGTAGAGCGTGAATATGCTATAGACCTCGTGCATGGTTATGATTGGGGATGGGACAGTGCCCATATTGAGATTGCTTTGTGGCGTCAAGTTCCAAAGCGTCAAATTGGCAAAACTGGCTCTAATTATCCCCCGTCACCTTGGGAACGTGAACAGCGTTTCAAGGAGTTGCTTCCCGCTGTTTATGCGGCGCAACATCGGCCATAGACTTATGGCAATCTATGCAAACTAACAAGCGGGCGGCCTCTATGCGAGAGCCGCCCGTTTTGCATCCCTGCGGGCCAGTCGGAGACTGGCCAGCGGCAACTATATTCCGGGCCAAATCTTTTGGACGCGCCTTAGCGCCCCGGATGGCTACGCAGCGTTAAGCTAGTGCCTAACAATGTGTCAGTAAGCTCGTCAACGTAAACCTCTCTCAGAATTGAGGACGGTTTGCGTTGATCCCCCTAGCAGGCCCCCCTTGTTCCTGTATACACTTACTGACACAAAAAACAGGAGTTAAAGAATATGTGCAGTGATTTAATCCATATCGACGGGCAGCAATTTGCCTGTCGTAAGTGTAATGAGTGCATCACCGCTAGGAAAAATGGTTGGGTTGCTCGTGCGATGGCTGAAAAGGCCGTCACTGCTGAAACTTTCAGCGTAACCCTAACTTATAATGATGCTACTCAGGAAAGCCGCGATGCGGCAAAGACGTTCGAGTATCGACACGTCAAAAACTGGATTAAGAACCTTAGGCGTCAAATAGAGTACACCACCGGCCAGACTGGCCTTCTCCGCTACCTTGTAGCGGGTGAGCGCGGTTCCGATAAGGGCCGCTGCCACTGGCATGTAATCTTGTTCTGTAACGCCGATATTTTAACGCTAGGAAAAATGACGCACTGGCCCTCTGGCAAACTTGCTGCCGATAGGGCCGAAATTATAACTACAGGAAAAAGGAAAAAACGTATTAACTGGTCTCTCTGGCCGTATGGCTTCGTCAGCTTTCAAGAGCCTGACCAGTGGGGCATGGAGTACGCCTTAGCGTATGCCCTCAAAGATCAGTTTAACATTGTCTCCGCTGCCGGTACGGCGCGGGAGGCTCACGTTTCCCGCACTTCTGCGGGTATGTTCCGCATGTCCAAAAAACCCCCAATCGGTTTCCCCTTTCTTGAGCGCAAGCTCAACGCGCTAGATGCGCGTGGTCAGCTTCCAGTTGACCTAAAAATAAGGGTTCCCGATTACAAAGGGTACTGGTATCCTACGGGAGCTATCCGCGAGTACATGCTCGACCGCTTGCGGATCAGCAATGAGCTTTACAAAGCTCAGCATGGGCGTAATGCGCCACAATGGACCTCGCTAACGCGAAGCGTTGAGCAAAACGAAAAAGATTGGGAAAGGTTAATCCATGGCACCGAAGCGCAAGAAGAGGAGCAAGTCGAAGACTTCGAGGAGTGGCAACGCTCAATCCTCCTCCGTACAAAAGAAATACGCCAACAACAGATCGACAGAGACACGCGAAAGCGATGTGGCGGGCTTTCTGCTTGCTCACGATGCCTTAACAGCCTCACGCCGCAAGATTTCGACGCGGCGGCGAGATGGGCGGAACGCCAAGCTCGAAAACACGGCGGCTATGACGCCGCCGAAAAACACTACCGCAGCGAAAACCGCTGCAATCCCTATTGTGGCTACAGGGAACTACCAACCCAGAAAAGAGCCTTCAAAAAAGGAGCATGACGACAGGAA

>ICBM5_gene_1
MADKATSPTTVSKKDRDDFELHVRLLSRAGYKTAEARTIAWLRGPAGLQEMLAGVKVAS

>ICBM5_gene_2
MSAIAGALISGGASLLGGLFGRSSASKQQARQNEYNKPINIRKRAEEGGFNPLLWAGQGNIQMQPGPSGIMGSAIANAGLALADGMSEQRQLDLERTKLKQDQERLDALIEKQTIRPKVGGIYAGSQQTPSVARAPGRPLMNGAPQPGSAPVFNPPTEYNPIPDDGPRLQTKVMRSDGMTSADPENPAEMEGDWWTWAREGTFWQNNNEILRRNTPETLHYKGRDALFPKMIDGARKAHKKAQEDFEKNPPKRRKLKGVNPNLNDKKW

>ICBM5_gene_3
MVMNMSKYQRPTNTRRESRTIAGRFRGGKLAPVMASAFRESESAILSQQVTYELDPIAGRMITPIMAELISVYVPVQAIDALKNPEEAYAGNTEVVRDKLLSGTPLFGLEDESEISKRLGVNPISVGGVKKVNEAARLAHNCAVNFLRQRKYVNTVKLLADNMNVTPALISQTVLDRLNAVLDPEDRVNGAVQLDLGNVRMPVEGVGVEYNAVPVTTPGWNDTENDRAADPHYFSDHAAGIRLAIASENGRPAVYAVGEGQNAGGISLTDFYNAELMDSLVRQMRQIVDDNPEYGEEMVTRWAHGLSVDNGKTPWILHQSQQMFGNQFRRAMDGANLDVAQSDMMQSLEFTVPVPPTELGGVVITFASVKPDETLGSQPHPFLSTTWEATNYVSDELARDPEAVTMRHLNSDVPMVDEDTRALYIGHNGLKKAYISYGFNRHLDPTTVEAKTAIWQLEVPMSVTPESVIYPEDLDHYPFADQLAEVCTYQVSSTATVRTPMVFGPTPVEELAQIETDNVFEDV

>ICBM5_gene_4
MQVNDAKRSVLLVGLNGGEVSVISADGEIIATEGVTAGRHKCSSWVPFMSNEGDELSFSGDVVPMVPNGGRVRPMAYGPGQFESGANPDFVVTSADRMARELDHKIRGLDQTAKKVEARIAQLNNLAERAKTPVEVAKEEKVDVIDDDDVSDVRSSGDDSDDTGPDFGAEKVGAE

>ICBM5_gene_5
MNDAPKKSNSGRKQSWLRAYARGLKGGTPKTLCESLRLWSIPQEQAQDVTKKTYLNAADGPMIWLGEQIVKRMEKAGYPSRIFCGYRSPEQQDKEFAEGDSKARAYQSPHQFYEAVDIIHKTKAWNVSQDYWDTLAAIVRVVEREYAIDLVHGYDWGWDSAHIEIALWRQVPKRQIGKTGSNYPPSPWEREQRFKELLPAVYAAQHRP

>ICBM5_gene_6
MCSDLIHIDGQQFACRKCNECITARKNGWVARAMAEKAVTAETFSVTLTYNDATQESRDAAKTFEYRHVKNWIKNLRRQIEYTTGQTGLLRYLVAGERGSDKGRCHWHVILFCNADILTLGKMTHWPSGKLAADRAEIITTGKRKKRINWSLWPYGFVSFQEPDQWGMEYALAYALKDQFNIVSAAGTAREAHVSRTSAGMFRMSKKPPIGFPFLERKLNALDARGQLPVDLKIRVPDYKGYWYPTGAIREYMLDRLRISNELYKAQHGRNAPQWTSLTRSVEQNEKDWERLIHGTEAQEEEQVEDFEEWQRSILLRTKEIRQQQIDRDTRKRCGGLSACSRCLNSLTPQDFDAAARWAERQARKHGGYDAAEKHYRSENRCNPYCGYRELPTQKRAFKKGA
