## Supplementary material for "New Microviridae isolated from Sulfitobacter reveals two cosmopolitan subfamilies of ssDNA phages infecting marine and terrestrial Alphaproteobacteria": SI_file_4

Table 1: Prophages predicted in this study in different bacterial strains

|  | **Bacterial strain containing prophage** | | | | | | | | | | **Bacterial strain without prophage, used for delimitation of prophage region** | | | | |
| --- | --- | --- | --- | --- | --- | --- | --- | --- | --- | --- | --- | --- | --- | --- | --- |
| **Name** | | **Accession number** | **Contig size (bp)** | **DNA type** |  | **Prophage region (first extraction)** | | | | | **Name** | **Accession number** | **Contig size (bp)** | **DNA type** | **Site prophage insertion** |
|  |  |  |  |  | Coordinates (bp) | Size (bps) | Size (bps) after truncation |  | Confidence borders | |  |  |  |  |  |
|  |  |  |  |  |  |  |  | General | Left prophage | Right prophage |  |  |  |  |  |
| *Acinetobacter lwoffii* strain SU1904 | | NZ_JACWEU010000076 | 5.673 | Contig | - | 5.673 | 5.673 | Ok | No borders | No borders | - | - | - | - | - |
| *Agrobacterium larrymoorei* strain CFBP5477 | | NZ_SWKE01000015 | 657.589 | ND* | 276.685 – 282.985 | 6.301 | 5.819 | ok | clear border | clear border | *Agrobacterium larrymoorei* strain CFBP5473 | CP039691.1 | 2.968.843 | circular chromosome |  |
| *Agrobacterium tumefaciens* strain 1D1108 plasmid pAt1D1108a | | NZ_CP032923 | 502.2­74 | plasmid | 443.354 – 449.213 | 5.860 | 5.860 | ok | clear border | clear border | *Agrobacterium sp.* H13-3 plasmid pAspH13-3a | CP002250 | 601.551 | circular chromosome |  |
| *Alistipes* sp. isolate P1-1 tig00001311 | | SCPE01000001 | 3.006.009 | ND | 641.176 – 646.874 | 5.699 | 5.699 | ok | clear border | clear border | *Alistipes onderdonkii* subsp. vulgaris 5NYCFAH2 | AP019738 | 3.312.682 | circular chromosome |  |
| *Alistipes onderdonkii* WAL 8169 = DSM 19147 C506DRAFT_scaffold00012.12 | | NZ_KB894552 | 100.104 | ND* | 77.340 – 83.132 | 5.793 | 4.177 | ok | clear border | clear border | *Alistipes onderdonkii* subsp. vulgaris 5CPYCFAH4 | AP019737 | 3.312.673 | circular chromosome |  |
| *Aphanizomenon* flos aquae WA102 3645 | | LJOW01000239 |  |  |  | 150 – 4055 cut | 3.588 |  | No borders | No borders |  |  |  |  |  |
| *Bacillus altitudinis strain DSM 26896* | | NZ_JXAI01000008 | 156.970 | whole genome shotgun sequence | - | 13.312 |  | Right border is missing | Clear border | Not clear | *Bacillus altitudinis strain BIM B-263 chromosome* | CP063360 | 3.763.717 | Circular chromosome |  |
| *Bacillus sp. X1(2014) strain DE0237 NODE_39* | | NZ_VECT01000039 | 33.618 | whole genome shotgun sequence | - | 7.309 |  | Left border is missing | Not clear | Clear border | *Bacillus sp. FJAT-22090* | CP012601 | 4.215.291 | Circular chromosome |  |
| *Bacillus thuringiensis serovar T01001* | | NZ_CM000748 | 6.323.123 | Circular chromosome | - | 11.378 |  | Right border is missing | Clear border | Not clear | *Bacillus thuringiensis strain JW-1 plasmid p1* | CP045023 | 359.606 | Circular plasmid |  |
| *Bacillus wiedmannii strain FSL J3-0113 NODE_13* | | NZ_LXFN01000004 | 109.662 | whole genome shotgun sequence | - | 9.571 |  | Ok | Clear border | Clear border | *Bacillus wiedmannii bv. thuringiensis strain FCC41 plasmid pFCC41-1-490K* | CP024685 | 490.693 | Circular plasmid |  |
| *Bacteroides caccae* strain 2789STDY5834946 | | NZ_CZBL01000008 | 221.420 | ND* | 86.697 – 92.267 | 5.571 | 4.542 | ok | clear border | clear border | *Bacteroides caccae* strain ATCC 43185 | CP022412 | 4.570.803 | circular chromosome |  |
| *Bacteroides caccae strain* AF46-5GN AF46-5GN.Scaf95 | | QRNA01000095 | 19.734 | ND* |  | 5.370 | 4.634 | Right border is missing | clear border | Not clear | *Bacteroides caccae* strain ATCC 43185 | CP022412 | 4.570.803 | circular chromosome |  |
| *Bacteroides eggerthii* DSM 20697 | | NZ_DS995510 | 19.345 | ND* |  | 6.195 | 6.195 | ok | clear border | clear border | *Bacteroides* sp. A1C1 | CP036491 | 4.502.190 | circular chromosome |  |
| *Bacteroides finegoldii strain D54t1_190329_G10 NODE_5* | | NZ_JADNKZ010000005 | 317.103 | whole genome shotgun sequence | - | 6.471 | 6.471 | Ok | Clear border | Clear border | *Bacteroides sp. HF-5287 chromosome* | CP059856 | 5.244.518 | Circular chromosome |  |
| *Bacteroides ovatus* isolate Bacteroides_ovatus_MC1 | | NZ_CAAKNR010000141 | 59.930 | ND* |  | 6.402 | 5.167 | ok | clear border | clear border | *Bacteroides ovatus* strain FDAARGOS_733 | CP046397 | 6.571.392 | circular chromosome |  |
| *Bacteroides plebeius*  DSM 17135 | | NZ_ABQC02000012 | 569.841 | whole genome shotgun sequence | 90.389 – 95.448 | 5.059 | 5.059 | ok |  |  | ND* |  |  |  |  |
| *Bacteroides sp. 2_2_4* supercont1.3 | | NZ_EQ973357 | 605.506 | ND* |  | 6.303 | 6.303 | ok | clear border | clear border | *Bacteroides xylanisolvens strain H207* | CP041230 | 6.499.434 | circular chromosome |  |
| *Bacteroides sp.* AF29-11 AF29-11.Scaf2 | | NZ_QTLY01000002 | 622.059 | ND* |  | 6.481 | 5.440 | ok | clear border | clear border | *Bacteroides uniformis* NBRC 113350 | AP019724 | 4.734.883 | circular chromosome |  |
| *Bacteroides thetaiotaomicron strain* 19_BTHE 1034_17932_216065 | | NZ_JVQR01000241 | 17.932 | ND* |  | 6.056 | 6.056 | ok | clear border | clear border | *Bacteroides thetaiotaomicron F9-2 DNA,* nearly complete genome | AP022660 | 6.224.560 | circular chromosome |  |
| *Bacteroides xylanisolvens* strain AF38-2 AF38-2.Scaf17 | | NZ_QROO01000017 | 141.458 | ND* | 11.921 – 20.144 | 8.224 | 5.470 | ok | clear border | clear border | *Bacteroides xylanisolvens* XB1A | FP929033 | 5.976.145 | circular chromosome |  |
| Candidatus *Rhodobacter lobularis* isolate IGS | | NZ_LFTY01000002rueg | 3.995.303 | ND* | 1240322-1248061 | 7.740 | 4.335 | Uncertain borders, probably includes bacterial regions |  |  | ND* |  |  |  |  |
| *Clostridioides difficile CD129* | | NZ_AVHP01000907 | 5.697 | Contig | - | 5.697 |  | Ok | No borders | No borders | *-* | - | - | - | - |
| *Coprobacter fastidiosus isolate CIM:MAG 335* | | QALP01000027 | 20.190 | whole genome shotgun sequence | - | 6.609 | 6.609 | Ok | Clear border | Clear border | *Coprobacter sp. 2CBH44 DNA* | AP023322 | 4.171.466 | Circular chromosome |  |
| *Devosia chinhatensis* strain IPL18 | | NZ_JZEY01000054 | 2139066 | ND* | 1315127-1322625 | 7.499 | 4.697 |  | Low match alignment, kept more bases in the prophage. | Low match alignment, kept more bases in the prophage. | *Devosia* sp. I507 | CP026747 | 4005916 | circular chromosome | 3642584-3641978 |
| *Devosia* sp. YR412 | | NZ_FOFL01000005 | 221727 | ND* | 161269-172673 | 11.405 | 5.794 | Uncertain borders, probably includes bacterial regions |  |  | ND* |  |  |  |  |
| *Dysgonomonas macrotermitis* strain DSM 27370 | | NZ_FQUC01000005 | 265.550 | ND* | 30.733 – 36.871 | 6.139 | 4.862 | Left border is missing | Not clear | clear border | *Dysgonomonas* sp. HDW5A | CP049857 | 4.298.355 | circular chromosome |  |
| *Dysgonomonas* sp. 521 scaffold10_size73398 | | NZ_QVMO01000010 | 73.398 | ND* | 63.894 – 70.709 | 6.816 | 4.286 | Right border is missing | clear border | Not clear | *Dysgonomonas sp.* HDW5B | CP049858 | 4.402.003 | circular chromosome |  |
| *Elizabethkingia anophelis* strain E6809 2b | | NZ_MAHS01000003 | 1.150.786 | ND* | 987128-991519 | 4.392 | 3.338 | ok | Clear border | Clear border | Elizabethkingia anophelis strain 0422 | CP016370 | 3986480 | Chromosome, cicular | 3547444 |
| *Epibacterium ulvae* strain U95 | | NZ_PHJF01000001 | 804.761 | ND* | 626474-634343 | 7.870 | 5.418 | Uncertain borders, probably includes bacterial regions |  |  | ND* |  |  |  |  |
| *Erysipelatoclostridium* sp. An15 An15_contig_1, | | NZ_NFLA01000001 | 201.071 | ND* | 158.396 – 164.886 | 6.491 | 4.566 | Not ok | Not clear | Not clear | *Erysipelotrichaceae bacterium* GAM147 | AP018537 | 2.753.585 | circular chromosome |  |
| *Escherichia coli strain OLC1558 OLC1558* | | NZ_NWRV01000002 | 351.548 | whole genome shotgun sequence | - | 4.477 | 4.477 | Ok | Clear border | Clear border | *Escherichia coli O157:H7 strain USDA5905 chromosome* | CP039837 | 5.431.628 | Circular chromosome |  |
| *Escherichia sp.* MOD1-EC6163 | | NZ_PTSA01000028 | 50.742 | ND* |  | 5.993 | 4.140 | ok | clear border | clear border | *Escherichia coli* strain NCTC11133 | LR134340 | 4.450.344 | circular chromosome |  |
| *Gramella jeungdoensis strain KCTC 23123 KCTC_32123* | | NZ_SNQI01000008 | 6.302 | Contig | - | 6.302 | 6.302 | Ok | No borders | No borders | *-* | - | - | - | - |
| *Kosakonia cowanii strain Esp_Z* | | NZ_CP022690 | 5.077.975 | Circular chromosome | - | 5.002 | 5.002 | Ok | Clear border | Clear border | *Kosakonia cowanii strain FBS 223 chromosome* | CP035129 | 4.685.999 | Circular chromosome |  |
| *Labrys sp. WJW* | | NZ_LYXZ01000012 | 94.779 | ND* | 49150-58517 | 9.368 | 5.762 | Uncertain borders, probably includes bacterial regions |  |  | ND* |  |  |  |  |
| *Listeria monocytogenes strain C5* | | NZ_MDQI01000007 | 1.240.180 | whole genome shotgun sequence | - | 10.033 |  | Ok | Clear border | Clear border | *Listeria monocytogenes strain N12-0935 chromosome* | CP038642 | 2.986.680 | Circular chromosome |  |
| *Listeria monocytogenes strain 2014L-6088 NODE_2* | | NZ_JABXNI010000003 | 593.093 | whole genome shotgun sequence | - | 6.184 |  | Ok | Clear border | Clear border | *Listeria monocytogenes strain PIR00540 chromosome* | CP025568 | 3.026.819 | Circular chromosome |  |
| *Mammaliicoccus sciuri strain GDM7P051A C-1* | | NZ_WIVK01000025 | 5.855 | Contig | - | 5.855 | 5.855 | Ok | No borders | No borders | *-* | - | - | - | - |
| *Maritimibacter sp. LZ-17 NODE_6* | | NZ_SWKQ01000016 | 114.540 | whole genome shotgun sequence | - | 8.679 | 8.005 | Ok | Not clear | Clear border | *MAG Maritimibacter sp. isolate ih11 chromosome* | CP051232 | 4.675.343 | Circular chromosome |  |
| *Mesorhizobium sp. 3P27G6 NODE_30* | | NZ_SCNN01000030 | 67.438 | ND* | 41.558 – 47.582 | 6.025 | 5.251 | ok | clear border | clear border | *Mesorhizobium sp. NZP2234* | CP033364 | 6.749.717 | circular chromosome |  |
| *Mesorhizobium composti strain* CC-YTH430 contig01 | | NZ_SSNY01000001 | 699.301 | ND* | 612.933 – 618.763 | 5.831 | 5.624 | ok | clear border | clear border | *Mesorhizobium composti* strain CC-YTH430 contig01 | NZ_SSNY01000001 | 699.301 | circular chromosome |  |
| *Mesorhizobium sp. M7A.F.Ca.ET.027.02.1.1 NODE_4* | | NZ_RZOL01000004 | 56.052 | ND* |  | 6.940 | 5.158 | ok | clear border | clear border | *Mesorhizobium sp. WSM1497* | CP021070 | 6.666.492 | circular chromosome |  |
| *Mesorhizobium sp. M1A.F.Ca.IN.020.30.1.1 NODE_39* | | NZ_RZSO01000039 | 17.913 | ND* |  | 7.714 | 5.264 | ok | clear border | clear border | *Mesorhizobium sp. M1D.F.Ca.ET.043.01.1.1* | CP034444 | 7.127.907 | circular chromosome |  |
| *Mesorhizobium sp. M1A.F.Ca.ET.072.01.1.1 NODE_40* | | NZ_RZSF01000040 | 32.967 | ND* |  | 8.388 | 5.854 | ok | clear border | clear border | *Mesorhizobium sp. M1B.F.Ca.ET.045.04.1.1* | CP034448 | 7.779.814 | circular chromosome |  |
| *Mesorhizobium sp. M1A.F.Ca.IN.022.06.1.1* | | CP034455 | 6.278.406 | circular chromosome | 27.176 – 34.340 | 7.165 | 5.591 | ok | clear border | clear border | *Mesorhizobium sp. M1E.F.Ca.ET.045.02.1.1* | CP034447 | 7.348.390 | circular chromosome |  |
| *Mesorhizobium sp. isolate N.Cr.TU.016.05.1 NODE_86* | | SASU01000086 | 23.594 | ND* |  | 6.165 | 5.598 | ok | clear border | clear border | *Mesorhizobium* sp. M6A.T.Cr.TU.016.01.1.1 | CP034452 | 6.784.580 | circular chromosome |  |
| *Mesorhizobium sp. M5C.F.Ca.IN.020.29.1.1 NODE_361* | | NZ_RZSW01000361 | 6.491 | Contig | - | 6.491 | 6.491 | Ok | No borders | No borders | *-* | - | - | - | - |
| *Neorhizobium* sp. T17_20 | | NZ_PVBG01000001 | 571.600 | ND* | 485319-491745 | 6.427 | 5.896 | Ok (it could be shorter than the true prophage, but not by long) | Good aligment, clear border. | Border given by the 2nd repeat. | *Neorhizobium galegae* | HG938353 | 4647962 | circular chromosome | 1787722 |
|  |  |  |  |  | 485319-498276 | 12.958 |  |  | Good aligment, clear border. | The reference has a 115 bases unaligned region, followed by good alignment with the lysogenic strain. Therefore, on this side the border is unclear. |  |  |  |  |  |
| *Novosphingobium tardaugens* NBRC 16725 | | NZ_BASZ01000013 | 102.866 | ND* | 78084-82527 | 4.443 |  |  | Clear borders, but broader on both sides than what is mentioned in the Citromicrobium paper | |  |  |  |  |  |
| *Nioella nitratireducens* strain SSW136 | | NZ_MNBW01000014 | 95.120 | ND* | 65144-70877 | 5.734 | 5.233 | ok | This is only the repeat region, with a high confidence that it includes only phage sequence. | | *Roseibacterium elongatum* DSM 19469 | CP004372 | 3555109 | Chromosome, circular | 753094 |
|  |  |  |  |  | 64851-76635 | 11.785 |  |  | The longer region (see below) has uncertain borders on both sides. | |  |  |  |  |  |
| *Nitrospira sp. isolate RSF5* | | SWDO01000020 | 96.140 | whole genome shotgun sequence | - | 6.250 |  | Ok | Clear border | Clear border | *Nitrospira sp. KM1 DNA* | AP022671 | 4.509.223 | Circular chromosome |  |
| *Oceanicola* sp. S124 | | NZ_AFPM01000156 | 38.215 | ND* | 3924-13038 | 9.115 | 5.231 | Uncertain borders, probably include bacterial regions |  |  | ND* |  |  |  |  |
| *Oscillibacter sp. PC13* | | NZ_FOXE01000003 | 177.285 | ND* | 141.478 – 146.894 | 5.417 | 4.843 | ok | clear border | clear border | *Oscillibacter sp. PEA192 DNA* | AP018532 | 3.320.462 | circular chromosome |  |
| *Paenibacillus odorifer strain DSM 15391* | | NZ_CP009428 | 6.812.473 | Circular chromosome | - | 8.259 | 8.259 | Left border is missing | Not clear | Clear border | *Paenibacillus odorifer isolate MGYG-HGUT-02414* | LR698998 | 6.802.552 | Linear chromosome |  |
| *Parabacteroides distasonis str. 3999B* | | NZ_JNHQ01000100 | 85.604 | ND* | 74.055 – 78.680 | 4.626 | 4.626 | ok | clear border | clear border | *Parabacteroides distasonis strain FDAARGOS_759* | CP054012 | 4.926.033 | circular chromosome |  |
| *Parabacteroides sp. AF39-10AC* | | NZ_QTLP01000027 | 76.059 | ND* | 33.785 – 38.367 | 4.583 | 4.583 | ok | clear border | clear border | *Parabacteroides sp. CT06* | CP022754 | 5.372.666 | circular chromosome |  |
| *Parabacteroides sp. AM27-42* | | NZ_QTNL01000010 | 114.974 | ND* | 53.179 – 58.161 | 4.983 | 4.018 | ok | clear border | clear border | *Parabacteroides sp. CT06* | CP022754 | 5.372.666 | circular chromosome |  |
| *Paracoccus alkenifer* strain DSM 11593 | | NZ_FNXG01000002 | 871.604 | ND* | 841755-849952 | 8.198 | 4.929 | Uncertain borders, probably include bacterial regions |  |  | ND* |  |  |  |  |
| *Phaeobacter inhibens* strain DOK1-1 | | NZ_CP019307 | 3.718.082 | Chromosome, circular | 411908-417829 | 5.922 | 5.477 | ok | Good aligment, clear border. | Good aligment, clear border. | Phaeobacter inhibens *DSM 17395* | NC_018290.1 | 3821831 | Chromosome, circular | 1630134 |
| *Phascolarctobacterium faecium DSM 14760* | | NZ_QLTS01000001 | 796.791 | ND* | 90.809 – 95.325 | 4.517 | 4.297 | ok | clear border | clear border | *Phascolarctobacterium faecium JCM 30894* | AP019004 | 2.454.371 | circular chromosome |  |
| *Prevotella bergensis* DSM 17361 | | NZ_GG704781 | 1.108.685 | whole genome shotgun sequence | 390.637 – 395.974 | 5.337 | 5.337 | ok | - | - | ND* | - | - | - |  |
| *Prevotella buccalis* ATCC 35310 | | NZ_ADEG01000016 | 88.363 | whole genome shotgun sequence | 70.195 – 75.170 | 4.975 | 4.975 | ok | clear border | clear border | ND* | - | - | - |  |
| *Prevotella sp. CAG:1185* | | HF992559 | 34.922 | whole genome shotgun sequence | - | 6.349 | 6.349 | Ok | Clear border | Clear border | *Prevotella sp. WR041 DNA* | AP024484 | 3.524.789 | Circular chromosome |  |
| *Prevotella salivae F0493* | | NZ_AWGW01000011 | 6.888 | Contig | - | 6.888 | 6.888 | Ok | No borders | No borders | *-* | - | - | - | - |
| *Ralstonia solanacearum strain UW179* | | NZ_CDLZ01000001 | 5.426.414 | whole genome shotgun sequence | - | 6.503 |  | Ok | Clear border | Clear border | *Ralstonia solanacearum strain CFBP 8695 plasmid unnamed* | CP047139 | 1.814.285 | Circular chromosome |  |
| *Rhizobium pusense* strain CCGM10 | | NZ_KV878023 | 486.269 | ND* | 21543-30486 | 8.944 | 5.207 | Uncertain borders, probably include bacterial regions |  |  | ND* |  |  |  |  |
| *Rhizobium sp. BG4 plasmid pRPC1* | | CP044126 | 1.609.137 | Circular plasmid | - | 6.609 | 6.609 | Ok | Clear border | Clear border | *Rhizobium sp. BG6 chromosome* | CP044009 | 3.928.948 | Circular chromosome |  |
| *Rhizobium sp. MHM7A NODE_12* | | NZ_VCHT01000012 | 221.567 | whole genome shotgun sequence | - | 6.733 | 6.542 | Left border is missing | Not clear | Clear border | *Rhizobium sp. 007 chromosome* | CP064187 | 4.296.727 | Circular chromosome |  |
| *Rhizobium* sp. Root564 | | NZ_LMGN01000011 | 1.029.951 | ND* | 90483-101008 | 10.526 | 5.781 | Uncertain borders, probably include bacterial regions |  |  | ND* |  |  |  |  |
| *Rhizobium* sp. Root651 | | NZ_LMHB01000005 | 167.271 | ND* | 59726-68152 | 8.427 | 5.254 | Uncertain borders, probably include bacterial regions |  |  | ND* |  |  |  |  |
| *Rhizobium* sp. UR51a | | NZ_JYFU01000029 | 277.936 | ND* | 236062-242003 | 5.942 | 5.139 |  | Good alignment, clear border. | The reference has a 2274 bases unaligned region, followed by good alignment with the lysogenic strain. Therefore, on this side the border is unclear. | *Rhizobium* sp. Y9 | CP017999 | 2746556 | Chromosome, circular | 89840 |
| *Rhizobium* sp. YS-1r | | NZ_JPYQ01000001 | 657.089 | ND* | 592218-600787 | 8.570 | 5.748 | Uncertain borders, probably include bacterial regions |  |  | ND* |  |  |  |  |
| *Rhizobium straminoryzae strain SM12* | | NZ_VJMG01000008 | 175.733 | whole genome shotgun sequence | - | 7.361 | 5.995 | Ok | Clear border | Clear border | *Rhizobium pseudoryzae strain DSM 19479 chromosome* | CP049241 | 3.646.231 | Circular chromosome |  |
| *Rhodobacter capsulatus* YW2 | | NZ_AYPZ01000002 | 316.953 | ND* | 251239-257664 | 6.427 | 5.323 | ok | Good alignment, clear border. | Good alignment, clear border. | *Rhodobacter capsulatus* SB 1003 | NC_014034 | 3738958 | Chromosome, circular | 1983981 |
| *Rhodobacter capsulatus* R121 | | NZ_AYQC01000019 | 418.689 | ND* | 140693-147303 | 6.611 | 5.616 | ok | Good alignment, clear border. | Good alignment, clear border. |  |  |  |  | 1983981 |
| *Rhodobacter* sp. SW2 | | NZ_ACYY01000005 | 188.067 | ND* | 100643-110387 | 9.745 | 5.104 | Uncertain borders, probably include bacterial regions |  |  | ND* |  |  |  |  |
| *Rhodovulum sulfidophilum* strain DSM 2351 | | NZ_AP014801 | 111.306 | Plasmid , circular | 79745-85953 | 6.209 | 5.265 | ok | Good alignment, clear border | Good alignment, clear border | *Rhodovulum sulfidophilum* DSM 1374 | NZ_CP015420.1 | 102180 | Plasmid, circular? | 6489 |
| *Rhodovulum* sp. MB263 | | NZ_CP020384 | 3.860.570 | Chromosome, circular | 1996124-2002706 | 6.583 | 5.262 |  | Good alignment, clear border | The reference has 301 bases of unaligned region, followed by good alignment with the lysogenic. Therefore, on this side the border is unclear. | *Rhodovulum sulfidophilum* strain SNK001 | CP015421 | 4082971 | Chromosome, circular | 2715097 |
| *Roseovarius* sp. GCL-8 | | NZ_QITQ01000003 | 282.961 | ND* | 195522-205912 | 10.391 | 5.603 | Uncertain borders, probably include bacterial regions |  |  | - | - | - | - | - |
|  | |  |  |  | 197814-204437 | 6.624 |  | Borders unclear, they align well, but with different LCBs |  |  | Sulfitobacter sp. SK025 chromosome | CP025808 | 3.035*.350 | Chromosome, circular | - |
| *Ruegeria sp. THAF57* | | NZ_CAIWNQ010000009 | 210.182 | whole genome shotgun sequence | - | 13.152 | 6.442 | Right border is missing | Clear border | Not clear | *Ruegeria sp. AD91A plasmid unnamed1* | CP031947 | 743.267 | Circular plasmid |  |
| *Ruegeria mobilis* strain S1942 | | NZ_JXYG01000006 | 603.960 | ND | 213142-218944 | 5.802 | 5.369 | Ok | Good alignment, clear border | Good alignment, clear border | *Ruegeria mobilis* strain DSM 23403 | NZ_FNNK01000005.1 | 322.985 | ND | 85656 |
| *Stenotrophomonas maltophilia strain AS012690* | | NZ_VLGL01000002 | 130.645 | whole genome shotgun sequence | - | 11.045 |  | Right border is missing | Clear border | Not clear | *Stenotrophomonas maltophilia strain NEB515 chromosome* | CP051467 | 4.785.515 | Circular chromosome |  |
| *Stenotrophomonas rhizophila strain JC1* | | CP050062 | 4.268.161 | circular chromosome | 2.031.617 – 2.038.577 | 6.961 |  | Right border is missing | clear border | Not clear | *Stenotrophomonas rhizophila strain DSM14405* | CP007597 | 4.648.976 | circular chromosome |  |
| *Xanthomonas citri pv. citri strain LE3-1* | | NZ_LN647176 | 5.045.425 | whole genome shotgun sequence | - | 6.748 |  | Ok | Clear border | Clear border | *Xanthomonas citri pv. citri strain DAR84832 chromosome* | CP060460 | 5.223.652 | Circular chromosome |  |

*ND – not determined

(Yellow = new prophages, red = not *Microviridae*, grey = old prophages, blue = sequencing contaminants)

Table 2: Predicted prophages: host habitat and host classification

| **Lysogenic strain** | **Lineage** | **Order** | **Habitat / isolation site** | **Reference habitat** |
| --- | --- | --- | --- | --- |
| [*Acinetobacter lwoffii*](https://lpsn.dsmz.de/species/acinetobacter-lwoffii) strain SU1904 | Bacteria; Proteobacteria; Gammaproteobacteria; Pseudomonadales; Moraxellaceae; Acinetobacter; Acinetobacter calcoaceticus/baumannii complex | Pseudomonadales | Clinical isolate | [Hayashi et al. 2020](https://pubmed.ncbi.nlm.nih.gov/33250496/) |
| [*Agrobacterium larrymoorei*](https://lpsn.dsmz.de/species/agrobacterium-larrymoorei) strain CFBP5477 | Bacteria; Proteobacteria; Alphaproteobacteria; Hyphomicrobiales; Rhizobiaceae; Rhizobium/Agrobacterium group; Agrobacterium | Hyphomicrobiales | isolated from tumours growing on pruned branches of Ficus benjamina | [Bouzar et al. 2001](https://pubmed.ncbi.nlm.nih.gov/11411669/) |
| [*Agrobacterium tumefaciens*](https://lpsn.dsmz.de/species/agrobacterium-tumefaciens) strain 1D1108 plasmid pAt1D1108a | Bacteria; Proteobacteria; Alphaproteobacteria; Hyphomicrobiales; Rhizobiaceae; Rhizobium/Agrobacterium group; Agrobacterium | Hyphomicrobiales | Terrestrial, Soil, plant pathogens (tumor producing, not nitrogen fixer) | [Conn et al. 1942](https://pubmed.ncbi.nlm.nih.gov/16560572/) |
| [*Alistipes*](https://lpsn.dsmz.de/genus/alistipes) sp*.* isolate P1-1 | Bacteria; Bacteroidetes; Bacteroidia; Bacteroidales; Rikenellaceae; Alistipes | Bacteroidales | Human specimens | [Song et al. 2006](https://pubmed.ncbi.nlm.nih.gov/16902041/) |
| [*Alistipes onderdonkii*](https://lpsn.dsmz.de/species/alistipes-onderdonkii) WAL 8169 = DSM 19147 | Bacteria; Bacteroidetes; Bacteroidia; Bacteroidales; Rikenellaceae; Alistipes | Bacteroidales | Human specimens | [Song et al. 2006](https://pubmed.ncbi.nlm.nih.gov/16902041/) |
| [*Aphanizomenon flos aquae*](https://lpsn.dsmz.de/species/aphanizomenon-flosaquae) WA102 3645 | Bacteria; Cyanobacteria; Nostocales; Aphanizomenonaceae; Aphanizomenon. | Nostocales | Freshwater | [NCBI entry](https://www.ncbi.nlm.nih.gov/nuccore/LJOW01000239) |
| [*Bacillus altitudinis*](https://lpsn.dsmz.de/species/bacillus-altitudinis) strain DSM 26896 | Bacteria; Firmicutes; Bacilli; Bacillales; Bacillaceae; Bacillus | Bacillales | Marine, gut of marine fish | [Esakkiraj et al. 2012](https://www.sciencedirect.com/science/article/abs/pii/S0960308511001167?via%3Dihub) |
| [*Bacillus*](https://lpsn.dsmz.de/genus/bacillus) *sp. X1*  strain DE0237 NODE_39 | Bacteria; Firmicutes; Bacilli; Bacillales; Bacillaceae; Bacillus | Bacillales | Environmental sample | [Zhang et al. 2020](https://pubmed.ncbi.nlm.nih.gov/32747578/) |
| [*Bacillus thuringiensis*](https://lpsn.dsmz.de/species/bacillus-thuringiensis) serovar thuringiensis | Bacteria; Firmicutes; Bacilli; Bacillales; Bacillaceae; Bacillus; Bacillus cereus group | Bacillales | Moth larvae, animal tissue | [Ibrahim et al. 2010](https://www.ncbi.nlm.nih.gov/pmc/articles/PMC3035146/) |
| [*Bacillus wiedmannii*](https://lpsn.dsmz.de/species/bacillus-wiedmannii) strain FSL J3-0113 | Bacteria; Firmicutes; Bacilli; Bacillales; Bacillaceae; Bacillus; Bacillus cereus group | Bacillales | Raw milk | [NCBI entry](https://www.ncbi.nlm.nih.gov/nuccore/NZ_LXFN00000000.1) |
| [*Bacteroides caccae*](https://lpsn.dsmz.de/species/bacteroides-caccae) strain 2789STDY5834946 | Bacteria; Bacteroidetes; Bacteroidia; Bacteroidales; Bacteroidaceae; Bacteroides | Bacteroidales | Human feces | [NCBI entry](https://www.ncbi.nlm.nih.gov/nuccore/NZ_CZBL00000000.1) |
| [*Bacteroides caccae*](https://lpsn.dsmz.de/species/bacteroides-caccae) strain AF46-5GN | Bacteria; Bacteroidetes; Bacteroidia; Bacteroidales; Bacteroidaceae; Bacteroides | Bacteroidales | Human feces | [NCBI entry](https://www.ncbi.nlm.nih.gov/nuccore/QRNA01000001.1) |
| [*Bacteroides eggerthii*](https://lpsn.dsmz.de/species/bacteroides-eggerthii) DSM 20697 | Bacteria; Bacteroidetes; Bacteroidia; Bacteroidales; Bacteroidaceae; Bacteroides | Bacteroidales | Human feces | [NCBI entry](https://www.ncbi.nlm.nih.gov/nuccore/NZ_ABVO00000000.1) |
| [*Bacteroides finegoldii*](https://lpsn.dsmz.de/species/bacteroides-finegoldii)  strain D54t1_190329_G10 | Bacteria; Bacteroidetes; Bacteroidia; Bacteroidales; Bacteroidaceae; Bacteroides | Bacteroidales | Human feces | [NCBI entry](https://www.ncbi.nlm.nih.gov/nuccore/NZ_JADNKZ000000000.1) |
| [*Bacteroides ovatus*](https://lpsn.dsmz.de/species/bacteroides-ovatus) isolate Bacteroides_ovatus_MC1 | Bacteria; Bacteroidetes; Bacteroidia; Bacteroidales; Bacteroidaceae; Bacteroides | Bacteroidales | Human feces | [Eggerth et al. 1933](https://pubmed.ncbi.nlm.nih.gov/16559622/) |
| [*Bacteroides plebeius*](https://lpsn.dsmz.de/species/phocaeicola-plebeius)  DSM 17135  New name: *Phocaeicola plebeius* | Bacteria; Bacteroidetes; Bacteroidia; Bacteroidales; Bacteroidaceae; Bacteroides | Bacteroidales | Human feces | [NCBI entry](https://www.ncbi.nlm.nih.gov/nuccore/NZ_ABQC00000000.2) |
| [*Bacteroides*](https://lpsn.dsmz.de/genus/bacteroides) sp. 2_2_4 supercont1.3 | Bacteria; Bacteroidetes; Bacteroidia; Bacteroidales; Bacteroidaceae; Bacteroides | Bacteroidales | Human feces | [NCBI entry](https://www.ncbi.nlm.nih.gov/nuccore/NZ_ABZZ00000000.1) |
| [*Bacteroides*](https://lpsn.dsmz.de/genus/bacteroides) sp. AF29-11 | Bacteria; Bacteroidetes; Bacteroidia; Bacteroidales; Bacteroidaceae; Bacteroides | Bacteroidales | Human feces | [NCBI entry](https://www.ncbi.nlm.nih.gov/nuccore/NZ_QTLY00000000.1) |
| [*Bacteroides thetaiotaomicron*](https://lpsn.dsmz.de/species/bacteroides-thetaiotaomicron) strain 19_BTHE 1034_17932_216065 | Bacteria; Bacteroidetes; Bacteroidia; Bacteroidales; Bacteroidaceae; Bacteroides | Bacteroidales | Human sample | [Castellani A et al. 1919](https://www.ncbi.nlm.nih.gov/Taxonomy/Browser/wwwtax.cgi?mode=Info&id=818&lvl=3&lin=f) |
| [*Bacteroides xylanisolvens*](https://lpsn.dsmz.de/species/bacteroides-xylanisolvens) strain AF38-2 | Bacteria; Bacteroidetes; Bacteroidia; Bacteroidales; Bacteroidaceae; Bacteroides | Bacteroidales | Human feces | [Chassard C et al. 2008](https://pubmed.ncbi.nlm.nih.gov/18398210/) |
| [Candidatus *Rhodobacter lobularis*](https://lpsn.dsmz.de/species/rhodobacter-lobularis) isolate IGS | Bacteria; Proteobacteria; Alphaproteobacteria; Rhodobacterales; Rhodobacteraceae; Rhodobacter | Rhodobacterales | Marine, sponge-associated (in Mediterranean Sea) | [Jourda et al. 2015](https://pubmed.ncbi.nlm.nih.gov/26337883/) |
| [*Clostridioides difficile*](https://lpsn.dsmz.de/species/clostridioides-difficile) CD129 | Bacteria; Firmicutes; Clostridia; Eubacteriales; Peptostreptococcaceae; Clostridioides | Eubacteriales | Human feces | [Lawson et al. 2016](https://pubmed.ncbi.nlm.nih.gov/27370902/) |
| [*Coprobacter fastidiosus*](https://lpsn.dsmz.de/species/coprobacter-fastidiosus) isolate CIM:MAG 335 contig_33578, | Bacteria; Bacteroidetes; Bacteroidia; Bacteroidales; Barnesiellaceae; Coprobacter | Bacteroidales | Human feces | [Shkoporov et al. 2013](https://pubmed.ncbi.nlm.nih.gov/23771624/) |
| [*Devosia chinhatensis*](https://lpsn.dsmz.de/species/devosia-chinhatensis) strain IPL18 | Bacteria; Proteobacteria; Alphaproteobacteria; Hyphomicrobiales; Hyphomicrobiaceae; Devosia | Hyphomicrobiales | Terrestrial, Soil (oil contaminated) | [Hassan et al. 2015](https://www.ncbi.nlm.nih.gov/pmc/articles/PMC4541263/) |
| [*Devosia*](https://lpsn.dsmz.de/genus/devosia) sp. YR412 | Bacteria; Proteobacteria; Alphaproteobacteria; Hyphomicrobiales; Hyphomicrobiaceae; Devosia. | Hyphomicrobiales | Terrestrial, root associated | [JGI entry](https://gold.jgi.doe.gov/project?id=Gp0136776) |
| [*Dysgonomonas macrotermitis*](https://lpsn.dsmz.de/species/dysgonomonas-macrotermitis) strain DSM 27370 | Bacteria; Bacteroidetes; Bacteroidia; Bacteroidales; Dysgonamonadaceae; Dysgonomonas | Bacteroidales | from the hindgut of a fungus-growing termite *Macrotermes barneyi* | [Yang YJ et al. 2014](https://pubmed.ncbi.nlm.nih.gov/24899656/) |
| [*Dysgonomonas*](https://lpsn.dsmz.de/genus/dysgonomonas) sp. 521 | Bacteria; Bacteroidetes; Bacteroidia; Bacteroidales; Dysgonamonadaceae; Dysgonomonas | Bacteroidales | Alimentary canal | [NCBI entry](https://www.ncbi.nlm.nih.gov/nuccore/NZ_QVMO00000000.1) |
| [*Elizabethkingia anophelis* strain E6809 2b](https://lpsn.dsmz.de/species/elizabethkingia-anophelis) | Bacteria; Bacteroidetes; Flavobacteriia; Flavobacteriales; Flavobacteriaceae; Elizabethkingia | Flavobacteriales | Human blood | [Nicholson et al. 2016](https://pubmed.ncbi.nlm.nih.gov/26966213/) |
| [*Epibacterium ulvae* strain U95](https://lpsn.dsmz.de/species/epibacterium-ulvae) | Bacteria; Proteobacteria; Alphaproteobacteria; Rhodobacterales; Rhodobacteraceae; Epibacterium | Rhodobacterales | Marine, isolated from macro algae | [Breider et al. 2019](https://environmentalmicrobiome.biomedcentral.com/articles/10.1186/s40793-019-0343-5) |
| [*Erysipelatoclostridium*](https://lpsn.dsmz.de/genus/erysipelatoclostridium) sp. An15 | Bacteria; Firmicutes; Erysipelotrichia; Erysipelotrichales; Erysipelotrichaceae; Erysipelatoclostridium; unclassified Erysipelatoclostridium | Erysipelotrichales | Animal tissue (Rooster) | [NCBI entry](https://www.ncbi.nlm.nih.gov/nuccore/NZ_NFLA00000000.1) |
| [*Escherichia coli*](https://lpsn.dsmz.de/species/escherichia-coli) strain OLC1558 | Bacteria; Proteobacteria; Gammaproteobacteria; Enterobacterales; Enterobacteriaceae; Escherichia | Enterobacterales | unknown | NCBI entry |
| [*Escherichia*](https://lpsn.dsmz.de/genus/escherichia) sp. MOD1-EC6163 | Bacteria; Proteobacteria; Gammaproteobacteria; Enterobacterales; Enterobacteriaceae; Escherichia | Enterobacterales | Animal associated  (duck feces) | NCBI entry |
| [*Gramella jeungdoensis*](https://lpsn.dsmz.de/species/gramella-jeungdoensis) strain KCTC 23123 | Bacteria; Bacteroidetes; Flavobacteriia; Flavobacteriales; Flavobacteriaceae; Gramella | Flavobacteriales | Solar saltern | [Joung et al. 2011](https://pubmed.ncbi.nlm.nih.gov/22203568/) |
| [*Kosakonia cowanii*](https://lpsn.dsmz.de/species/kosakonia-cowanii) strain Esp_Z chromosome | Bacteria; Proteobacteria; Gammaproteobacteria; Enterobacterales; Enterobacteriaceae; Kosakonia | Enterobacterales | human specimens | [Inoue et al. 2000](https://pubmed.ncbi.nlm.nih.gov/11080391/) |
| *Labrys* sp. WJW | Bacteria; Proteobacteria; Alphaproteobacteria; Hyphomicrobiales; Xanthobacteraceae; Labrys | Hyphomicrobiales | soil | [NCBI entry](https://www.ncbi.nlm.nih.gov/nuccore/NZ_LYXZ00000000.1) |
| *Listeria monocytogenes* strain C5 | Bacteria; Firmicutes; Bacilli; Bacillales; Listeriaceae; Listeria | Bacillales | Freshwater | [NCBI entry](https://www.ncbi.nlm.nih.gov/nuccore/NZ_MDQI00000000.1) |
| [*Listeria monocytogenes*](https://lpsn.dsmz.de/species/listeria-monocytogenes) strain 2014L-6088 | Bacteria; Firmicutes; Bacilli; Bacillales; Listeriaceae; Listeria | Bacillales | Animal associated (rabbit) | [ATCC](https://www.lgcstandards-atcc.org/Products/All/15313) |
| [*Mammaliicoccus sciuri*](https://lpsn.dsmz.de/species/mammaliicoccus-sciuri) strain GDM7P051A C-1_NDMS04239_1 | Bacteria; Firmicutes; Bacilli; Bacillales; Staphylococcaceae; Mammaliicoccus | Bacillales | Animal associated (squirrel) | [Kloos et al. 1976](https://www.microbiologyresearch.org/content/journal/ijsem/10.1099/00207713-26-1-22) |
| [*Maritimibacter*](https://lpsn.dsmz.de/genus/maritimibacter) sp. LZ-17 NODE_6 | Bacteria; Proteobacteria; Alphaproteobacteria; Rhodobacterales; Rhodobacteraceae; Maritimibacter; unclassified Maritimibacter | Rhodobacterales | Plant associated (dinoflagellate) | [NCBI entry](https://www.ncbi.nlm.nih.gov/nuccore/NZ_SWKQ01000016) |
| [*Mesorhizobium sp.*](https://lpsn.dsmz.de/genus/mesorhizobium) 3P27G6 | Bacteria; Proteobacteria; Alphaproteobacteria; Hyphomicrobiales; Phyllobacteriaceae; Mesorhizobium | Hyphomicrobiales | alpine spring water | [NCBI entry](https://www.ncbi.nlm.nih.gov/nuccore/NZ_SCNN00000000.1) |
| [*Mesorhizobium composti*](https://lpsn.dsmz.de/species/mesorhizobium-composti) strain CC-YTH430 | Bacteria; Proteobacteria; Alphaproteobacteria; Hyphomicrobiales; Phyllobacteriaceae; Mesorhizobium | Hyphomicrobiales | from a compost sample in Taiwan | [Lin et al. 2019](https://pubmed.ncbi.nlm.nih.gov/31055717/) |
| [*Mesorhizobium sp.*](https://lpsn.dsmz.de/genus/mesorhizobium) M7A.F.Ca.ET.027.02.1.1 NODE_4 | Bacteria; Proteobacteria; Alphaproteobacteria; Hyphomicrobiales; Phyllobacteriaceae; Mesorhizobium | Hyphomicrobiales | Plant root nodule (*Cicer arietinum*) | [NCBI entry](https://www.ncbi.nlm.nih.gov/nuccore/NZ_RZOL01000004) |
| [*Mesorhizobium sp.*](https://lpsn.dsmz.de/genus/mesorhizobium) M1A.F.Ca.IN.020.30.1.1 NODE_39 | Bacteria; Proteobacteria; Alphaproteobacteria; Hyphomicrobiales; Phyllobacteriaceae; Mesorhizobium | Hyphomicrobiales | Plant root nodule (*Cicer arietinum*) | [NCBI entry](https://www.ncbi.nlm.nih.gov/nuccore/NZ_RZSO01000039) |
| [*Mesorhizobium sp.*](https://lpsn.dsmz.de/genus/mesorhizobium) M1A.F.Ca.ET.072.01.1.1 NODE_40 | Bacteria; Proteobacteria; Alphaproteobacteria; Hyphomicrobiales; Phyllobacteriaceae; Mesorhizobium | Hyphomicrobiales | Plant root nodule (*Cicer arietinum*) | [NCBI entry](https://www.ncbi.nlm.nih.gov/nuccore/NZ_RZSF01000040) |
| [*Mesorhizobium sp.*](https://lpsn.dsmz.de/genus/mesorhizobium) M1A.F.Ca.IN.022.06.1.1 | Bacteria; Proteobacteria; Alphaproteobacteria; Hyphomicrobiales; Phyllobacteriaceae; Mesorhizobium | Hyphomicrobiales | Plant root nodule (*Cicer arietinum*) | [NCBI entry](https://www.ncbi.nlm.nih.gov/nuccore/CP034455) |
| [*Mesorhizobium sp.*](https://lpsn.dsmz.de/genus/mesorhizobium) isolate N.Cr.TU.016.05.1 NODE_86 | Bacteria; Proteobacteria; Alphaproteobacteria; Hyphomicrobiales; Phyllobacteriaceae; Mesorhizobium | Hyphomicrobiales | Plant root nodule  (*Cicer arietinum*) | [NCBI entry](https://www.ncbi.nlm.nih.gov/nuccore/SASU01000086) |
| [*Mesorhizobium sp.*](https://lpsn.dsmz.de/genus/mesorhizobium) M5C.F.Ca.IN.020.29.1.1 NODE_361 | Bacteria; Proteobacteria; Alphaproteobacteria; Hyphomicrobiales; Phyllobacteriaceae; Mesorhizobium | Hyphomicrobiales | Plant root nodule  (*Cicer arietinum*) | [NCBI entry](https://www.ncbi.nlm.nih.gov/nuccore/NZ_RZSW01000361) |
| [*Neorhizobium*](https://lpsn.dsmz.de/species/neorhizobium-tomejilense) *sp.* T17_20 | Bacteria; Proteobacteria; Alphaproteobacteria; Hyphomicrobiales; Rhizobiaceae; Rhizobium/Agrobacterium group; Neorhizobium. | Hyphomicrobiales | Terrestrial, soil | [NCBI entry](https://www.ncbi.nlm.nih.gov/nuccore/PVBG01000001.1) |
| [*Nioella nitratireducens*](https://lpsn.dsmz.de/species/nioella-nitratireducens) strain SSW136 | Bacteria; Proteobacteria; Alphaproteobacteria; Rhodobacterales; Rhodobacteraceae; Nioella | Rhodobacterales | surface sediment of the Jiulong River transered into estuary water | [Liu et al. 2017](https://www.microbiologyresearch.org/content/journal/ijsem/10.1099/ijsem.0.001798) |
| [*Nitrospira*](https://lpsn.dsmz.de/genus/nitrospira) sp. isolate RSF5 | Bacteria; Nitrospirae; Nitrospirales; Nitrospiraceae; Nitrospira | Nitrospirales | rapid sand filter | [NCBI entry](https://www.ncbi.nlm.nih.gov/nuccore/SWDO01000020) |
| [*Novosphingobium tardaugens*](https://lpsn.dsmz.de/species/novosphingobium-tardaugens) NBRC 16725  New name: *Caenibius tardaugens* | Bacteria; Proteobacteria; Alphaproteobacteria; Hyphomicrobiales; Hyphomicrobiaceae; Novosphingobium | Rhodobacterales | Marine | [Zheng et al. 2018](https://www.frontiersin.org/articles/10.3389/fmicb.2018.01418/full) |
| [*Oceanicola*](https://lpsn.dsmz.de/genus/oceanicola) *sp.* S124 | Bacteria; Proteobacteria; Alphaproteobacteria; Rhodobacterales; Rhodobacteraceae; Oceanicola. | Rhodobacterales | Marine | [Kwon et al. 2012](https://journals.asm.org/doi/10.1128/JB.01614-12) |
| [*Oscillibacter*](https://lpsn.dsmz.de/genus/oscillibacter) sp. PC13 | Bacteria; Firmicutes; Clostridia; Clostridiales; Oscillospiraceae; Oscillibacter; unclassified Oscillibacter | Clostridiales | isolated from sheep rumen | [Seshadri et al. 2018](https://www.nature.com/articles/nbt.4110.pdf) |
| [*Paenibacillus odorifer*](https://lpsn.dsmz.de/species/paenibacillus-odorifer) strain DSM 15391 chromosome | Bacteria; Firmicutes; Bacilli; Bacillales; Paenibacillaceae; Paenibacillus | Bacillales | Soil from wheat roots | [NCBI entry](https://www.ncbi.nlm.nih.gov/nuccore/NZ_CP009428)  [Berge et al. 2002](https://pubmed.ncbi.nlm.nih.gov/11931174/) |
| [*Parabacteroides distasonis*](https://lpsn.dsmz.de/species/parabacteroides-distasonis) str. 3999B T(B) 6 | Bacteria; Bacteroidetes; Bacteroidia; Bacteroidales; Tannerellaceae; Parabacteroides | Bacteroidales | human feces | [Sakamoto et al. 2006](https://pubmed.ncbi.nlm.nih.gov/16825636/) |
| [*Parabacteroides*](https://lpsn.dsmz.de/genus/parabacteroides) sp. AF39-10AC | Bacteria; Bacteroidetes; Bacteroidia; Bacteroidales; Tannerellaceae; Parabacteroides | Bacteroidales | Human feces | [NCBI entry](https://www.ncbi.nlm.nih.gov/nuccore/NZ_QTLP00000000.1) |
| [*Parabacteroides*](https://lpsn.dsmz.de/genus/parabacteroides) sp. AM27-42 | Bacteria; Bacteroidetes; Bacteroidia; Bacteroidales; Tannerellaceae; Parabacteroides | Bacteroidales | Human feces | [NCBI entry](https://www.ncbi.nlm.nih.gov/nuccore/QTNL01000001.1) |
| [*Paracoccus alkenifer*](https://lpsn.dsmz.de/species/paracoccus-alkenifer) strain DSM 11593 | Bacteria; Proteobacteria; Alphaproteobacteria; Rhodobacterales; Rhodobacteraceae; Paracoccus. | Rhodobacterales | Terestrial, biofilter for waste gas treatment | [Lipski et al. 1998](https://pubmed.ncbi.nlm.nih.gov/9731294/) |
| [*Phaeobacter inhibens*](https://lpsn.dsmz.de/species/phaeobacter-inhibens) strain DOK1-1 | Bacteria; Proteobacteria; Alphaproteobacteria; Rhodobacterales; Rhodobacteraceae; Phaeobacter. | Rhodobacterales | Marine, seawater | [NCBI entry](https://www.ncbi.nlm.nih.gov/nuccore/CP019307.1) |
| [*Phascolarctobacterium faecium*](https://lpsn.dsmz.de/species/phascolarctobacterium-faecium) DSM 14760 | Bacteria; Firmicutes; Negativicutes; Acidaminococcales; Acidaminococcaceae; Phascolarctobacterium | Acidaminococcales | first isolated from the faeces of koala | [Del Dot et al. 1993](https://www.sciencedirect.com/science/article/abs/pii/S0723202011802699) |
| [*Prevotella bergensis*](https://lpsn.dsmz.de/species/prevotella-bergensis)  DSM 17361 | Bacteria; Bacteroidetes; Bacteroidia; Bacteroidales; Prevotellaceae; Prevotella | Bacteroidales | Human skin | [Downes et al 2006](https://pubmed.ncbi.nlm.nih.gov/16514036/) |
| [*Prevotella buccalis*](https://lpsn.dsmz.de/species/prevotella-buccalis) ATCC 35310 | Bacteria; Bacteroidetes; Bacteroidia; Bacteroidales; Prevotellaceae; Prevotella | Bacteroidales | human vaginal cavity | [NCBI entry](https://www.ncbi.nlm.nih.gov/nuccore/NZ_ADEG01000016) |
| [*Prevotella sp.*](https://lpsn.dsmz.de/genus/prevotella) CAG:1185 | Bacteria; Bacteroidetes; Bacteroidia; Bacteroidales; Prevotellaceae; Prevotella | Bacteroidales | human gut | [NCBI entry](https://www.ncbi.nlm.nih.gov/nuccore/HF992559) |
| [*Prevotella salivae*](https://lpsn.dsmz.de/species/prevotella-salivae) F0493 contig0002 | Bacteria; Bacteroidetes; Bacteroidia; Bacteroidales; Prevotellaceae; Prevotella | Bacteroidales | Human oral cavity | [Sakamoto et al. 2004](https://pubmed.ncbi.nlm.nih.gov/15143039/) |
| [*Ralstonia solanacearum*](https://lpsn.dsmz.de/species/ralstonia-solanacearum) strain UW179 | Bacteria; Proteobacteria; Betaproteobacteria; Burkholderiales; Burkholderiaceae; Ralstonia | Burkholderiales | Plant associated | [DSMZ entry](https://lpsn.dsmz.de/species/ralstonia-solanacearum) |
| [*Rhizobium pusense*](https://lpsn.dsmz.de/species/rhizobium-pusense) strain CCGM10 | Bacteria; Proteobacteria; Alphaproteobacteria; Hyphomicrobiales; Rhizobiaceae; Rhizobium/Agrobacterium group; Rhizobium. | Hyphomicrobiales | Terestrial, root nodules  (*Phaseolus vulgaris*) | [Aguilar et al. 2016](https://www.frontiersin.org/articles/10.3389/fmicb.2016.01720/full) |
| [*Rhizobium*](https://lpsn.dsmz.de/genus/rhizobium) sp. BG4 plasmid pRPC1 | Bacteria; Proteobacteria; Alphaproteobacteria; Hyphomicrobiales; Rhizobiaceae; Rhizobium/Agrobacterium group; Rhizobium. | Hyphomicrobiales | Root nodules (*Prosopis cineraria*) | [NCBI entry](https://www.ncbi.nlm.nih.gov/nuccore/CP044126.1) |
| [*Rhizobium*](https://lpsn.dsmz.de/genus/rhizobium) *sp.* MHM7A NODE_12 | Bacteria; Proteobacteria; Alphaproteobacteria; Hyphomicrobiales; Rhizobiaceae; Rhizobium/Agrobacterium group; Rhizobium. | Hyphomicrobiales | Pea root nodules  (*Pisum sativum*) | [NCBI entry](https://www.ncbi.nlm.nih.gov/nuccore/NZ_VCHT01000012) |
| [*Rhizobium*](https://lpsn.dsmz.de/genus/rhizobium) sp. Root564 | Bacteria; Proteobacteria; Alphaproteobacteria; Hyphomicrobiales; Rhizobiaceae; Rhizobium/Agrobacterium group; Rhizobium. | Hyphomicrobiales | Terestrial, root nodules  (*Arabidopsis thaliana*) | [Bai et al. 2015](https://pubmed.ncbi.nlm.nih.gov/26633631/) |
| [*Rhizobium*](https://lpsn.dsmz.de/genus/rhizobium) sp. Root651 | Bacteria; Proteobacteria; Alphaproteobacteria; Hyphomicrobiales; Rhizobiaceae; Rhizobium/Agrobacterium group; Rhizobium. | Hyphomicrobiales | Terestrial, root nodules (*Arabidopsis thaliana*) | [Bai et al. 2015](https://pubmed.ncbi.nlm.nih.gov/26633631/) |
| [*Rhizobium*](https://lpsn.dsmz.de/genus/rhizobium) sp. UR51a | Bacteria; Proteobacteria; Alphaproteobacteria; Hyphomicrobiales; Rhizobiaceae; Rhizobium/Agrobacterium group; Rhizobium. | Hyphomicrobiales | Terestrial, root nodules (*Oryza sativa*) | [Souza et al. 2015](https://pubmed.ncbi.nlm.nih.gov/25838497/) |
| [*Rhizobium*](https://lpsn.dsmz.de/genus/rhizobium) *sp. YS-1r* | Bacteria; Proteobacteria; Alphaproteobacteria; Hyphomicrobiales; Rhizobiaceae; Rhizobium/Agrobacterium group; Rhizobium. | Hyphomicrobiales | Terestrial, decaying wood from thermal pond | [Jackson et al. 2017](https://sfamjournals.onlinelibrary.wiley.com/doi/epdf/10.1111/jam.13401) |
| [*Rhizobium straminoryzae*](https://lpsn.dsmz.de/species/rhizobium-straminoryzae) strain SM12 | Bacteria; Proteobacteria; Alphaproteobacteria; Hyphomicrobiales; Rhizobiaceae; Rhizobium/Agrobacterium group; Rhizobium | Hyphomicrobiales | Plant associated Rhizosphere (*Oryza sativa*) | [NCBI entry](https://www.ncbi.nlm.nih.gov/nuccore/NZ_VJMG01000008) |
| [*Rhodobacter capsulatus*](https://lpsn.dsmz.de/species/rhodobacter-capsulatus) YW2 | Bacteria; Proteobacteria; Alphaproteobacteria; Rhodobacterales; Rhodobacteraceae; Rhodobacter. | Rhodobacterales | Terrestrial, forest | [Weaver et al. 1975](https://link.springer.com/article/10.1007%2FBF00447139) |
| [*Rhodobacter capsulatus*](https://lpsn.dsmz.de/species/rhodobacter-capsulatus) R121 | Bacteria; Proteobacteria; Alphaproteobacteria; Rhodobacterales; Rhodobacteraceae; Rhodobacter. | Rhodobacterales | Pond water | [Ding et al. 2014](https://www.ncbi.nlm.nih.gov/pmc/articles/PMC3924369/)  [ATCC](https://www.atcc.org/products/33303) |
| [*Rhodobacter*](https://lpsn.dsmz.de/genus/rhodobacter) sp. SW2 | Bacteria; Proteobacteria; Alphaproteobacteria; Rhodobacterales; Rhodobacteraceae; Rhodobacter. | Rhodobacterales | Terrestrial, freshwater mud samples | [Ehrenreich et al. 1994](https://pubmed.ncbi.nlm.nih.gov/7811087/) |
| [*Rhodovulum*](https://lpsn.dsmz.de/genus/rhodovulum) *sp.* MB263 | Bacteria; Proteobacteria; Alphaproteobacteria; Rhodobacterales; Rhodobacteraceae; Rhodovulum. | Rhodobacterales | Marine, coastal mud flat | [Hiraishi and Ueda 1995 et al.](https://pubmed.ncbi.nlm.nih.gov/7537066/) |
| [*Rhodovulum sulfidophilum*](https://lpsn.dsmz.de/species/rhodovulum-sulfidophilum) strain DSM 2351 | Bacteria; Proteobacteria; Alphaproteobacteria; Rhodobacterales; Rhodobacteraceae; Rhodovulum. | Rhodobacterales | Marine, mud from intertidal flats | [Hansen and Veldkamp et al. 1973](https://link.springer.com/article/10.1007%2FBF00409510) |
| *[Roseovarius](https://lpsn.dsmz.de/genus/roseovarius)* sp. GCL-8 | Bacteria; Proteobacteria; Alphaproteobacteria; Rhodobacterales; Rhodobacteraceae; Roseovarius | Rhodobacterales | sand | [NCBI entry](https://www.ncbi.nlm.nih.gov/nuccore/NZ_QITQ00000000.1) |
| [*Ruegeria mobilis*](https://lpsn.dsmz.de/species/ruegeria-mobilis) (different strains)  New name: *Tritonibacter mobilis* | Bacteria; Proteobacteria; Alphaproteobacteria; Rhodobacterales; Rhodobacteraceae; Ruegeria. | Rhodobacterales | Marine, seawater, sediment and copepod, | [Sonnenschein et al. 2017](https://www.nature.com/articles/ismej2016111) |
| [*Ruegeria sp. THAF57*](https://lpsn.dsmz.de/genus/ruegeria) | Bacteria; Proteobacteria; Alphaproteobacteria; Rhodobacterales; Rhodobacteraceae; Ruegeria. | Rhodobacterales | marine | [NCBI entry](https://www.ncbi.nlm.nih.gov/nuccore/NZ_CAIWNQ010000009) |
| [*Stenotrophomonas maltophilia*](https://lpsn.dsmz.de/species/stenotrophomonas-maltophilia) strain AS012690 | Bacteria; Proteobacteria; Gammaproteobacteria; Xanthomonadales; Xanthomonadaceae; Stenotrophomonas; Stenotrophomonas maltophilia group | Xanthomonadales | human specimens | [NCBI entry](https://www.ncbi.nlm.nih.gov/nuccore/NZ_VLGL01000002) |
| [*Stenotrophomonas rhizophila*](https://lpsn.dsmz.de/species/stenotrophomonas-rhizophila) strain JC1 | Bacteria; Proteobacteria; Gammaproteobacteria; Xanthomonadales; Xanthomonadaceae; Stenotrophomonas | Xanthomonadales | Rhizosphere of potato and rape | [Wolf et al. 2002](https://pubmed.ncbi.nlm.nih.gov/12508851/) |
| [*Xanthomonas citri*](https://lpsn.dsmz.de/species/xanthomonas-citri) strain LE3-1 | Bacteria; Proteobacteria; Gammaproteobacteria; Xanthomonadales; Xanthomonadaceae; Xanthomonas | Xanthomonadales | Plant associated  (C. aurantifolia) | [NCBI entry](https://www.ncbi.nlm.nih.gov/nuccore/NZ_LN647176) |

Table 3: Predicted prophages and their categories

| **Name** | **DNA type** | **Size (bps)** | **Contig size (bps)** | **True prophage** | **Potentially episomes** | **Not Microviridae** | **Sequencing contaminants** |
| --- | --- | --- | --- | --- | --- | --- | --- |
| *Acinetobacter lwoffii* strain SU1904 | Contig | 5.673 | 5.673 |  | X |  |  |
| *Agrobacterium larrymoorei* strain CFBP5477 | ND* | 6.301 | 657.589 | X |  |  |  |
| *Agrobacterium tumefaciens* strain 1D1108 plasmid pAt1D1108a ┼ | plasmid | 5.860 | 502.2­74 | X |  |  |  |
| *Alistipes sp.* isolate P1-1 tig00001311 | ND* | 5.699 | 3.006.009 | X |  |  |  |
| *Alistipes onderdonkii* WAL 8169 = DSM 19147 | ND* | 5.793 | 100.104 | X |  |  |  |
| *Aphanizomenon* flos aquae WA102 3645 | Contig | 4.055 | - |  | X |  |  |
| *Bacillus altitudinis strain DSM 26896* | whole genome shotgun sequence | 13.312 | 156.970 |  |  | X |  |
| *Bacillus sp. X1(2014) strain DE0237 NODE_39* | whole genome shotgun sequence | 7.309 | 33.618 |  |  | X |  |
| *Bacillus thuringiensis serovar T01001* | Circular chromosome | 11.378 | 6.323.123 |  |  | X |  |
| *Bacillus wiedmannii strain FSL J3-0113 NODE_13* | whole genome shotgun sequence | 9.571 | 109.662 |  |  | X |  |
| *Bacteroides caccae* strain 2789STDY5834946 | ND* | 5.571 | 221.420 | X |  |  |  |
| *Bacteroides caccae strain* AF46-5GN AF46-5GN.Scaf95 | ND* | 5.370 | 19.734 | X |  |  |  |
| *Bacteroides eggerthii* DSM 20697 Scfld2 ┼ | ND* | 6.195 | 19.345 | X |  |  |  |
| *Bacteroides finegoldii strain D54t1_190329_G10 NODE_5* | whole genome shotgun sequence | 6.471 | 317.103 | X |  |  |  |
| *Bacteroides ovatus* isolate Bacteroides_ovatus_MC1 | ND* | 6.402 | 59.930 | X |  |  |  |
| *Bacteroides plebeius* DSM 17135 | whole genome shotgun sequence | 5.059 | 569.841 | X |  |  |  |
| *Bacteroides sp. 2_2_4* supercont1.3 ┼ | ND* | 6.303 | 605.506 | X |  |  |  |
| *Bacteroides sp.* AF29-11 AF29-11.Scaf2 | ND* | 6.481 | 622.059 | X |  |  |  |
| *Bacteroides thetaiotaomicron strain* 19 ┼ | ND* | 6.056 | 17.932 | X |  |  |  |
| *Bacteroides xylanisolvens* strain AF38-2 AF38-2.Scaf17 | ND* | 8.224 | 141.458 | X |  |  |  |
| Candidatus *Rhodobacter lobularis* isolate IGS | ND* | 7.740 | 3.995.303 | X |  |  |  |
| *Clostridioides difficile CD129* | Contig | 5.697 | 5.697 |  |  |  | X |
| *Coprobacter fastidiosus isolate CIM:MAG 335* | whole genome shotgun sequence | 6.609 | 20.190 | X |  |  |  |
| *Devosia chinhatensis* strain IPL18 | ND* | 7.499 | 2.139.066 | X |  |  |  |
| *Devosia* sp. YR412 | ND* | 11.405 | 221.727 | X |  |  |  |
| *Dysgonomonas macrotermitis* strain DSM 27370 | ND* | 6.139 | 265.550 | X |  |  |  |
| *Dysgonomonas* sp. 521 scaffold10_size73398 | ND* | 6.816 | 73.398 | X |  |  |  |
| *Elizabethkingia anophelis* strain E6809 2b | ND* | 4.392 | 1.150.786 | X |  |  |  |
| *Epibacterium ulvae* strain U95 | ND* | 7.870 | 804.761 | X |  |  |  |
| *Erysipelatoclostridium* sp. An15 An15 | ND* | 6.491 | 201.071 | X |  |  |  |
| *Escherichia coli strain OLC1558 OLC1558* ┼ | whole genome shotgun sequence | 4.477 | 351.548 | X |  |  |  |
| *Escherichia sp.* MOD1-EC6163 | ND* | 5.993 | 50.742 | X |  |  |  |
| *Gramella jeungdoensis strain KCTC 23123 KCTC_32123* | Contig | 6.302 | 6.302 |  | X |  |  |
| *Kosakonia cowanii strain Esp_Z* | Circular chromosome | 5.002 | 5.077.975 | X |  |  |  |
| *Labrys sp. WJW* | ND* | 9.368 | 94.779 | X |  |  |  |
| *Listeria monocytogenes strain C5* | whole genome shotgun sequence | 10.033 | 1.240.180 |  |  | X |  |
| *Listeria monocytogenes strain 2014L-6088 NODE_2* | whole genome shotgun sequence | 6.184 | 593.093 | X |  | X |  |
| *Mammaliicoccus sciuri strain GDM7P051A C-1* | Contig | 5.855 | 5.855 |  | X |  |  |
| *Maritimibacter sp. LZ-17 NODE_6* | whole genome shotgun sequence | 8.679 | 114.540 | X |  |  |  |
| *Mesorhizobium sp. 3P27G6 NODE_30* | ND* | 6.025 | 67.438 | X |  |  |  |
| *Mesorhizobium composti strain* CC-YTH430 | ND* | 5.831 | 699.301 | X |  |  |  |
| *Mesorhizobium sp. M7A.F.Ca.ET.027.02.1.1 NODE_4* | ND* | 6.940 | 56.052 | X |  |  |  |
| *Mesorhizobium sp. M1A.F.Ca.IN.020.30.1.1 NODE_39* | ND* | 7.714 | 17.913 | X |  |  |  |
| *Mesorhizobium sp. M1A.F.Ca.ET.072.01.1.1 NODE_40* | ND* | 8.388 | 32.967 | X |  |  |  |
| *Mesorhizobium sp. M1A.F.Ca.IN.022.06.1.1* ┼ | circular chromosome | 7.165 | 6.278.406 | X |  |  |  |
| *Mesorhizobium sp. isolate N.Cr.TU.016.05.1 NODE_86* | ND* | 6.165 | 23.594 | X |  |  |  |
| *Mesorhizobium sp. M5C.F.Ca.IN.020.29.1.1 NODE_361* | Contig | 6.491 | 6.491 |  | X |  |  |
| *Neorhizobium* sp. T17_20 | ND* | 6.427 | 571.600 | X |  |  |  |
| *Nioella nitratireducens* strain SSW136 | ND* | 5.734 | 95.120 | X |  |  |  |
| *Nitrospira sp. isolate RSF5* | whole genome shotgun sequence | 6.250 | 96.140 | X |  |  |  |
| *Novosphingobium tardaugens* NBRC 16725 | ND* | 4.443 | 102.866 | X |  |  |  |
| *Oceanicola* sp. S124 | ND* | 9115 | 38.215 | X |  |  |  |
| *Oscillibacter sp. PC13* | ND* | 5.417 | 177.285 | X |  |  |  |
| *Paenibacillus odorifer strain DSM 15391* ┼ | Circular chromosome | 8.259 | 6.812.473 | X |  |  |  |
| *Parabacteroides distasonis str. 3999B* ┼ | ND* | 4.626 | 85.604 | X |  |  |  |
| *Parabacteroides sp. AF39-10AC* ┼ | ND* | 4.583 | 76.059 | X |  |  |  |
| *Parabacteroides sp. AM27-42* | ND* | 4.983 | 114.974 | X |  |  |  |
| *Paracoccus alkenifer* strain DSM 11593 | ND* | 8.198 | 871.604 | X |  |  |  |
| *Phaeobacter inhibens* strain DOK1-1 | Chromosome, circular | 5.922 | 3.718.082 | X |  |  |  |
| *Phascolarctobacterium faecium DSM 14760* | ND* | 4.517 | 796.791 | X |  |  |  |
| *Prevotella bergensis* DSM 17361 ┼ | whole genome shotgun sequence | 5.337 | 1.108.685 | X |  |  |  |
| *Prevotella buccalis* ATCC 35310 ┼ | ND* | 4.975 | 88.363 | X |  |  |  |
| *Prevotella sp. CAG:1185* ┼ | whole genome shotgun sequence | 6.349 | 34.922 | X |  |  |  |
| *Prevotella salivae F0493* | Contig | 6.888 | 6.888 |  | X |  |  |
| *Ralstonia solanacearum strain UW179* | whole genome shotgun sequence | 6.503 | 5.426.414 |  |  | X |  |
| *Rhizobium pusense* strain CCGM10 | ND* | 8.944 | 486.269 | X |  |  |  |
| *Rhizobium sp. BG4 plasmid pRPC1* | Circular plasmid | 6.609 | 1.609.137 | X |  |  |  |
| *Rhizobium sp. MHM7A NODE_12* ┼ | whole genome shotgun sequence | 6.733 | 221.567 | X |  |  |  |
| *Rhizobium* sp. Root564 | ND* | 10.526 | 1.029.951 | X |  |  |  |
| *Rhizobium* sp. Root651 | ND* | 8.427 | 167.271 | X |  |  |  |
| *Rhizobium* sp. UR51a | ND* | 5.942 | 277.936 | X |  |  |  |
| *Rhizobium* sp. YS-1r | ND* | 8.570 | 657.089 | X |  |  |  |
| *Rhizobium straminoryzae strain SM12* | whole genome shotgun sequence | 7.361 | 175.733 | X |  |  |  |
| *Rhodobacter capsulatus* YW2 | ND* | 6.427 | 316.953 | X |  |  |  |
| *Rhodobacter capsulatus* R121 | ND* | 6.611 | 418.689 | X |  |  |  |
| *Rhodobacter* sp. SW2 | ND* | 9.745 | 188.067 | X |  |  |  |
| *Rhodovulum sulfidophilum* strain DSM 2351 | Plasmid , circular | 6.209 | 111.306 | X |  |  |  |
| *Rhodovulum* sp. MB263 | Chromosome, circular | 6.583 | 3.860.570 | X |  |  |  |
| *Roseovarius* sp. GCL-8 | ND* | 6.624 | 282.961 | X |  |  |  |
| *Ruegeria sp. THAF57* | whole genome shotgun sequence | 13.152 | 210.182 | X |  |  |  |
| *Ruegeria mobilis* strain S1942 | ND | 5.802 | 603.960 | X |  |  |  |
| *Stenotrophomonas maltophilia strain AS012690* | whole genome shotgun sequence | 11.045 | 130.645 |  |  | X |  |
| *Stenotrophomonas rhizophila strain JC1* | circular chromosome | 6.961 | 4.268.161 |  |  | X |  |
| *Xanthomonas citri pv. citri strain LE3-1* | whole genome shotgun sequence | 6.748 | 5.045.425 |  |  | X |  |

*ND – not determined

┼ no 16S rRNA sequences available
