## Supplementary material for "New Microviridae isolated from Sulfitobacter reveals two cosmopolitan subfamilies of ssDNA phages infecting marine and terrestrial Alphaproteobacteria": SI_file_5

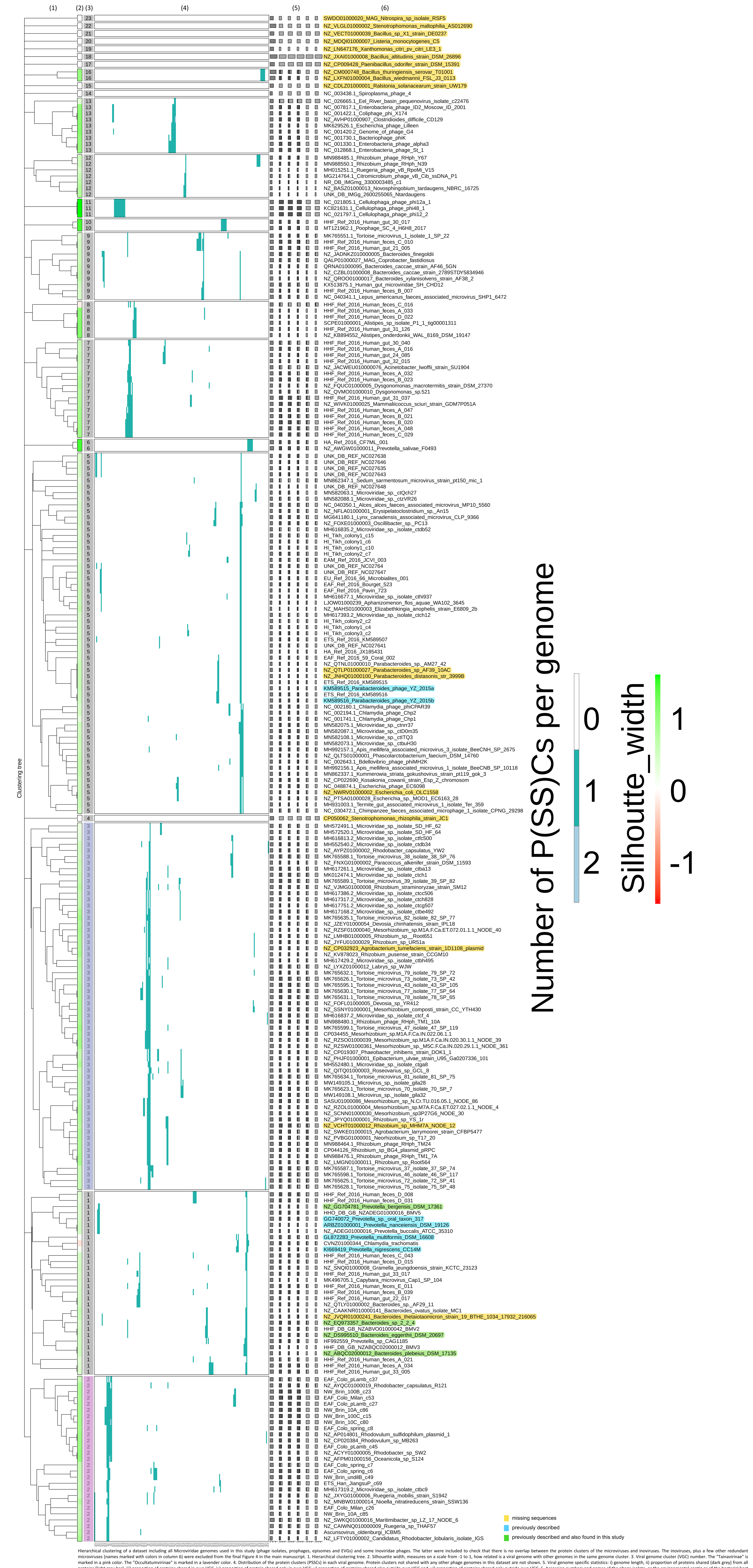


Hierarchical clustering of a dataset including all Microviridae genomes used in this study (phage isolates, prophages, episomes and EVGs) and some Inoviridae phages. The latter were included to check that there is no overlap between the protein clusters of the microviruses and inoviruses. The inoviruses, plus a few other redundant microviruses (names marked with colors in column 6) were excluded from the final Figure 4 in the main manuscript. 1. Hierarchical clustering tree. 2. Silhouette width, measures on a scale from -1 to 1, how related is a viral genome with other genomes in the same genome cluster. 3. Viral genome cluster (VGC) number. The "Tainavirinae" is marked in a pink color. The "Occultatumvirinae" is marked in a lavender color. 4. Distribution of the protein clusters (PSSCs) in each viral genome. Protein clusters not shared with any other phage genomes in this dataset are not shown. 5. Viral genome specific statistics: i) genome length, ii) proportion of proteins shared (dark grey) from all proteins (light grey bar), iii) proportion of proteins shared in own VGC, iv) proportion of protein shared only in own VGC, v) proportion of proteins shared also outside own VGC and, vi) proportion of proteins shared only outside own VGC. 6. Accession numbers and names of the phage isolate, or the environmental contig or of the bacterial host in which a (pro)-phage was predicted.
