## Supplementary figures and images for "New Microviridae isolated from Sulfitobacter reveals two cosmopolitan subfamilies of ssDNA phages infecting marine and terrestrial Alphaproteobacteria"

### SI_file_7

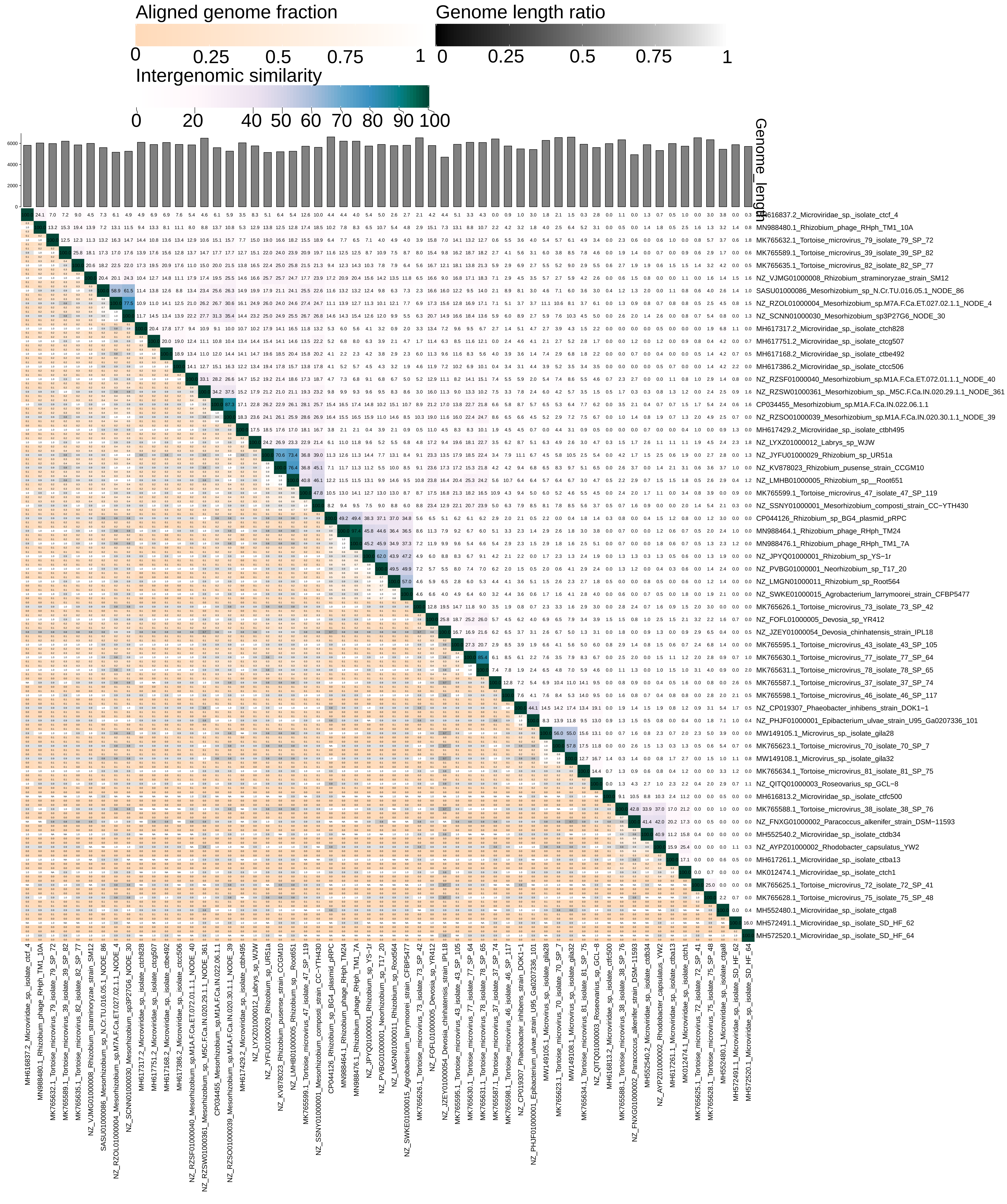

### SI_file_8

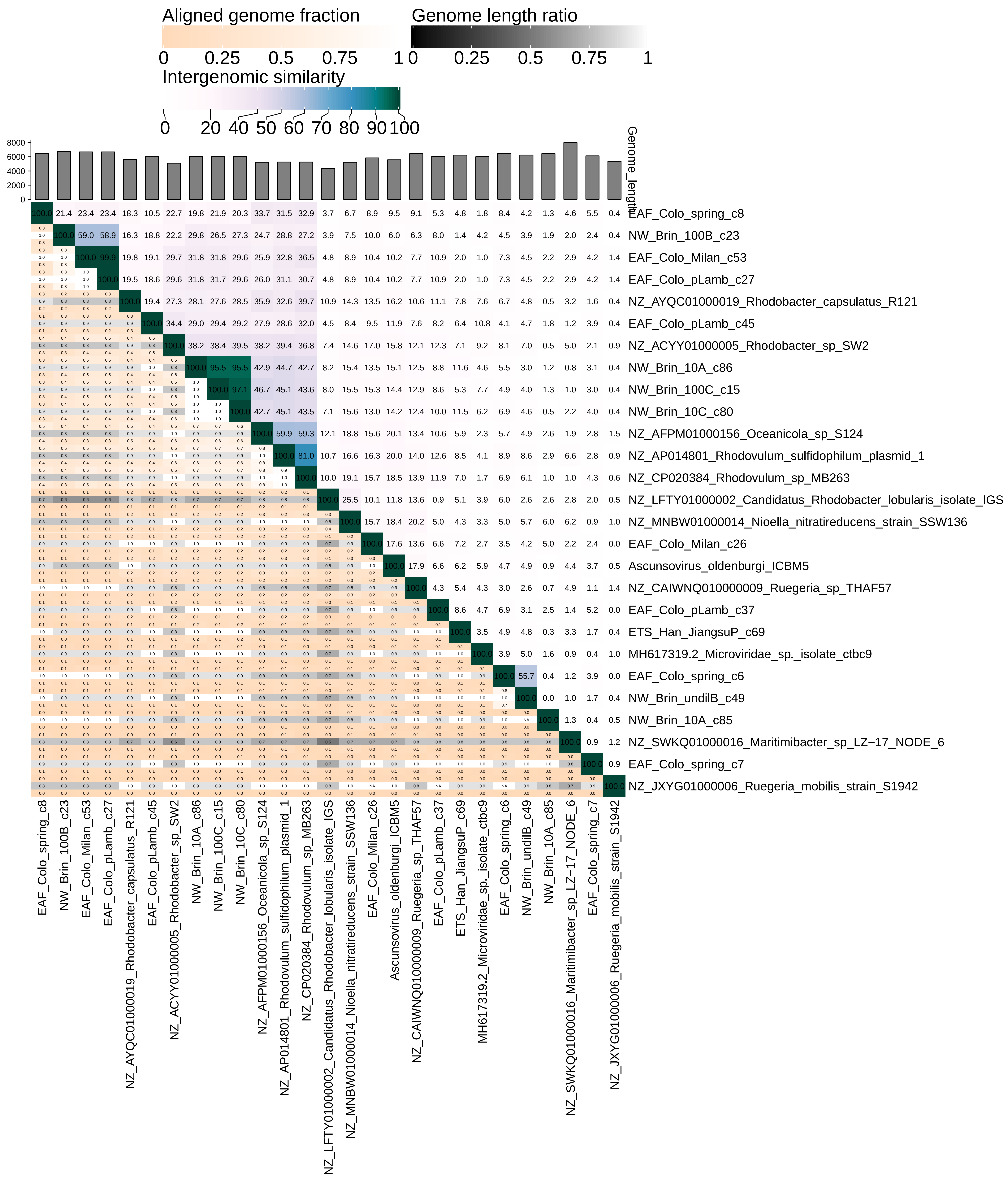
